# The effect of virtual reality modality level of immersion and locomotion on spatial learning and gaze measures

**DOI:** 10.1101/2025.03.09.642230

**Authors:** Michal Gabay, Tom Schonberg

**Affiliations:** School of Neurobiology, Biochemistry and Biophysics, The George S. Wise Faculty of Life Sciences and Sagol School of Neuroscience, Tel Aviv University, Israel

**Keywords:** Spatial learning, Strategy, Virtual reality, Immersion, Proprioception, Eye-tracking

## Abstract

The widespread adoption of head-mounted display (HMD) virtual reality (VR) systems has emerged in various fields, including spatial learning research. This study investigated the effects of VR modality level of immersion, locomotion interface, and proprioception on spatial learning and physiological measures using eye-tracking (ET) in VR. We translated the classic T-maze task from Barnes et al. (1980) to humans for the first time, comparing three VR modalities: 3D HMD VR with physical walking, 3D HMD VR with controller-based movement, and 2D desktop VR. Results revealed that human participants employed a mixture of cue, place, and response strategies when navigating the virtual T-maze, mirroring rodent behavior. In both samples, no significant differences were found between the two HMD VR conditions in learning performance, nor consistent ones in strategy choices. However, 2D desktop navigation was associated with slower initial learning, though this discrepancy diminished in subsequent sessions. These results were supported by spatial presence, immersion, and naturalness reports. Gaze measures showed that participants who physically walked devoted more visual attention to environmental cues compared to controller users. Predictive models for identifying spatial learning strategies based on ET and behavioral measures demonstrated significant accuracy in some models, particularly in the VR walking condition and second session. Our findings enhance the understanding of spatial learning strategies and the effects of VR modality on cognition and gaze behavior. This work demonstrates the potential of integrated ET data and holds implications for early detection and personalized rehabilitation of neurodegenerative conditions related to spatial cognition.

## 1 Introduction

### 1.1 Spatial learning strategies

Spatial navigation is a multifaceted cognitive process involving diverse strategies that allow individuals across all species to orient themselves and locate goals in their environment. Two core spatial learning strategies, place, and response, have been the focus of research since Tolman’s seminal work (Tolman et al. 1946). In that study, Tolman et al. constructed a cross-maze paradigm (a T-maze with two possible opposite starting points, one in training and the other in the test phase) and demonstrated that rodents could navigate using a place strategy, that relies on forming a cognitive map based on environmental spatial cues, or a response strategy, that depends on fixed motor patterns such as always turning left or right. These findings introduced the concept of flexible navigation, challenging behaviorist paradigms that emphasized stimulus-response (S-R) associations (Goodman 2021).

These strategies are also frequently referred to as allocentric and egocentric navigation. Allocentric navigation (place strategy) involves creating a representation of the environment from an external perspective, anchored to spatial landmarks or geometric relationships independent of the individual’s position (O’Keefe and Nadel 1978). In contrast, egocentric navigation (response strategy) relies on a self-centered perspective, encoding routes or turns relative to the individual’s body axis. These complementary systems enable flexible and habitual navigation and are crucial for adapting to dynamic environments (Iaria et al. 2003).

Building on Tolman’s foundation, another study (Barnes et al. 1980) expanded the T-maze paradigm to include a third strategy in addition to place and response, the cue (beacon) strategy, where animals navigate toward a distinct intra-maze landmark, which was a rubber mat in their design. Their experimental design included three types of probe trials, in which each strategy was tested by rotating the starting point in 180^0^, rotating the outer-maze objects in 180^0^, or shifting the intra-maze cue (see illustration in Fig. 1). This innovative study allowed the dissociation of three navigation strategies and the examination of age-related preferences. Older rodents were shown to favor simpler response strategies over hippocampal-dependent place strategies, highlighting the dynamic interaction between these systems.

**Fig. 1.**
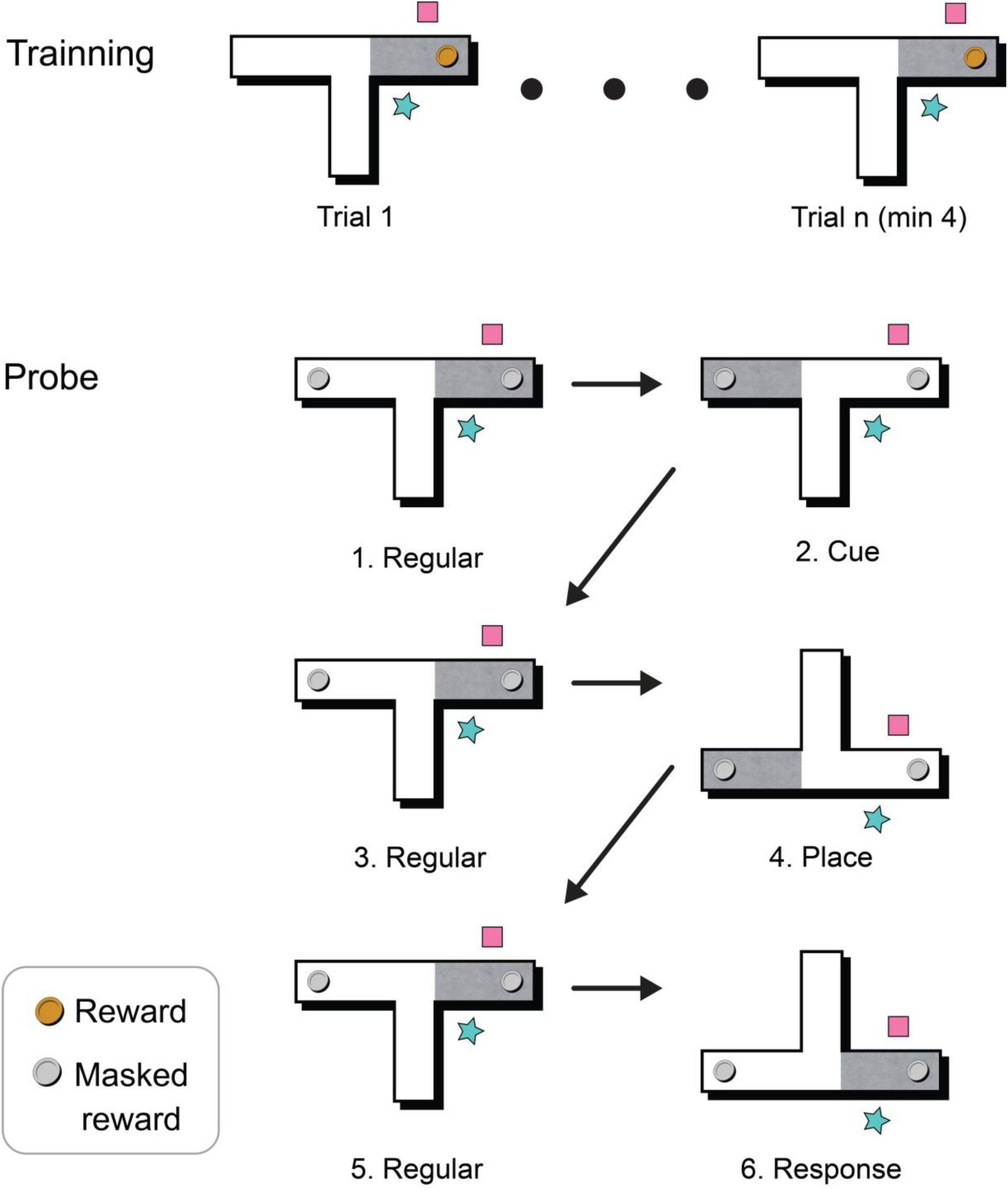
Task design. The test probe trials include cue (2), place (4), and response (6), and the regular probe trials (1,3 and 5) are identical to training. The pink rectangle and cyan star schematically represent two out of the five external cues corresponding to the actual objects shown in Fig. 2

In addition to the T-maze, other paradigms provided additional insights into the place and response spatial learning strategies, where the Morris Water Maze (MWM) (Morris 1981) and the Radial Arm Maze (RAM) (Olton and Samuelson 1976) tasks are most widely used. The RAM consists of multiple arms (8 in the initial version) radiating from a central hub, some of which contain food rewards (Olton and Samuelson 1976). It tests working memory (avoiding re-entry to previously visited arms) and reference memory (learning which arms consistently contain rewards). In the MWM (Morris 1981), rodents were trained to locate a hidden platform submerged in a circular pool of opaque water. In the original version of the MWM, rats were required to use distal environmental cues for navigation (place-based). Variations of the task allowed animals to use cue (beacon) strategy and also test for using response strategy when altering the starting point (Morris 1984).

### 1.2 Translation to humans

Due to the wealth of knowledge derived from animal studies, translating these paradigms to humans has been a significant area of research in cognitive neuroscience. To name a few, the version of the two possible strategies of the T-maze (Levy et al. 2005; Astur et al. 2016; Lin et al. 2022; Gammeri et al. 2022), RAM (Moffat et al. 1998; Iaria et al. 2003; Astur et al. 2004, 2016; Bohbot et al. 2007; Etchamendy et al. 2012; Konishi et al. 2013; Mennenga et al. 2014; Robaey et al. 2016; Kim et al. 2018; Ben-Zeev et al. 2020; Palombi et al. 2022), and MWM (Astur et al. 2004; Levy et al. 2005; Schoenfeld et al. 2017; Ben-Zeev et al. 2020), and even some of their combinations (Astur et al. 2004; Levy et al. 2005; Ben-Zeev et al. 2020) have all been adapted for human studies, with varying degrees of immersion and technological implementation. These tasks focused either on memory (RAM), or one or two possible strategies of place, and response or cue (T-maze, MWM). Some consider the T-maze as the best paradigm to evaluate learning behavior in mice and the simplest way to assess spatial learning strategies (Sharma et al. 2010). However, to the best of our knowledge (Moffat 2009), there are no published studies yet that explore the cue strategy introduced by (Barnes et al. 1980) in addition to place and response, in the T-maze task in humans.

The particular use of one of three spatial learning strategies of place, response, and cue (beacon) was studied in humans in a 2D virtual plus maze (de Condappa and Wiener 2016) and compared between young and old age groups (Wiener et al. 2013). In this paradigm, each plus junction included two landmarks, one on each side. The associative cue strategy involved learning a specific directional response at a landmark, while the beacon strategy entailed moving towards a landmark associated with the goal (the learned turn), and the allocentric strategy involved the creation of a cognitive map of the configuration of the landmarks, which was the only one similar to (Barnes et al. 1980). There are a few key differences between the plus maze and the T-maze paradigms, even though they both examine three strategies. The test trials in the plus maze were approached from different angles, and only using the place strategy led to the correct choice, while in the T-maze, all test trials were approached from the same angle as in training, and the maze setup was altered such that there was no right or wrong choice. In addition, the egocentric strategies in the plus maze were defined by the same landmarks that defined the place strategy which may lead to some ambiguity of the strategies especially in the 180^0^ angle, when it seems that place and beacon strategies may concur, whereas in the T-maze, all the strategies were completely independent of each other. Another study investigated navigating these three strategies in a virtual sunburst maze, which was a replica of Tolman’s classic rodent study (Doner et al. 2023). In the test phase, the original fixed path was blocked, and participants faced 18 radiating hallways (a “sunburst” configuration) to choose an alternative route. It was found that when participants were required to take a shortcut (direct paths to the goal), they consistently employed non-optimal response strategies either following the encoded route or using light as a beacon cue, even when distal boundary cues or enhanced idiothetic cues were added, both in the 2D desktop and the head-mounted displayed virtual reality (HMD VR) versions. In this study as well, there was an optimal route in the test phase that was based on allocentric strategy, differently than in the T-maze.

While these human adaptations of animal paradigms provide valuable insights into spatial learning strategies, it is important to recognize that maze-based laboratory tasks differ substantially from wayfinding in the real world. Everyday navigation encompasses tasks such as search, exploration, route following, and planning, each involving distinct cognitive processes and levels of spatial knowledge (Wiener et al. 2009). Compared to these naturalistic contexts, maze paradigms have reduced ecological validity and limited generalizability. Nevertheless, their constrained design allows the systematic isolation of specific cognitive processes and strategies (cue, place, response) due to experimental control. This complementarity highlights how laboratory-based paradigms can provide theoretical grounding for broader wayfinding research.

### 1.3 Spatial learning in VR

Naturally, the first translations of the spatial learning tasks to humans used a 2D screen display of a virtual environment (VE), and as in other fields of research, in the last decade, the 3D HMD VR has also spread to spatial learning. This approach offers several advantages over traditional 2D or non-immersive paradigms. First, HMD VR provides a more ecologically valid environment for assessing navigation abilities (Vigliocco et al. 2024). For example, the effects of mild aerobic exercise on spatial learning performance in both MWM and RAM were investigated in an HMD VR while navigating with the controller following head movements (Ben-Zeev et al. 2020). In addition, HMD VR allows for full-body movement, as was recently studied in a physically walkable T-maze (Lin et al. 2022). Eventually, it increases the sense of presence that closely mimics real-world navigation (Huffman and Ekstrom 2019). This increased validity may lead to more accurate assessments of spatial cognition and strategy use.

Together with the advantages of advancing from 2D research setups to immersive 3D HMD environments in spatial learning, it is essential to compare these modalities to understand how differences in interaction and immersion affect spatial learning outcomes. Such comparisons are critical for accurately interpreting results across platforms (Huffman and Ekstrom 2019) and for ensuring that findings from 3D HMD studies are grounded in the extensive body of work developed in 2D environments over decays (Hejtmanek et al. 2020). This approach mirrors the importance of bridging findings and paradigms between animal and human research, allowing for deeper insights into spatial cognition across contexts and species (Schoenfeld et al. 2017).

Efforts to investigate how VR modalities influence navigation-related cognitive processes have led to direct comparisons between real-world and VR environments, as well as between different VR systems and interfaces. For instance, the navigation performance in real-world environments was compared to HMD VR setups (Martelli et al. 2019), revealing differences in the use of spatial cues and exit times between these conditions. This study demonstrated that immersive VR could effectively replicate some aspects of real-world navigation while highlighting the impact of visual and environmental fidelity on spatial learning. Similarly, differences in navigation and learning across VR systems with varying levels of immersion were explored (Hejtmanek et al. 2020), including real-world learning, HMD VR with an omnidirectional treadmill, and desktop VR using a mouse and keyboard. They found that physically engaging systems, such as real-world and treadmill-based VR, enhanced spatial memory and cognitive map accuracy compared to static desktop VR. In addition, HMD VR and desktop versions of an 8-arm RAM were compared (Kim et al. 2018), measuring working memory errors, reference memory errors, and other behavioral metrics. The study found that while performance did not significantly differ between the two modalities, HMD VR elicited greater subjective presence and immersion, emphasizing the potential advantages of immersive systems for engaging participants in spatial tasks. Additionally, fully virtual environments reconstructed from real-world spaces were compared to augmented reality (AR) in real-world settings (Mayrose and Maidenbaum 2024), and no significant differences were found in subjective reports as immersion or objective spatial memory performance, suggesting that AR and VR may be used interchangeably in some contexts.

When the role of proprioception, body-based information of movement and rotation, on spatial learning was studied (Ruddle et al. 2011), it was found that proprioceptive feedback significantly improved the accuracy of cognitive maps in VR environments. Notwithstanding, another study suggested that the neural correlates of spatial retrieval tasks with functional magnetic resonance imaging (fMRI) are modality-independent based on the three VR levels of body-based and joystick movement tested (Huffman and Ekstrom 2019). The impact of locomotion interfaces in HMD VR was further examined by comparing controller-based navigation interfaces and assessing their influence on performance and subjective experiences such as presence and immersion (Lim et al. 2022). This study highlighted how interface design could modulate cognitive load and navigation efficiency, with more intuitive interfaces generally yielding better outcomes.

These studies demonstrate substantial research exploring the effects of VR modality and locomotion interfaces on spatial learning, focusing on metrics such as accuracy, immersion, memory, and neural correlates. However, no studies to date have directly tested the influence of VR modality and locomotion interface on spatial learning strategies of cue, place, and response. While strategy usage across two desktop VR samples and one HMD VR sample was recently examined (Doner et al. 2023), the HMD VR sample was significantly smaller than the 2D, and the participants completed all conditions sequentially, unlike the 2D samples, where participants were assigned to a single condition. These studies collectively underscore the need to systematically compare 2D and various 3D VR modalities, levels of immersion as well as different locomotion interfaces, to fully understand their effects on spatial learning, and highlight the need for more controlled comparisons to isolate the effects of VR modality on strategy selection.

### 1.4 Physiological representation of spatial learning

Visual information processing plays a critical role in how we represent and navigate space (Ekstrom 2015), pointing at eye-tracking (ET) as a valuable method for investigating cognitive processes during spatial navigation tasks. ET offers a unique, real-time window into the visual and cognitive processes guiding navigation and has been shown to reveal subtle differences in strategy usage. One study, (Livingstone-Lee et al. 2011) demonstrated that gaze behavior within the first second of a navigation trial could differentiate between allocentric and egocentric strategies in a VR Morris Water Maze (MWM). Another study, (de Condappa and Wiener 2016) examined gaze and pupillometry data to identify navigation strategies in a plus maze, finding that gaze behavior reflected the movement direction during learning and decision-making which was unrelated to the used strategy. They additionally observed that increases in pupil dilation indicated a shift to a more cognitively demanding place strategy. Similarly, gaze dynamics in naturalistic settings have revealed cue preferences (e.g., landmarks versus geometry) that correspond to specific navigation strategies (Bécu et al. 2020), pointing to the utility of visual data for understanding navigation mechanisms. These studies demonstrate the variability in gaze behavior and its relation to strategy across studies, stressing the need for further assessment of this relation.

The integration of ET with VR technology has opened new avenues for spatial navigation research. The viewing behavior during VR navigation was studied using a graph-theoretical analysis approach, revealing complex patterns of visual attention allocation in urban environments (Walter et al. 2022). In addition, another study (Zhu et al. 2022) demonstrated that gaze patterns dynamically adapted to environmental complexity and task demands, balancing attention between environmental transitions and hidden goals during spatial planning in controller-based movement. Likewise, another study (Drewes et al. 2021) compared gaze behavior in VR with controller-based movement to real-world locomotion, highlighting both similarities and differences in visual exploration strategies.

One significant advantage of modern VR headsets is the integration of ET capabilities directly into the device. This built-in functionality eliminates the need for additional equipment, simplifying data collection and improving ecological validity. For example, (Hougaard et al. 2021) utilized the ET capabilities of an HTC Vive headset to study gaze patterns in a virtual museum environment, demonstrating the feasibility of such integrated systems for spatial cognition research.

While ET provides valuable insights, other physiological signals have also been explored in VR spatial navigation studies. A previous study (Lin et al. 2022) combined a physically walkable T-maze with random reward location in VR with EEG recordings and found neural correlates of navigation decisions. Another study, (Do et al. 2021) recorded EEG signals during VR navigation with physical walking, providing insights into brain dynamics during active spatial exploration related to head direction. These studies highlight the potential of combining multiple physiological measures with VR to gain a more comprehensive understanding of spatial cognition. Nevertheless, ET offers unique advantages as a physiological measure in VR navigation studies. Unlike EEG or other techniques requiring additional equipment, ET is seamlessly integrated into many VR headsets, minimizing participant discomfort and maintaining a high degree of ecological validity.

### 1.5 Current research

Considering the growing interest in revealing the effect VR modalities have on cognitive processes, this work aimed to assess how the VR modality level of immersion and locomotion interface affect spatial learning measures, particularly the utilization of spatial learning strategies. For this purpose, we translated for the first time the classic T-maze task form (Barnes et al. 1980) to humans. Therefore, we also aimed to find out whether this elegant task allows the disentanglement of three spatial learning strategies of cue, place, and response for humans as in rodents. We compared three conditions to explore how VR modality and locomotion interfaces influence learning measures in this task. In two of them, the environment was displayed on a 3D HMD VR while physically walking in the first and standing in place and using the controller in the second, and in the third, the environment was displayed on a regular 2D screen using the mouse and keyboard for navigation. In addition, we collected spatial presence and awareness questionnaires, as well as widely used spatial abilities tests to test for their relation to strategy usage.

In addition, in the two HMD VR conditions, we aimed to leverage the advantage of the headset’s built-in ET device by analyzing ET data collected during our translation of the T-maze task for a two pronged-approach. First, we aimed to examine how VR modality and locomotion methods affect gaze behavior. By comparing ET data across different VR conditions (walking vs. controller-based movement), we aimed to study how the level of immersion and physical engagement influence visual exploration and strategy selection in spatial navigation. Second, we aimed to identify the different spatial learning strategies based on physiological (i.e., ET) data. We tested whether gaze features would differentiate strategy selection, as in (Livingstone-Lee et al. 2011) where gaze distinguished allocentric and egocentric navigation, or display mixed association to strategies as in (de Condappa and Wiener 2016) where gaze behavior was unrelated to the used strategy. To achieve this, we developed predictive models to infer navigation strategies from these data alone. Such models could have significant implications for understanding individual differences in spatial cognition and potentially aid in the early detection and rehabilitation of navigation impairments associated with various neurological conditions such as Parkinson’s disease (PD) (Schneider et al. 2017) and Alzheimer’s disease (AD) (Lithfous et al. 2013).

Although the T-maze is a highly simplified paradigm with only one intersection and therefore cannot fully capture the complexity of navigation in real-world environments, especially allocentric navigation, it has nevertheless been successfully used in human studies to dissociate place and response strategies (Astur et al. 2016; Gammeri et al. 2022). Furthermore, it is worth noting that some everyday navigation tasks can be decomposed into a sequence of such single decision points. From this perspective, the T-maze may serve as a useful analog for simple navigational choices encountered in daily life, while preserving experimental control.

Therefore, the specific aims of the present study are as follows:

1. Translate the classical T-maze task from rodents to humans in virtual reality and test whether it elicits the three strategies in humans.
2. Investigate how VR modality and locomotion interface influence spatial learning measures, including strategy use, and subjective experience (e.g., spatial presence, perception, awareness).
3. Examine gaze behavior during spatial navigation, how it is influenced by VR modality, and its relationship to strategy choice.

Although the present study was primarily exploratory, we employed two samples to test data-driven hypotheses. Analyses of the pilot sample were used to inform expectations for the powered sample. Specifically, we hypothesized that effects observed in the pilot would replicate in the powered sample, whereas effects not obtained in the pilot were not expected to emerge.

## 2 Methods

### 2.1 Code and data availability

All the data and the codes for data handling, pre-processing, analysis, and Unity task projects are available in the Open Science Framework (OSF, project page: https://osf.io/arfy9/, preregistration: https://osf.io/4b7kx).

### 2.2 Preregistration

Following data collection on the first sample, we pre-registered the behavioral results and the required sample size for their replication with 80% power with p-value <= 0.05 (see the preregistration section in the OSF project). All analyses that were added following the second sample’s data collection and exploration were mentioned explicitly in the text. The ET analyses were performed following data collection of the two samples. We analyzed the first sample’s results and preregistered the models and hypotheses based on these results prior to observing the second sample’s data. Briefly, all the significant effects that were obtained on the first sample were preregistered, and those that were obtained only on the second sample were not.

This two-sample structure reflects a sequential pilot-powered design. The first sample was used to explore candidate effects and estimate effect sizes. Based on these results, we preregistered hypotheses and an analysis plan for the second sample, which was independently powered to test the identified effects. All statistical inferences are based on the second sample. Full results from the first sample are reported in Appendix A for transparency, but were not used for inferential conclusions.

### 2.3. Participants

The participants recruited for this study were healthy, with intact or corrected vision, and aged 18 - 40. The pre-defined experiment exclusion criteria were: participants with dizziness, pregnant women, and participants with eyeglasses only for near vision correction (not contact lenses; according to eye-tracker manufacturer instructions). Participants gave their informed consent to participate in the experiment and received monetary compensation for their time of ∼ 12$ per hour. The study was approved by the institutional ethics committee.

Participants were recruited through multiple channels, including advertisements posted on campus, announcements in designated Facebook groups, and outreach via friends and colleagues of lab members.

We utilized a mixed design. Each participant was assigned to one of three VR conditions: 3D HMD with physical walking, 3D HMD with controller-based movement, or 2D desktop navigation, so that each participant experienced only one condition (between-subjects). Within each condition, participants completed two sessions with complementary maze setups. The assignment to a pair of setups (out of eight possible configurations) was counterbalanced across participants (for additional details on the maze setups and procedure, see section *2.5.2* *Behavioral Task*). The sessions and the repeated trials within each session served as within-subjects factors.

This study was performed on two samples. We aimed to obtain 48 valid participants for the first sample, 16 in each of the three experimental conditions. This resulted in a target of two participants in each setup option and each experimental condition. For this purpose, we recruited a sample of 53 participants. Two participants did not complete the task due to their difficulty using the mouse and keyboard to navigate the environment, which led to the incompletion of the experiment at its allotted time. Participants were considered valid if the data for the two task sessions were collected and saved properly. Five participants had only one valid session due to software failure and shutdown in the middle of the session. Therefore, these participants were excluded from further analysis. This concluded the first sample with 46 valid participants (mean age: 26.4, SD:5.3, 27 females).

We performed a bootstrapped power analysis on the behavioral results of this sample and calculated the minimal sample size required to obtain 80% power with p-value <= 0.05 for the effect sizes that were obtained on the first sample. Power analysis resulted in a target sample of 75 valid participants, 25 in each experimental condition, which was noted in the preregistration: https://osf.io/4b7kx. A total of 88 participants were recruited, and recruitment was stopped once 75 valid participants were obtained (mean age: 27.48, SD:5.3, 49 females). Thirteen participants were excluded due to the following reasons: one participant stopped the experiment due to their report of dizziness; two participants did not learn to navigate the environment in the experiment’s allotted time (one in the 2D condition and one using the controller); and ten participants had only one valid session due to software failure or shut down in the middle of the session. The demographic details of the two samples are shown in Table 1.

**Table 1:** Participants’ demographic data.

| S (N) | Condition (N) | Age mean (SD) | Frequency of playing computer games mean (SD) | VR experience N (%) | Females N (%) |
| --- | --- | --- | --- | --- | --- |
| **1 (46)** | 2D (15) | 25.6 (5.2) | 2.1 (1) | 11 (73) | 6 (40) |
|  | Controller (15) | 26.3 (5.4) | 1.6 (0.9) | 12 (80) | 11 (73) |
|  | Walk (16) | 27.5 (5.3) | 1.75 (1.1) | 9 (56) | 10 (67) |
| **2 (75)** | 2D (25) | 27 (4.5) | 1.96 (1.4) | 17 (68) | 13 (52) |
|  | Controller (25) | 26.9 (5.9) | 1.92 (1) | 15 (60) | 18 (72) |
|  | Walk (25) | 28.5 (5.1) | 1.76 (1.3) | 21 (84) | 18 (72) |

The participants' demographic data is presented according to sample (noted as S) and experimental condition. The frequency of playing computer games was selected on a 5-point scale from 1 (not at all) to 5 (every day). SD stands for standard deviation (see section 3.7 *Demographic data* in the *Results* for the corresponding statistical analysis).

Although both samples were recruited similarly, drawn from the same population, and tested under identical experimental conditions, the first sample served as a pilot to inform the preregistration of the second. Consequently, only the second sample was used for confirmatory statistical inference, and the two samples were not combined.

### 2.4 Experimental setup

All the virtual environments were executed on a computer equipped with an i9-10900f CPU running at 2.80 GHz, 64 GB of RAM, an NVIDIA GeForce RTX 2080 super GPU, and a 64-bit Windows 10 operating system. The virtual environment and task code were developed in C# in Unity 3D (version 2019.4). The 2D virtual environment was displayed on a screen with a 1920× 1080 resolution and a 60 Hz refresh rate.

The 3D VR conditions (the walking and using the controller conditions) were performed using a wireless VR headset adapted with a built-in binocular eye-tracking (ET) device (HTC VIVE Pro Eye). The eye-tracker (manufactured by Tobii^®^) provided data on eye openness, pupil diameter, eye gaze, and head movement during participants’ engagement in the VR task. The manufacturer’s ET accuracy was 0.5-1.1^0,^ and the sampling frequency was 120 Hz. However, the samples were collected at ∼90 Hz de facto, probably due to using the wireless adaptor rather than a direct connection to the computer. The VIVE Pro Eye displayed a resolution of 1440 x 1600 pixels per eye, with a trackable field of view of 110 degrees and a 90 Hz refresh rate. The acquisition of ET data was seamlessly integrated into the task through the HTC SRanipal software development kit (SDK) for Unity, which was customized to meet the task’s design and specifications. Eye-tracking was only implemented in the 3D HMD conditions; no eye-tracking data were collected in the 2D desktop condition.

The VR task and scene were designed according to the design of Barnes et al. (Barnes et al. 1980). The objects were uniquely modeled in Blender and imported to Unity. The participants controlled navigating the embedded task instructions using a trigger press on the controller in the VR conditions or the escape key in the 2D condition.

### 2.5 Experimental procedure

With the purpose of studying individual differences in spatial learning strategies, we translated the classical spatial learning 3-arm T-maze task (Barnes et al. 1980) for the first time to humans in VR.

#### 2.5.1 General Design

The maze was centered in the middle of the VR arena and was surrounded by five external cues (two of them are presented in the scheme in Fig. 1). In addition to these cues, one of the arms was textured with brick texture, differently than the others, according to task conditions as was the cued arm in the original task. The illustration of the environment shown in Fig. 1 is characterized by a right-cued arm. The task consisted of navigating to one of two arms in each trial to find the reward (in Fig. 1 it is positioned at the right arm).

#### 2.5.2 Behavioral Task

The task started with training trials by placing the participant at the beginning of the start arm. Each trial finished when the participant reached the end of one of the two T arms to find the reward which was a golden coin (Fig. 2). When participants reached the end of the rewarded arm, a golden coin appeared in front of them (in the right arm in the illustration in Fig.1), and upon their collection of it the message “+1 coin added” appeared in front of them. When participants reached the end of the unrewarded arm the message “+0 coin added” appeared in front of them. We added a return path to the beginning of the start arm between each two consecutive trials. As in the original article (Barnes et al. 1980) once four consecutive success trials were achieved the probe phase began.

**Fig. 2.**
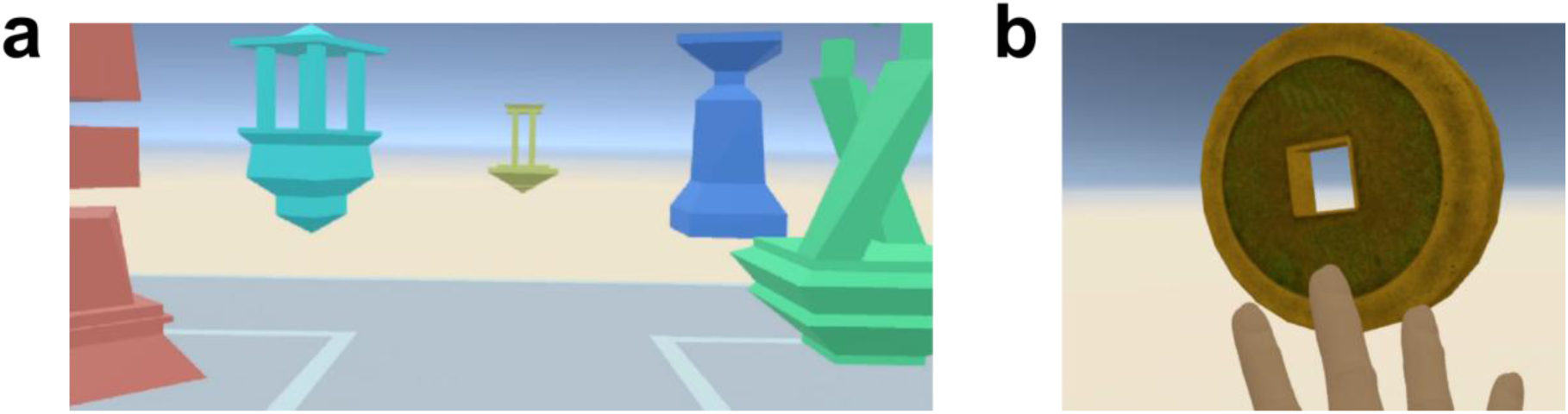
The T-maze setup. **a:** The view when entering the center of the T-maze from the start arm. **b:** A representation of a participant’s hand reaching to collect the found coin (link to the task video https://youtu.be/8gXAYO-h6qs)

Following the training phase, three types of test probe trials were performed, during which a turn to one of the maze arms represented an explicit choice in the strategy tested over the two others. Each strategy testing probe trial was preceded and followed by a regular probe trial identical to a training trial (see Fig .1). Therefore, the test probe trials were given as a series of six trials, where trials 1, 3, and 5 were regular trials (set up as in the training phase) and trials 2, 4, and 6 were each of the three test probe trials that were organized in a random order (an example for such a setting is shown in Fig. 1). Three consequent probe series were performed, which generated three test probe trials for each strategy.

To avoid new learning of the probe setups or extinction of the training, during the probe trials, both goal arms contained a masked silver coin, and upon the participants’ collecting it, the message “? coin added” appeared in front of them. Before they began the task, the participants were informed that the value of a silver coin could either be equivalent to that of a gold coin or have no value at all. The participants were instructed to find as many gold coins and silver coins with gold coin value as possible and were incentivized to receive an additional monetary bonus proportional to the number of these collected coins (without their knowing that all of them received the same bonus of ∼2.6$ at the end). We explained to the participants that these are two distinguished parts of the task and that from the moment they would first find a masked silver coin, they would always find masked silver coins.

##### Cue probe

The textured arm interchanged with the other (Fig. 1 (2)). When referring to the example presented in Fig. 1, in case the participant continued to turn to the cued arm they used a different turning response (left instead of right) and ended in a different place. Thus, by elimination, they used the ‘cue’ strategy.

##### Place probe

The entire maze was rotated by 180^0^ (Fig. 1 (4)). This implies that, for the example presented in Fig. 1, if the participant turned to the same spatial place, they used a different response and went to an arm with a different cue. This implies that they used a ‘place’ strategy.

##### Response probe

The start arm was rotated by 180^0^ (Fig. 1 (6)). For the example presented in Fig. 1, if the participant used the same turning response (right) they went to an arm with a different cue and ended in a different place. Hence, this participant used a ‘response’ strategy.

The participants were assigned to one of the 8 counterbalanced conditions that determined the maze setup: whether the textured arm (cue) was the rewarded arm, the position of the goal arm to the left or the right of the start arm (response), and the location of the start arm in relation to the spatial cues (place).

Each participant performed two sessions that included training trials and the subsequent probe trials for two complementary maze setups, such that no matter which strategy was used, new learning was required to find the reward during the second session.

We compared three levels of immersion and locomotion (Fig. 3): 1) participants wore a 3D HMD VR headset and physically walked in the arena (Fig. 3A); 2) participants wore a 3D HMD VR headset and used the VIVE controller for movement while they were physically standing in place (Fig. 3B); and 3) participants set in front of a 2D regular screen display and used the mouse and keyboard for movement (Fig. 3C). In all experimental setups, virtual avatar of the participant’s dominant hand appeared in front of them and changed position and perspective according to their movement in the scene. In the VR conditions, the participants reached out their dominant hand that held the controller to collect the coin when they approached it. In the 2D condition, the participants moved towards the coin using the mouse and keyboard until it collided with the coin and picked it up as a result.

**Fig. 3.**
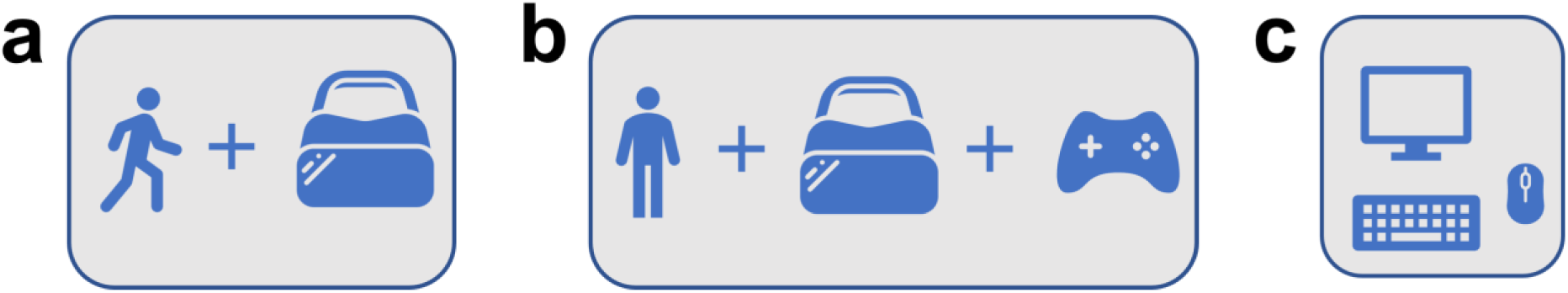
Experimental groups. **a:** 3D HMD with actual walking. **b:** 3D HMD standing and walking with a controller. **c:** 2D screen display using a mouse and a keyboard for movement

### 2.6 Raw data

The data was logged with a time stamp readout in microseconds precision that allowed synchronization of the data with the task events timeline. In all the experimental conditions, the 3D (x,y,z) position and rotation of the participants were recorded. In the 3D HMD conditions additional ET measurements were gathered with the SRanipal SDK including 3D (x,y,z) gazed ray origin and direction, pupil diameter (mm), eye-openness (EO), and sample validity defined by the eye-tracker. We added the collection of the following variables: the gazed object name; the 3D (x,y,z) gazed object position, for which we considered the center of the object’s bounding box; and the 3D (x,y,z) gaze ray hit point, which was the location of collision of the gaze ray with the gazed object collider. Samples validity and time stamps were used for trial validity analysis as detailed in the *Eye-tracking data pre-processing* section below.

### 2.7 Data handling

Data extraction, parsing, arrangement, fusion, and processing were performed with Python 3.12.3 (https://www.python.org/).

### 2.8 Eye-tracking data pre-processing

ET samples with EO ≤ 0.5 were considered as blinks. NaN values or non-valid samples, as defined by SRanipal SDK, were removed from the analysis. Samples in a time window of 150 ms before and after each blink were scrubbed to remove transient signal artifacts surrounding blink onset and offset, consistent with previous practice in the lab (Salomon et al. 2019) and recent eye-tracking and pupillometry studies (Yoo et al. 2021).

Gaze metrics were computed directly from the gaze hit points on objects, without applying fixation-detection or duration-filtering algorithms. This object-based approach is standard in immersive 3D environments, where gaze samples correspond to collider hits rather than continuous 2D screen coordinates (Clay et al. 2019; Schuetz and Fiehler 2022; Hou et al. 2024; Hirschhorn et al. 2024), and avoids variability introduced by fixation-detection models (Komogortsev et al. 2010; Holmqvist et al. 2012; Andersson et al. 2017).

### 2.9 Extracted measures

#### 2.9.1 Learning measures

We defined the participants’ learning pace as the number of training trials to reach the learning criterion (the minimum was 4 trials). Learning effectiveness was considered as the number of successful probe trials that were identical to training trials (out of a total of 9 trials). Eventually, for each of the three strategies test probe trials, we examined the explicit choices in the tested strategy.

#### 2.9.2 Path measures

For all VR modalities, we extracted 10 descriptive path measures. We examined the path length and average speed from several perspectives: the entire trial, from the start position to the center of the maze (the section from point *S* to point *C* in Fig.4), from the center to the start of one of the maze’s arms (the section from point *C* to point *A* in Fig. 4), and from the beginning of the maze arm to its end (the section from point *A* to point *E* in Fig. 4). In addition, we calculated the number of visits to the maze’s center (point *C* in Fig. 4), and the number of enters to arms (visits in point *A* in Fig. 4) as measures of exploration or uncertainty in the trial, where the minimum per trial for both is one.

**Fig. 4.**
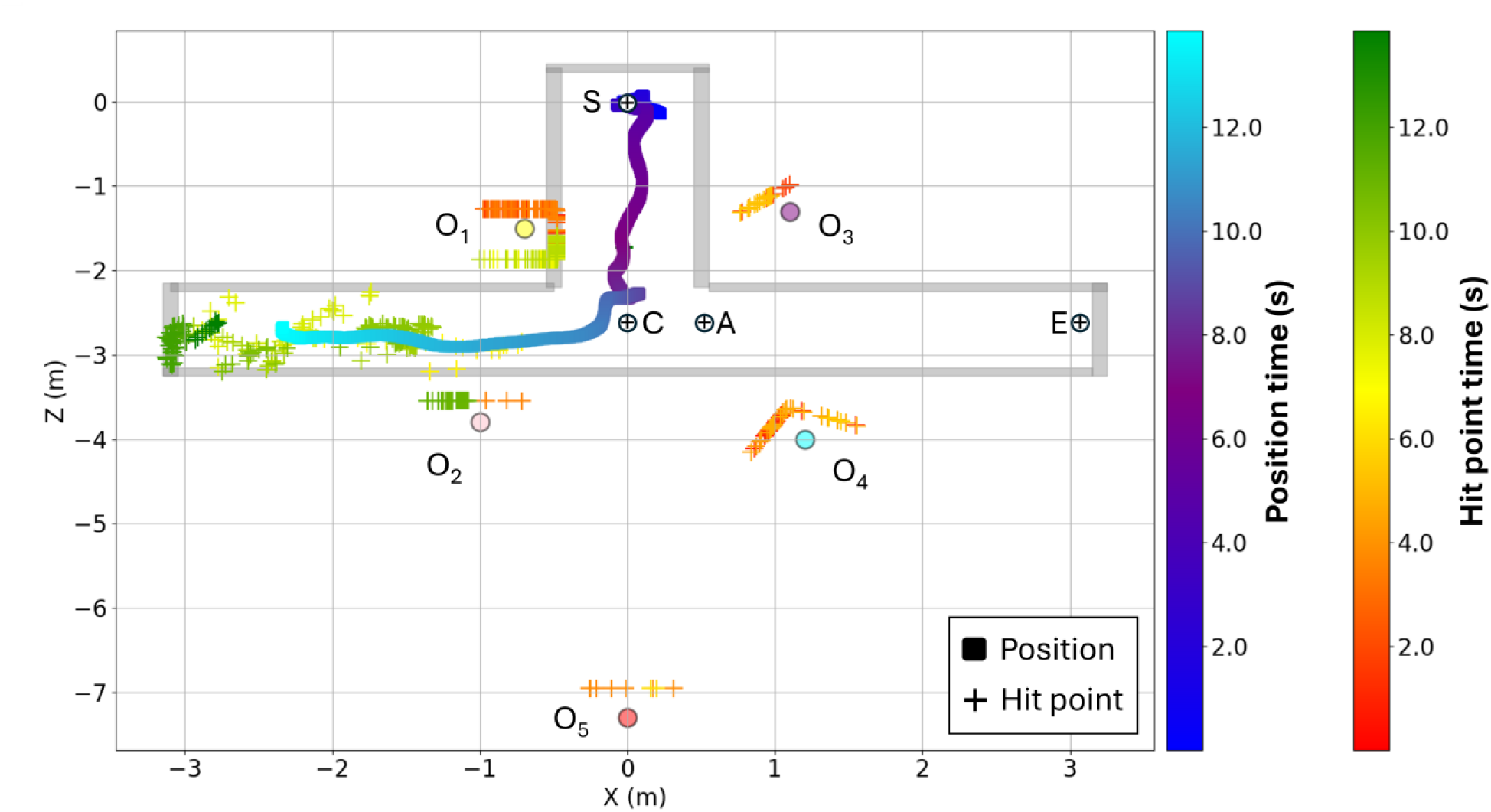
Physiological and behavioral data in the maze. Scatter plot of a participant’s data in one training trial. The participant’s position in the scene (filled rectangle) and their gaze hit point position (plus marker) are displayed on top of the top view of the maze scenery in the arena. The dimensions in the scheme represent the rendered physical dimensions in the virtual environment which are in 1:1 ratio to the real-world scale in meters. The data are color-coded by the time relative to the beginning of the trial, with blue to cyan colormap for the participant’s position and red to green colormap for gaze hit points position. Five colored circles with the notion O1 - O5 represent the position of the center of the maze’s five surrounding objects. The colored plus markers surrounding the centers of these objects represent the gaze hit points while looking at them and are positioned on their corresponding collider object. The points marked by a white circle with a black cross in the middle denote physical locations of interest in the maze, which are relevant to the path analysis. The starting point is denoted by S, the center of the maze by C, the beginning of one arm by A (the left one with respect to the participant is shown as an example), and the end of that arm by E (the left one with respect to the participant is shown as an example)

#### 2.9.3 Eye-tracking measures

We extracted ET features to encompass the gaze behavior in every trial based on (Gabay and Schonberg 2023), in which these measures were positively associated with participants’ subjective rankings of preferences and valence. We hypothesized that they could similarly capture individual differences in visual exploration of the environment and contribute to the prediction of spatial learning strategies. For each trial, we extracted global features that comprised of 6 pupil features, 3 per eye, pupil diameter (10th, 50th, 95th percentiles); and 5 blinks features including number of blinks, total blinks duration, minimum, maximum, and mean of blinks length based on constraints (Pietrock et al. 2019). In addition, we extracted object-related features that consisted of: 1 feature of percent-time the object was gazed at out of the entire trial duration;12 dispersion patterns of gaze hit points features which comprised of the (10, 50, 95) percentiles of the Euclidian distance between the gazed hit point to the centroid of the gazed object that we named as gaze shift distance; these distances normalized by the Euclidian distance between the gazed ray origin and the gazed object centroid; the difference in time of the gaze shift distance, and the Euclidian distance between the participant’s head position and the gazed object position; and 3 gaze speed features defined as the (10, 50, 95) percentiles of the gaze scan Euclidian distance per time unit. The object-related features were computed to assess gaze behavior while looking at a single object or at a cluster of objects with a common trait of interest during the trial. An example of gaze behavior in a trial is shown in Fig. 4.

### 2.10 Questionnaires and tests

Following the completion of the two task sessions, the participants filled out several questionnaires on a 2D screen display. We first collected their demographic data (see details in Table 1) and continued with other thematic questionnaires and tests that were performed in the order they are mentioned in this section. The questionnaires were presented on a Google form unless mentioned otherwise.

#### 2.10.1 Presence questionnaire

The sense of spatial presence was collected using a Hebrew translation of the ITC SOPI (Independent Television Commission Sense of Presence Inventory) presence questionnaire (Lessiter et al. 2001). This questionnaire consists of 38 statements for which the participants were asked to indicate the level of their agreement using a 5-point scale from 1 (strongly disagree) to 5 (strongly agree). The results were combined into 4-factor scores: Spatial presence (e.g., ‘I had a sense of being in the scenes displayed’), Engagement (e.g., ‘I vividly remember some parts of the experience’), Ecological Validity/ Naturalness (e.g., ‘The displayed environment seemed natural’), and Negative Effects (e.g., ‘I felt I had eyestrain’). Each factor is the mean value of all the items defined as contributing to the factor.

#### 2.10.2 Awareness questionnaire

The participants were asked for their awareness of the information each of the three strategies held in the task in a three-choice question consisting of: “yes”, “in some of the trials”, and “no”. For example, to test their awareness level of the cue, they were asked-“have you noticed a brick texture in one of the arms?”. The responses were transformed into a 3-level ordinal scale from 1 (“no”) to 3 (“yes”) and were used for analysis.

In addition, the participants were asked for their perception of to what degree they used each of the three strategies information on a five choice-scale of: “yes, in all trials”, “in most trials”, “in half of the trials”, “in less than half of the trials”, and “not at all, it seemed to me as a decoration”. For example, to test their perception of the level of using the cue strategy, they were asked-“have you used the arms’ texture (with/without bricks) to collect as many golden coins as possible?”. The responses were transformed into a 5-level ordinal scale from 1 (“not at all, it seemed to me as a decoration”) to 5 (“yes, in all trials”) and were used for analysis.

#### 2.10.3 Spatial abilities tests

##### Mental Rotations Test (MRT)

The MRT requires participants to identify two out of four rotated 3D objects that are identical to a reference figure (Vandenberg and Kuse 1978) later redesigned by (Peters et al. 1995). This task assesses spatial ability by challenging participants to mentally manipulate and compare the orientations of complex shapes within a time limit. We used a Hebrew version of the validated revised version of a standard set MRT-A test that was translated and used in (Lander et al. 2020). The MRT-A version included 24 sets of images where a point was given only when the two matching figures were identified. Therefore, the test score range was 0 to 24. Following the instructions, the participants practiced four sets, and then they were given three minutes for the first two pages, and after a short break, additional three minutes for the next two pages.

##### Spatial Orientation Test (SOT)

To examine spatial orientation ability we used a perspective-taking task that requires imagining how a scene looks from different angles (Hegarty and Waller 2004; Friedman et al. 2020). The participants were shown a spatial layout of objects and asked to imagine themselves at a specified location of one of the objects within the layout while facing another object. Then they indicated the relative direction of a target object from this imagined perspective, assessing their ability to mentally shift viewpoints. Given that the paper-based and the computer-based versions of the SOT produced similar performance (Friedman et al. 2020), we translated the computerized version to Hebrew with Python (task code available at: https://github.com/Yehuda-Bergstein/ptsot). After receiving the instructions, the participants practiced marking the answer with the mouse and were then given three examples with feedback. Lastly, the test included 12 trials with a time limit of 5 minutes. For each trial, the angular error was calculated as the deviation in degrees between the participant’s response and the correct direction, with smaller errors indicating better spatial orientation ability. Performance was scored as the mean angular error of all 12 test trials. Three participants with a mean SOT angular error greater than 90^0^ were excluded from the SOT analyses, as in (Kapaj et al. 2024), as this error is higher than chance performance.

### 2.11 Statistical analysis

Statistical analysis and figures were coded with R version 4.3.1. Mixed linear and mixed generalized linear models were formulated with lme4 (Bates et al. 2015) and contrast analysis with emmeans (Lenth 2024) packages. We used ggplot2 (Wickham 2016) for visualizations. The standard p-value of 0.05 was considered for assessing the significance of each effect. Bayes Factor analysis was formulated with BayesFactor (Morey and Rouder 2012), brms (Bürkner 2017), and bridgesampling (Gronau et al. 2020) packages.

Power analysis was implemented using a non-parametric bootstrap procedure applied to the behavioral data of the first sample, with custom code. Effect sizes were estimated by resampling the observed data to approximate the population distribution, and power was calculated for detecting these effects at α = 0.05. This procedure assumes that the pilot data distribution represents the population and that observations are independent.

#### 2.11.1 Learning measures

We analyzed the learning pace and learning effectiveness using mixed linear models with experimental condition and session number as fixed effects, and participants were considered as random effects. Since Shapiro–Wilk test showed that some of the data were not normally distributed, the models were performed on the rank of the learning measures. For each of the three strategy test probe trials, we analyzed the probability for an explicit choice in strategy using a generalized linear mixed model (GLMM) with a binomial distribution.

To enhance the interpretability of the GLMM, we calculated the Standardized Mean Difference (SMD, Cohen’s *d*) for all reported Odds Ratios (ORs) (Chinn 2000). This conversion was performed using the standard formula in equation 1:

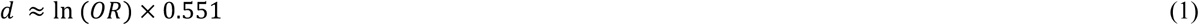

The need to evaluate the magnitude was guided by the principle articulated by (Azuero 2016).

The models’ default was the VR walking condition and the first session.

#### 2.11.2 ET measures

To test whether the level of immersion and locomotion affected gaze behavior, we examined general ET features and compared the two VR conditions. We analyzed ET measures obtained on all trials using mixed linear models with experimental condition and session number as fixed effects and participants were considered a random effect. The models’ default was the VR walking condition.

As an exploratory analysis, which was therefore not preregistered, we further examined whether visual attention to task-relevant objects predicted spatial learning performance. For each performance measure (learning pace and learning effectiveness), we ran three separate mixed linear models, with the average gaze time across training trials on (i) the cue on the reward side, (ii) reward-side objects, or (iii) all objects as fixed predictors, together with condition and session number. Participants were included as random intercepts, and SOT and MRT scores were added as covariates when available. Consistent with section *2.11.1* *Learning measures*, the models were performed on the rank of the learning measures.

#### 2.11.3 Presence factors

The effect of the level of immersion on the sense of presence was examined using linear models, where each of the 4 presence factors was a dependent variable and the experimental condition was a categorical fixed effect. Since these factors were of ordinal scale, the models were performed on the rank of the presence factors. The models’ default was the VR walking condition.

#### 2.11.4 Awareness measures

We examined the relation between the choice of strategy to the awareness level of the information in the scene each strategy holds, the perception of using it, and whether they were affected by the VR modality. For this purpose, we used linear models, where the dependent variable was the average number of choices in strategy across the two sessions for each strategy test probe trial, and the awareness of the strategy, the perception of its usage, and the experimental condition were considered as independent variables (including interaction with the condition). Since these variables were of ordinal scale, the models were performed on the ranks of the variables. The models’ default was the VR walking condition.

#### 2.11.5 Spatial abilities measures

The spatial abilities tests were collected only from the second sample. Therefore, we did not preregister their analysis. We tested the correlation between spatial abilities and the tendency to use each strategy with linear models. In each model, the dependent variable was the mean across sessions of the number of explicit choices in the tested strategy, and the VR modality and each of the test scores of the MRT and SOT were the independent variables including their interaction. Since the dependent variables were of ordinal scale, the models were performed on the ranks of the variables. The models’ default was the VR walking condition.

We tested differences between the participants’ spatial abilities across experimental conditions with linear models, where the dependent variable was each of the test scores of the MRT and SOT, and the VR modality was the independent variable.

#### 2.11.6 Demographic data

The differences between the two samples’ demographic data were not preregistered. We added them following the second sample’s data analysis. Some inconclusive findings between the two samples drove us to test for differences in the demographic data, as they might contribute to them. Each demographic variable was examined by a linear model with sample number and condition as independent variables (including their interaction).

### 2.12 Bayes factor analysis

All Bayes factor analyses were not preregistered. We used them as supportive analyses in cases where the results were inconclusive among the two samples, and where significant results were obtained only in one of them. These analyses were added only to the models with results that were not replicated in the second sample or found in the second sample and not in the first.

In the Bayes factor analyses, H1 and H0 denote the alternative and null statistical hypotheses for each tested effect, representing the presence or absence of the effect under examination, respectively.

Significant effects were further assessed with BF10, which is the ratio of evidence in favor of the alternative hypothesis (H1) over the null hypothesis (H0). On the other hand, in cases where no significant effects were obtained, we computed BF01, which is the ratio of evidence in favor of the null hypothesis (H0) over the alternative hypothesis (H1). For each analysis, the alternative hypothesis was formulated by the full model with all the fixed effects, intercept, and random effects. The null hypothesis was represented by the same model as the full model, without the fixed effect of interest.

To quantify the strength of evidence for one hypothesis over another we considered the widely used frameworks (Kass and Raftery 1995; Jeffreys 1998) that provide specific thresholds guiding interpretation. Values between 1 and 3 indicated anecdotal evidence, reflecting weak support; values from 3 to 10 suggested moderate evidence, providing substantial support for one hypothesis; values from 10 to 30 represented strong evidence; values between 30 and 100 indicated very strong evidence; and values exceeding 100 signified extreme or decisive evidence.

For the linear and mixed linear models, we used a wide Cauchy prior distribution with r-scale (Rouder et al. 2012) value of 0.5 and tested the robustness of the results with additional scaling factors of 1 and √2. We used Normal (0, 1.5) priors for GLMM fixed effects as recommended in (McElreath 2018), and Normal (0, 2.5), and Normal (0, 10) for the priors sensitivity analysis.

### 2.13 Prediction models

We designed machine learning (ML) models to predict individual strategy usage based on physiological ET and behavioral measures. The models were computed with the scikit-learn (Pedregosa et al. 2011) library, Python 3.12.3. This section was exploratory and was not preregistered.

A total of 10 path measures were used for all three VR modalities (detailed in section *2.9.2* *Path measures*). For the two VR conditions, we additionally used 75 ET features consisting of 6 pupil features, 5 blinks features, and 4 groups of object-related features with 16 features in each group (see details in section *2.9.3* *Eye-tracking measures*). The object groups included: the surrounding objects on the reward side of the maze (objects *O_1_*, and *O_2_* in Fig.4 for example if the reward was on the right side), the surrounding objects on the opposite side to the reward (objects *O_3_*, and *O_4_* in Fig.4 for example if the reward was on the right side), the cue on the reward side of the maze (the right arm of the maze in Fig. 4 for example if the reward was on the right side), and the cue on the opposite side to the reward (the left arm of the maze in Fig. 4 for example if the reward was on the right side). The scene design in the training phase determined the reference to the side of the reward of the objects and the cue for each participant.

All features were fed into support vector machine (SVM) models with radial basis function (RBF) classification or regression to predict the individual strategy usage. The models’ performance was assessed using leave one subject out cross-validation (CV). On each fold, the missing values were first filled with the median of the features. Then, backward elimination feature selection was performed to decrease the number of features in the model and reduce the chance of overfitting. We constructed three categories of models as follows:

1. Trial-by-trial classification of the choice in strategy at each test probe trial based on the features extracted during that trial.
2. General preference of strategy classification based on the probability of using it calculated on all test probe trials as in Barnes *et al*. (Barnes et al. 1980). The models were fed with the median value of the features across all training trials or all trials.
3. Regression of the number of explicit choices in strategy. The models were fed with the median value of the features across all training trials or all trials.

The models’ performance was assessed by accuracy for the classification models, and root mean square error (RMSE) and R^2^ for the regression models. The statistical significance of the models’ performance measures was assessed using permutation tests with 100 iterations. The permutation tests were performed only for models that obtained accuracy greater than 0.5 in the classification models, and R^2^ greater than 0 in the regression models. Otherwise, the P-value was noted as 1.

As this section represents an exploratory analysis, our focus was on estimating classification performance across multiple configurations rather than on detailed per-model error diagnostics. Therefore, to support interpretability while maintaining clarity, we did not include confusion matrices for each binary classifier (separately trained per strategy, session, condition, and sample). Furthermore, since each model was trained and tested using leave-one-subject-out cross-validation, each fold generated an independent model with its own confusion matrix and feature-importance profile. We did not summarize or report these per-fold metrics, which would have likely been unstable due to the small sample size per group and difficult to interpret in aggregate. Instead, we summarize classifier performance using balanced accuracy and permutation-based p-values.

To increase model performance interpretation, we also extracted the positive class proportion for each model, which is the proportion of trials or participants labeled as 1 (e.g., strategy chosen or preferred). Since significance testing was conducted via permutation, this proportion may help contextualize the likelihood of observing a given accuracy under the null. In particular, when class distributions are highly imbalanced, even high accuracies may not be statistically meaningful. Including the positive class proportion, therefore, supports transparent evaluation of classification performance under varying label distributions.

## 3 Results

Unless stated otherwise, the results presented in this chapter are based on the preregistered powered sample (the second sample), which was powered based on effect sizes observed in the first pilot sample. In sections 3.1–3.2 and 3.4-3.5, only the second sample is reported, and results from the first sample are included in *Appendix A* (sections A.1–A.4) for transparency but are not used for inferential conclusions. Where relevant, we briefly indicate whether an effect observed in the second sample also appeared in the first sample. Sections 3.3, 3.6, and 3.7 follow a different structure: section 3.3 includes exploratory analysis, therefore presents both samples’ results; section 3.6 presents data that were collected only for the second sample; and section 3.7 directly compares the two samples.

### 3.1 Learning measures

#### 3.1.1 Learning pace

The average number of training trials required to reach 4 consecutive successes across all samples, conditions, and sessions was 7.8 trials. In the powered sample, there was no difference in the learning pace between the two VR conditions (*estimate* = −1.4, *t*_(144)_ = −0.1, *p* = 0.91), a finding also observed in the first sample. In addition, no significant difference was found between the VR walking and 2D condition (*estimate* = 10.9, *t*_(144)_ = 0.9, *p* = 0.36, Fig. 5), with anecdotal support in favor of the null hypothesis (BF01 = 1, 1.2, and 1.2), which did not replicate the significant effect observed in the pilot sample (see *Appendix A.1* and Supplementary Table A.1.1).

**Fig. 5.**
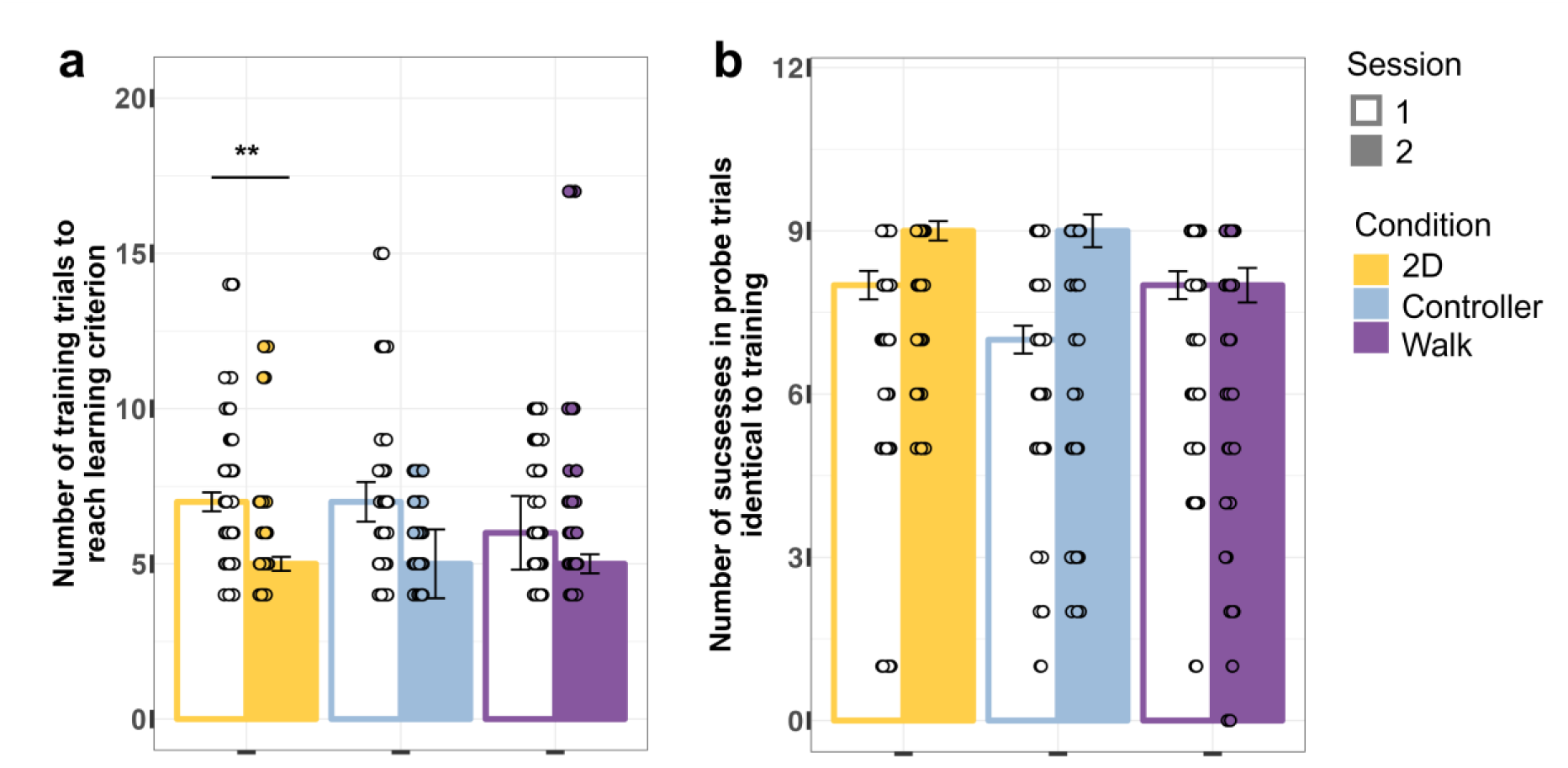
Learning measures. Bar plots of learning measures of the second sample according to VR modality and session. Since the models were performed on the measures’ ranks, the height of each bar presents the median, with error bars indicating ± 1 SEM (standard error of the mean). Individual data points are shown as dots overlaid on the bars, with y axis limit of mean ±2 SD (standard deviations). (p < 0.01 ‘**’, p < 0.05‘*’)

Furthermore, a main effect for the second session was not obtained (*estimate* = −12.3, *t*_(144)_ = −1, *p* = 0.29), nor any of its interactions with the other experimental conditions (*estimate* = −24.2, *t*_(144)_ = −1.4, *p* = 0.15) for Condition2D x Session2, and (*estimate* = −2.1, *t*_(144)_ = −0.1, *p* = 0.89) for ConditionController x Session2. This pattern of null results replicated the findings from the pilot sample (*Appendix A.1*).

Although there was no significant interaction between the 2D condition and session 2, we specifically tested the contrast of learning pace in the 2D condition across sessions, as this effect was observed in the pilot sample. In the powered sample, participants learned significantly faster in the second session compared to the first in the 2D condition (*estimate* = 36.6, *t*_(72)_ = 3.1, *p* = 0.003, Fig. 5a, Bonferroni corrected for 3 tests), replicating the result observed in the first sample (*Appendix A.1*).

This suggests that the relatively low ecological validity of navigating 3D virtual space on a 2D screen may have hindered learning during the first session, but performance improved in the second session as participants gained task-specific experience. This interpretation is further supported by the presence questionnaire results detailed in section *3.4* *Presence measures*.

#### 3.1.2 Learning effectiveness

The average success rate on probe trials identical to training was 79.5% across all samples, conditions, and sessions. In the powered sample, there was no significant difference in the learning effectiveness between the VR walking and 2D conditions (*estimate* = 0.5, *t*_(134.1)_ = 0.04, *p* = 0.97, Fig. 5b). Bayes factor analysis provided only anecdotal to no evidence for the null hypothesis over the alternative (BF01 = 1, 1.1, and 1.1). This result did not replicate the significant difference observed in the pilot sample (see *Appendix A.1* and Supplementary Table A.1.1).

A contrast focusing on the second session was tested based on the significant difference observed in the pilot sample for learning effectiveness between the 2D and VR walking conditions. In the powered sample, this effect was not replicated, as no significant difference was observed (estimate = –15.7, t(134.0) = –1.32, p = 0.57, Bonferroni corrected for 3 tests; see *Appendix A.1* and Supplementary Table A.1.1). Bayes factor analysis found anecdotal to no evidence for the null hypothesis over the alternative (BF01 = 1, 1.1, and 1).

There was no significant difference in the learning effectiveness between the two VR conditions (*estimate* = - 5.28, *t*_(134.1)_ = −0.44, *p* = 0.66). In addition, there was no significant difference in the learning effectiveness between the two sessions, as a main effect (*estimate* = −0.32, *t*_(72)_ = −0.03, *p* = 0.97) or as an interaction with the controller (*estimate* = 11.52, *t*_(72)_ = 0.8, *p* = 0.42) or the 2D conditions (*estimate* = 15.2, *t*_(72)_ = 1.06, *p* = 0.29). These results were consistent with the findings from the pilot sample (*Appendix A.1*).

#### 3.1.3 Learning strategy

Only 20.6% of all participants across both samples and all conditions explicitly chose only one strategy in both sessions. More than double of them chose only one strategy during one session (44.6% of both samples’ participants). All other participants (55.4%) used a combination of strategies during the different probe trials and sessions.

##### Cue strategy

In the powered sample, there were no significant effects for the two experimental conditions, where (OR = 0.54, 95% CI = [0.18, 1.55], *p* = 0.25, *d* = −0.34) for the 2D condition, and (OR = 0.78, 95% CI = [0.27, 2.25], *p* = 0.64, *d* = −0.14) for the controller condition. These results replicate the null effects observed in the pilot sample (see *Appendix A.1*, Supplementary Fig. A.1.2, and Supplementary Table A.1.2).

Nevertheless, there was a significant main effect for the second session (OR = 0.39, 95% CI = [0.18, 0.82], *p* = 0.014, *d* = −0.52), as well as interactions between the second session and both the 2D and controller conditions (OR = 3.73, 95% CI [1.29, 10.81], *p* = 0.015, *d* = 0.73) for Condition2D x Session2, and (OR = 3.25, 95% CI [1.11, 9.47], *p* = 0.03, *d* = −0.65) for ConditionController x Session2. Bayes factor analysis found moderate to extreme evidence in favor of the null (BF10 = 0.3, 0.1, and 0) for the session main effect, moderate to anecdotal support for the alternative hypothesis for the interaction with the 2D condition (BF10 = 3.8, 3.2, and 1.1), and mixed anecdotal evidence in favor of the null and alternative hypotheses for the interaction with the controller condition (BF10 = 2.3, 2, and 0.6). None of these effects were obtained in the pilot sample (*Appendix A.1*).

Contrast analysis revealed that the probability of using cue strategy significantly decreased in the second session relative to the first in the VR walking condition (OR = 2.57, 95% CI [1.2, 5.47], *p* = 0.014, *d* = 0.52, Fig.6), with moderate to no support for the alternative hypothesis (BF10 = 4.3, 2.9, and 0.8). This effect was also not observed in the first sample (*Appendix A.1*).

**Fig. 6.**
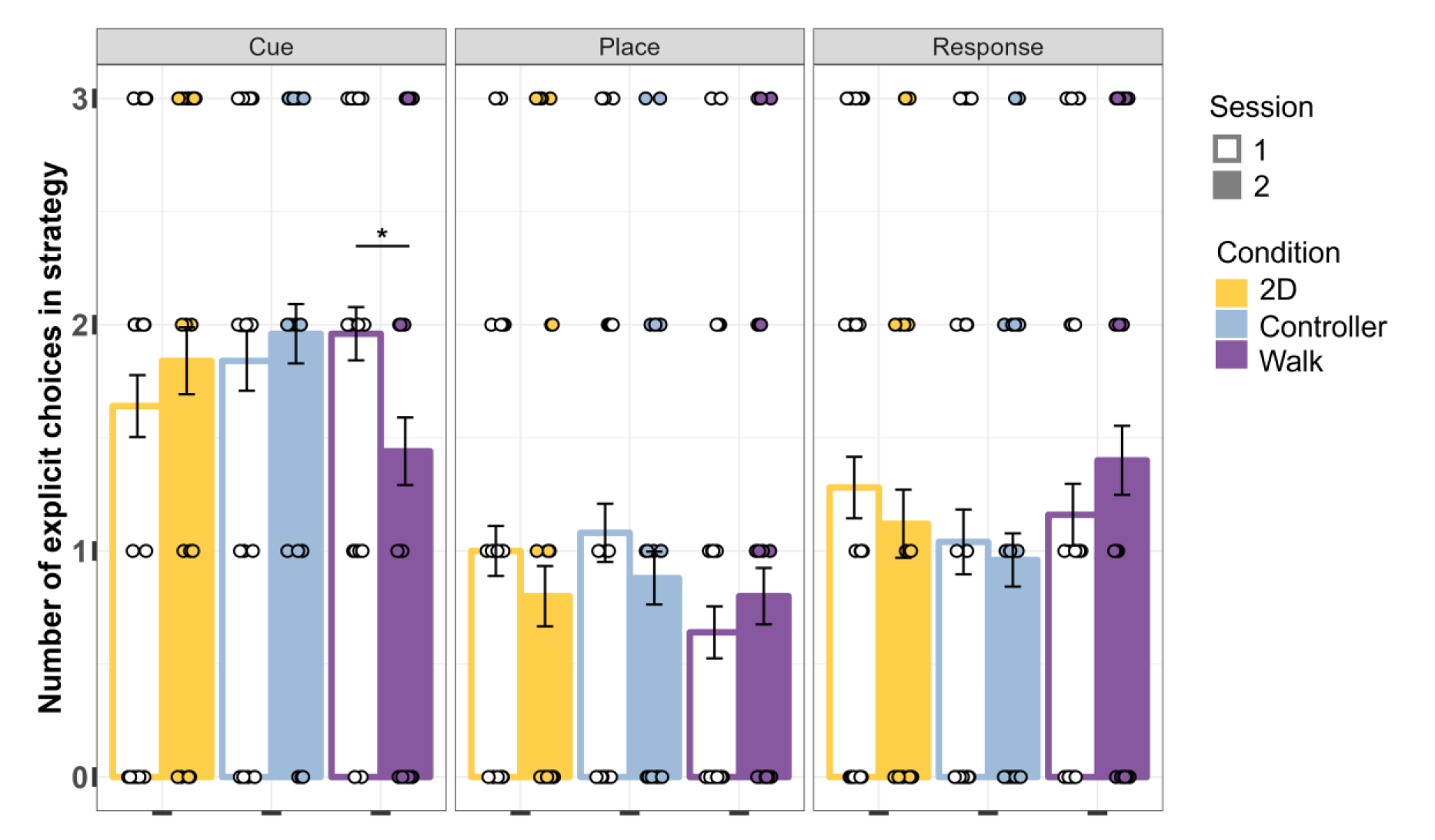
Choice of strategy. Bar plots of the number of explicit choices in each strategy during strategy test probe trials of the second sample according to VR modality and session. The height of each bar presents the mean, with error bars indicating ± 1 SEM. Individual data points are shown as dots overlaid on the bars. (p < 0.001 ‘***’, p < 0.01 ‘**’, p < 0.05‘*’)

##### Place strategy

In the model for the probability of an explicit choice of place strategy a significant main effect of the second session was not observed (OR = 1.45, 95% CI [0.62,3.39], *p* = 0.39, *d* = 0.2) nor was a significant interaction found between the second session and the VR controller condition (OR = 0.46, 95% CI [0.15, 1.47], *p* = 0.18, *d* = −0.43). Bayes factor analysis showed strong to extreme evidence in favor of the null hypothesis (BF01 = 18.7, 66, and 3360) for the session main effect, and anecdotal to moderate evidence that supports the null for the interaction with the controller condition (BF01 = 1.5, 2.1, and 7.5). These findings did not replicate the results obtained in the pilot sample (see *Appendix A.1*, Supplementary Fig. A.1.2, and Supplementary Table A.1.2).

Contrast analyses were conducted in the powered sample based on effects observed in the pilot sample. Specifically, we tested whether the probability of using the place strategy increased in the second session relative to the first within the VR walking condition, and whether it differed between the two VR conditions during the second session. Neither contrast yielded significant effects in the second sample (OR = 0.68, 95% CI [0.29, 1.61], *p* = 0.39, *d* = −0.21, Fig. 6, Bonferroni corrected for 3 tests) for the sessions VR walking condition contrast, and (OR = 0.99, 95% CI [0.26, 3.88], *p* = 1, *d* = −0.01, Bonferroni corrected for 3 tests) for the VR conditions contrast in the second session. Bayes factor analysis showed moderate to strong evidence for the null hypothesis, with (BF01 = 2.5, 4, and 16) for no difference between the sessions in the VR walking condition and (B01 = 2, 3.1, and 11.9) for the difference between the two VR conditions in the second session.

No significant main effects were found in the second sample for either the 2D condition (OR = 2.22, 95% CI [0.72, 6.81], *p* = 0.16, *d* = 0.44) or the controller condition (OR = 2.61, 95% CI [0.86, 7.98], *p* = 0.09, *d* = 0.53). Additionally, no significant interaction was observed between the 2D condition and session 2 (OR = 0.45, 95% CI [0.14, 1.46], *p* = 0.18, *d* = −0.44). These null results are consistent with the findings from the pilot sample (see *Appendix A.1*) and suggest that the probability of using a place strategy did not differ across sessions for the 2D and the controller conditions. *Response strategy:* In the powered sample, there were no significant differences in the probability of explicitly choosing a response strategy (turn left/right) between the VR walking condition and either the VR controller (OR = 0.86, 95% CI [0.27, 2.74], *p* = 0.8, *d* = −0.08) or the 2D condition (OR = 1.37, 95% CI [0.43, 4.35], *p* = 0.59, *d* = −0.17, Fig. 6). Bayes factor analysis found anecdotal to strong evidence in favor of the null (BF01 = 1.9, 3, and 12) for the controller condition, and (BF01 = 2.6, 3.8, and 17.3) for the 2D condition. These results did not replicate the significant effects found in the pilot sample (see *Appendix A.1*, Supplementary Fig. A.1.2, and Supplementary Table A.1.2).

A preregistered contrast analysis was conducted based on the significant effect observed in Sample 1. However, in the second sample, no significant difference was found in the first session between the VR walking and 2D conditions (OR = 0.72, 95% CI [0.18, 2.97], *p* = 1, *d* = −0.18, Bonferroni corrected for 3 tests), with Bayes factors indicating anecdotal to strong evidence in favor of the null (BF01 = 2.1, 3, and 11.1).

The session did not obtain a significant effect on the probability for using response strategy (OR = 1.64, 95% CI [0.75, 3.59], *p* = 0.21) for the session main effect (OR = 0.52, 95% CI [0.17, 1.75], *p* = 0.24, *d* = −0.36); for the ConditionController x Session2 interaction; and OR = 0.45, 95% CI [0.15, 1.33], *p* = 0.15, *d* = −0.44) for the Condition2D x Session2 interaction. These null findings are consistent with those from the pilot sample (*Appendix A.1*).

### 3.2 Eye-tracking measures

#### 3.2.1 Pupil size

Pupil size was assessed using the mean of the left pupil in each trial (after verifying similar results were obtained for the right pupil), following common practice in cognitive research (Partala and Surakka 2003; Laeng et al. 2011; Foroughi et al. 2017). Consistent with findings from the first sample, the pupil size was significantly smaller in the second session (*estimate* = −0.05, *t*_(1938)_ = −2, *p* = 0.04; see Fig. 7a). There was no main effect of condition (*estimate* = −0.14, *t*_(48.8)_ = −0.8, *p* = 0.44). However, an interaction between session and condition was found in the second sample (*estimate* = - 0.12, *t*_(1938)_ = −3.5, *p* = 0.0004), which was not observed in the first (see Supplementary Fig. A.4.1 and Supplementary Table A.4.1 in *Appendix A.4.1*).

**Fig. 7.**
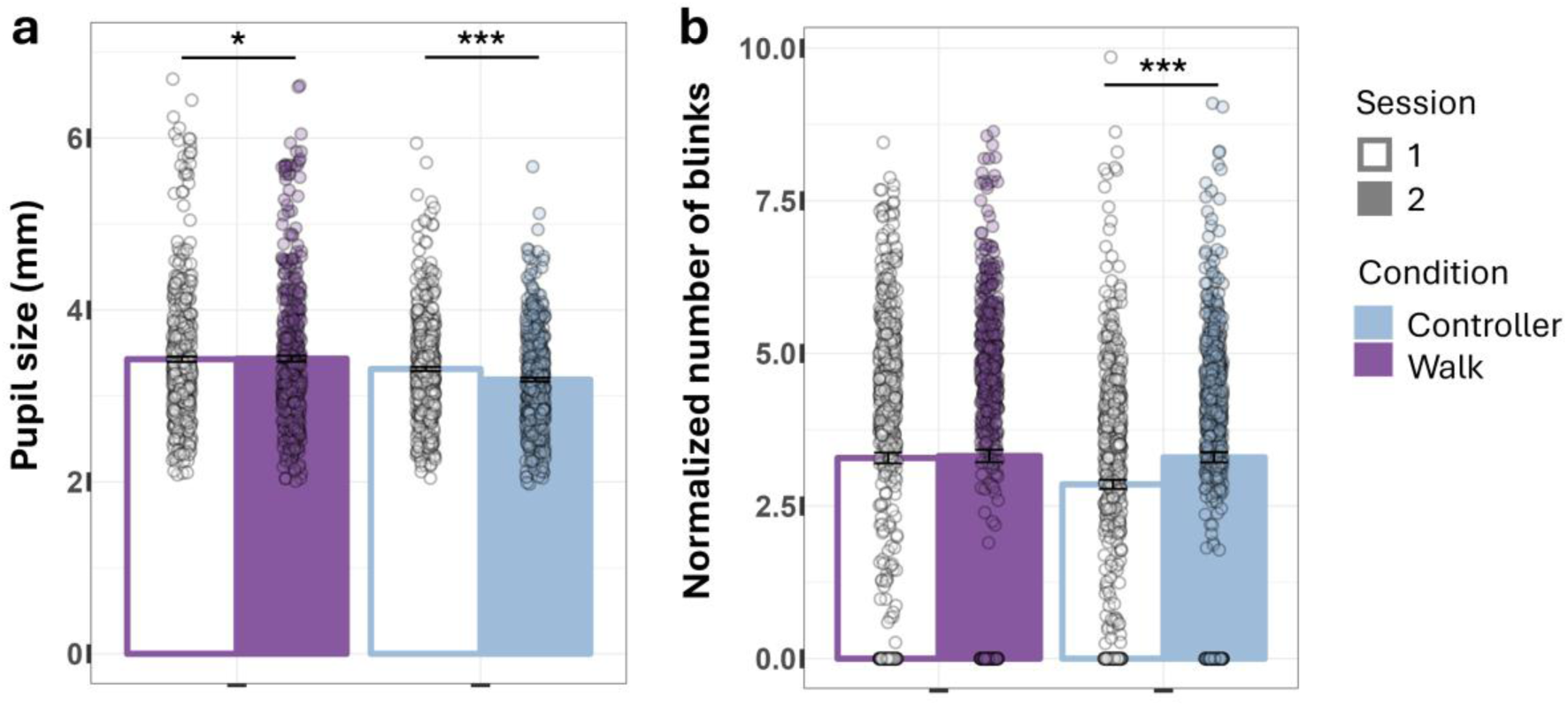
Eye-tracking (ET) measures. Bar plots of ET measures according to VR modality of the second sample. The height of each bar presents the mean, with error bars indicating ± 1 SEM. Individual trial-based data points are shown as dots overlaid on the bars. (p < 0.001 ‘***’, p < 0.01 ‘**’, p < 0.05‘*’)

Contrast analysis showed that the pupil size decreased from the first session to the second in both VR conditions, with (*estimate* = 0.05, *t*_(1938)_ = 2, *p* < 0.04) for the walking condition, and (*estimate* = 0.17, *t*_(1937)_ = 7.2, *p* < 0.0001; Bonferroni corrected for 2 tests) for the controller condition. This result replicated the within-condition session effect found in the first sample. These results suggest that the pupils’ size may slightly decrease after a few minutes in the VR headset, which could point to the participants’ familiarity with the VR environment and adaptation to the display. (Kiefer et al. 2016; Iskander et al. 2019).

#### 3.2.2 Blinks

Blink behavior was assessed as the mean number of blinks normalized by trial duration. A significant interaction was found between the condition and the session, with fewer blinks in the controller condition during the first session (*estimate* = 0.33, *t*_(2530)_ = 2.1, *p* = 0.04), corresponding to a marginal interaction observed in the first sample. No significant main effect of session was found (*estimate* = 0.1, *t*_(2532)_ = 1.1, *p* = 0.26), in contrast to the first sample, where this effect was present. In addition, no main effect of condition was observed (*estimate* = −0.35, *t*_(55)_ = −1.2, *p* = 0.23), replicating the first sample’s results.

Contrast analysis revealed a significant increase in the number of blinks from the first session to the second in the controller condition (*estimate* = −0.45, *t*_(2527)_ = −4.1, *p* < 0.0001, Bonferroni corrected for 2 tests, Fig. 7b), which was not observed in the first sample. No difference was found between sessions in the walking condition (*estimate* = −0.1, *t*_(2533)_ = −1.1, *p* = 0.26, Bonferroni corrected for 2 tests), in contrast to the first sample where such a difference was present (see Supplementary Fig. A.4.1b and Supplementary Table A.4.1 in *Appendix A.4.1*).

#### 3.2.3 Visual attention measured by gaze time

We speculated whether visual attention differed between objects in the scene across the two VR conditions. To address this, we examined the percentage of time objects were gazed at during each entire trial duration.

##### Gaze time at cue on the reward side

On average, participants gazed at the cue on the reward side for 4.6% of the trial duration. No significant main effects were observed for condition (*estimate* = −0.13, *t*_(59)_ = −0.2, *p* = 0.86) or session (*estimate* = −0.39, *t*_(1327)_ = −0.9, *p* = 0.35), replicating results from the first sample. Nevertheless, an interaction between condition and session was obtained (*estimate* = 1.69, *t*_(1324)_ = 2.9, *p* = 0.004, Fig. 8), which was not present in the first sample (see Supplementary Fig. A.4.2 and Supplementary Table A.4.2 in *Appendix A.4.1*).

**Fig. 8.**
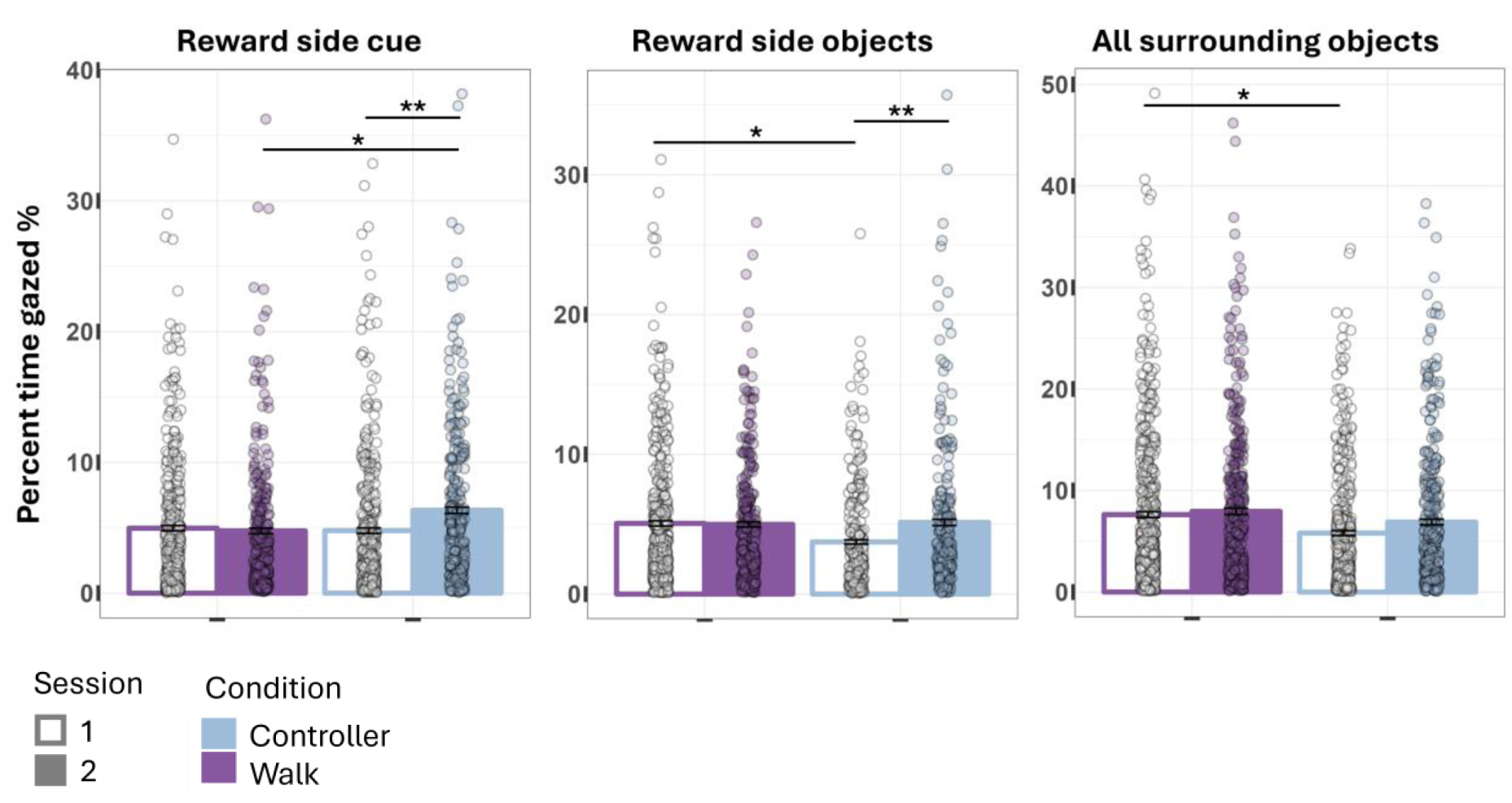
Gaze behavior measures. Bar plots of the percent time objects were gazed per trial according to VR modality of the second sample. The height of each bar presents the mean, with error bars indicating ± 1 SEM. Individual trial-based data points are shown as dots overlaid on the bars. (p < 0.001 ‘***’, p < 0.01 ‘**’, p < 0.05‘*’)

Contrast analysis showed that the proportion of gaze time on the cue in the controller condition was significantly higher in the second session compared to the first (*estimate* = −1.3, *t*_(1320)_ = −3.2, *p* = 0.0014, Bonferroni corrected for 2 tests, Fig. 8), and also significantly higher than in the walking condition in the second session (*estimate* = −1.6, *t*_(72.7)_ = −2, *p* = 0.046, Bonferroni corrected for 2 tests).

##### Gaze time at surrounding objects on the reward side

On average, participants gazed at the surrounding objects on the reward side for 4.8% of the trial duration (e.g., objects *O_1_*, and *O_2_* in Fig. 4, if the reward was on the right side). Gaze time was significantly lower by 1.5% in the controller condition relative to walking (*estimate* = −1.53, *t*_(66.6)_ = −2.4, *p* = 0.02), replicating the result from the first sample (see Supplementary Fig. A.4.2 and Supplementary Table A.4.2 in *Appendix A.4.1*). No main effect of session was obtained (*estimate* = −0.31, *t*_(1182)_ = −0.9, *p* = 0.37), unlike the significant session effect observed in the first sample. On the other hand, an interaction between condition and session was obtained in the second sample (*estimate* = 1.53, *t*_(1187_ = 2.9, *p* = 0.004), which was not present in the first.

Contrast analysis found that the percent time the objects in the reward side were gazed at significantly increased from the first session to the second in the controller condition (*estimate* = −1.22, *t*_(1190)_ = −3, *p* = 0.002, Bonferroni corrected for 2 tests, Fig. 8) the but not in the walking condition (*estimate* = 0.31, *t*_(1180)_ = 0.9, *p* = 0.37). This pattern is opposite to the one observed in the first sample. Furthermore, the percent time the objects on the reward side were gazed at, was significantly higher in the walking condition compared to the controller in the first session (*estimate* = 1.52, *t*_58.9)_ = 2.4, *p* = 0.02, Bonferroni corrected for 2 tests), but not in the second session (*estimate* = −0.005, *t*_(71.1)_ = −0.007, *p* = 0.99). In the first sample, this increase was observed in both sessions. Taken together, these results suggest that in the initial stages of the task, participants in the walking condition allocated more visual attention to surrounding objects on the reward side than those using the controller, and that this attention increased in the second session selectively in the controller condition.

##### Gaze time at all surrounding objects

Participants gazed at all the surrounding objects (objects *O_1_* – *O_5_* in Fig. 4) for 7.6% of the trial duration on average. The percentage of time all the objects were gazed at was significantly lower by 30% in the controller condition compared to the walking condition (*estimate* = −2.26, *t*_(58)_ = −2.2, *p* = 0.03, Fig. 8), replicating findings from the first sample. No difference was found between the two sessions as a main effect (*estimate* = 0.1, *t*_(1642)_ = 0.2, *p* = 0.8), or an interaction between condition and session (*estimate* = 0.64, *t*_(1645)_ = 1, *p* = 0.32, consistent with the null results in the first sample.

Contrast analysis showed that the effect of greater attention to the surrounding objects in the walking condition was significant only in the first session (*estimate* = 2.26, *t*_(55.1)_ = 2.2, *p* = 0.03), but not in the second session (*estimate* = 1.26, *t*_(60.6)_ = 1.5, *p* = 0.14, Bonferroni corrected for 2 tests, Fig. 8). This pattern replicates the first session effect but not the second, which was present in the first sample (see Supplementary Fig. A.4.2 and Supplementary Table A.4.2 in *Appendix A.4.1*). Suggesting that, the attention given to the surrounding objects as a group was higher in the first session in walking conditions compared to standing in place and using the controller.

In the exploratory analyses, marginal trends suggested that higher gaze time on all objects was associated with faster learning pace, and that lower SOT scores, but not MRT scores, were related to higher learning effectiveness in some models. These effects were marginal, and no other effects or interactions between gaze behavior and spatial learning performance were observed in the second sample, and none at all in the first (full model results are presented in Supplementary Tables A.4.3-A.4.14 in *Appendix A.4.2*).

### 3.3 Identification of individual strategy usage based on eye tracking and path features

Since differences in the ET features were observed between sessions, conditions, and samples (see section *3.2* *Eye-tracking measures*), for each category and data type, a model was constructed for each strategy, session, condition, and sample (for more details, see Tables A.5.1-A.5.10 in *Appendix A.5*).

In the attempt to predict the strategy used in each strategy test probe trial in the first sample, a trend accuracy of 0.6 was obtained for the place strategy in the second session (*p* = 0.069). Nevertheless, in the second sample, significant results were obtained for classification in the walking condition in the second session for place (*accuracy* = 0.76, *p* = 0.04), and response (*accuracy* = 0.64, *p* = 0.05) and in the first session for response using the controller (*accuracy* = 0.69, *p* = 0.02).

Predicting whether the cue strategy was the preferred one in sample 1 obtained an accuracy of 0.75 in the walking condition for the first session in sample 1 (*p* = 0.04) based on the training trials’ data, and an accuracy of 0.81 based on all trials data (*p* = 0.03). However, in the walking condition in the second sample, a significant accuracy of 0.72 was obtained for the response strategy in the second session (*p* = 0.05) when using only training trials, and an accuracy of 0.68 for the cue strategy when we used data from all the trials (*p* = 0.05).

Finally, regression for the number of explicit choices in strategy in the first sample in the walking condition provided with (*R^2^*= 0.2, *RMSE* = 1.07, *p* = 0.04) for the cue strategy in the second session when using only the training trials data, and for the response strategy when using all the data, in the first session (*R^2^* = 0.4, *RMSE* = 0.67, *p* = 0.01), and second session (*R^2^* = 0.43, *RMSE* = 0.7, *p* = 0.04). In the second sample, regression to the number of choices of cue strategy obtained (*R^2^* = 0.17, *RMSE* = 1.02, *p* = 0.04) in the first session using the controller when only training trials were used, and (*R^2^* = 0.08, *RMSE* = 1.2, *p* = 0.04) in the 2D condition, and (*R^2^* = 0.37, *RMSE* = 0.84, *p* = 0.01) for place in the walking condition both in the second session when all trials were used.

Taken together, even though the classification and regression results generally vary between sessions, strategies, experimental conditions, and samples, most of the significant models (all in sample 1) were obtained for the walking condition, and only one in the 2D condition, and most in sample 2 were for session 2.

### 3.4 Presence measures

#### 3.4.1 Spatial presence

In the powered sample, participants reported a significantly higher sense of spatial presence in the VR walking condition relative to the 2D condition (*estimate* = −28.2, *t*_(72)_ = −5.4, *p* = 8.6e-7, Fig. 9). Contrast analysis showed a significantly lower sense of spatial presence in the 2D condition relative to both VR conditions, with (*estimate* = 28.2, *t*_(72)_ = 5.38, *p* <0.0001) for the walking condition, and (*estimate* = −19.9, *t*_(72)_ = −3.8, *p* = 0.0009), for the controller condition (Bonferroni corrected for 3 tests). No significant difference was observed for the sense of spatial presence between the two VR conditions (*estimate* = −8.3, *t*_(72)_ = −1.59, *p* = 0.12). This pattern replicated preregistered findings from the first sample (see *Appendix A.2* and Supplementary Table A.2.1).

**Fig. 9.**
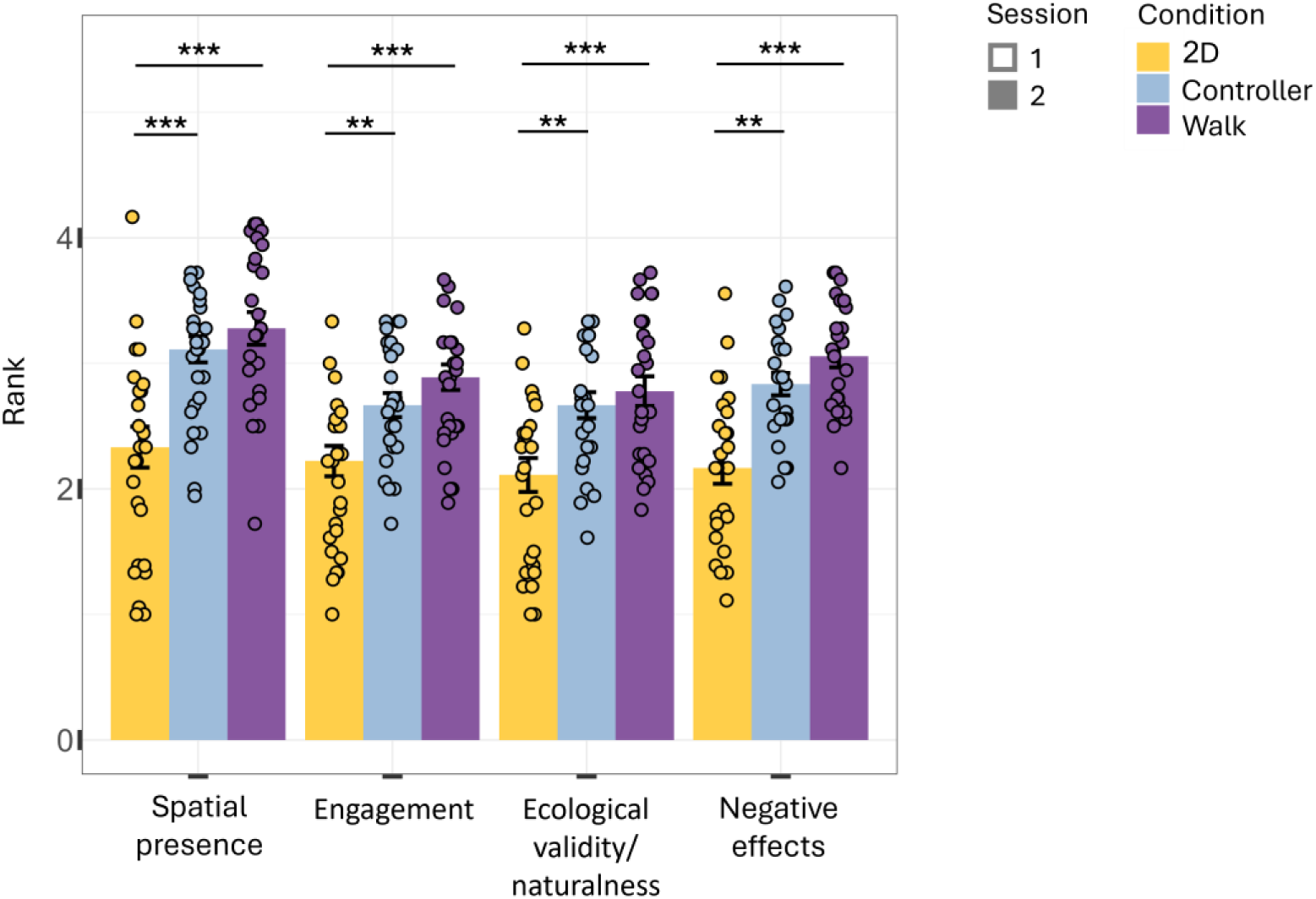
Sense of presence. Bar plots of presence factors according to VR modality of the second sample. The height of each bar presents the median, with error bars indicating ± 1 SEM. Individual data points are shown as dots overlaid on the bars. (p < 0.001 ‘***’, p < 0.01 ‘**’, p < 0.05‘*’)

#### 3.4.2 Engagement

In addition, a significantly higher sense of engagement was found in the VR walking condition relative to the 2D condition (*estimate* = −22.9, *t*_(72)_ = −4.2, *p* = 8.8e-5). Contrast analysis revealed a significantly lower sense of engagement in the 2D condition relative to both VR conditions, with (*estimate* = 22.9, *t*_(72)_ = 4.16, *p* = 0.0003) for the walking condition, and (*estimate* = −20.1, *t*_(72)_ = −3.6, *p* = 0.0015) for the controller condition (Bonferroni corrected for 3 tests, Fig. 9). Bayes factor analysis for the 2D-controller contrast showed moderate to strong evidence in favor of the alternative (BF10 = 7.1, 13, and 17.8). The difference between VR walking and 2D conditions replicated preregistered findings from the first sample, whereas the 2D-controller contrast was not observed in the initial sample and therefore was not preregistered (see *Appendix A.2* and Supplementary Table A.2.1). Eventually, no significant difference was found between the two VR conditions for this factor as well (*estimate* = −2.8, *t*_(72)_ = −0.51, *p* = 0.61), consistent with the null effect observed in the first sample.

#### 3.4.3 Ecological validity/naturalness

The ecological validity/ naturalness of the environment was found to be significantly higher in the VR walking condition relative to the 2D condition (*estimate* = −23.4, *t*_(72)_ = −4.2, *p* = 6.7e-5). Contrast analysis revealed that the ecological validity in the 2D condition was significantly lower relative to both VR conditions, with (*estimate* = −18.2, *t*_(72)_ = −3.28, *p* = 0.005) for the controller condition, and (*estimate* = 23.4, *t*_(72)_ = 4.2, *p* = 0.0002, Fig. 9, Bonferroni corrected for 3 tests), for the walking condition. These results replicate preregistered findings from the first sample. No significant difference was observed between the two VR conditions, for the ecological validity (*estimate* = −5.3, *t*_(72)_ = −0.95, *p* = 0.34), consistent with the null effect observed in the first sample.

#### 3.4.4 Negative effects

Significantly higher negative effects were obtained for the VR walking condition relative to 2D (*estimate* = −29, *t*_(72)_ = −5.6, *p* = 3.65e-7) and marginally higher than in the controller (*estimate* = −9.7, *t*_(72)_ = −1.9, *p* = 0.06). Contrast analysis found significantly lower negative effects in the 2D condition relative to both VR conditions (*estimate* = 29, *t*_(72)_ = 5.6, *p* < 0.0001) for the walking condition, and (*estimate* = −19.34, *t*_(72)_ = −3.73, *p* = 0.001) for the controller condition (Bonferroni corrected for 3 tests, Fig. 9). Bayes factor analysis indicated extreme evidence in favor of the alternative hypothesis for the 2D-walking contrast (BF10 = 1078, 2140, and 3028), and moderate to strong evidence for the 2D-controller contrast (BF10 = 8, 15.1, and 20.4). These results were not found in the first sample and were therefore not preregistered (see *Appendix A.2* and Supplementary Table A.2.1). No difference was observed for the negative effects between the two VR conditions (*estimate* = 1.2, *t*_(43)_ = 0.2, *p* = 0.81), consistent with the first sample.

### 3.5 Awareness measures

There was a significant positive relation between the perception of using the cue strategy and the number of cue-based choices (*estimate* = 0.54, *t*_(66)_ = 2.5, *p* = 0.014), replicating the effects observed in the first sample (see Supplementary Fig. A.3.1b and Supplementary Table A.3.1 in *Appendix A.3.1*).

In contrast, no relation was found between the perception of using place strategy and the actual number of place-based choices (*estimate* = 0.1, *t*_(66)_ = 0.4, *p* = 0.72), with Bayes factor analysis indicating none to moderate support for the null (BF01 = 0.8, 2.8, and 6.2). This result did not replicate the effect observed in the first sample.

No interaction was found between the perception of using the response strategy and the controller condition (*estimate* = −0.14 *t*_(66)_ = −0.48, *p* = 0.63), with Bayes factors indicating moderate support in the null (BF01 = 3,4.6, and 6); and a marginal main effect was obtained (*estimate* = 0.38, *t*_(66)_ = 1.8, *p* = 0.07). Post hoc analyses revealed a significant positive relation between perceived and actual use of the response strategy in the 2D condition (*estimate* = 0.55, *t*_(66)_ = 2.65, *p* = 0.01), marginal in the walking condition (*estimate* = 0.38, *t*_(66)_ = 1.82, *p* = 0.07), and non-significant in the controller condition (*estimate* = 0.24, *t*_(66)_ = 1.16, *p* = 0.25). Only the 2D effect replicated findings from the first sample (see Supplementary Fig. A.3.1 and Supplementary Table A.3.1 in *Appendix A.3.1*).

The awareness level of the strategy-relevant information was not related to the number of actual choices (p > 0.2 in all models for all strategies, for both main effects and interactions with the experimental conditions, consistent with findings from the first sample (see Supplementary Fig. A.3.2 and Supplementary Table A.3.2 in *Appendix A.3.1*; the models’ results are detailed in the summary tables in *Appendix A.3.2*). The participants were aware of the information of the strategies in the scene (the 99^th^ percentile of the awareness ranks was maximal (3) for all strategies and VR modalities. When considered alongside the models’ results, this implies that not choosing a strategy was not due to a lack of awareness of the information associated with it.

The summary of the relation between the perception of using strategy, and the awareness of strategy information to the number of choices linear models’ results for each effect and sample are shown in Supplementary Tables A.3.1-A.3.2, and Supplementary Fig. A.3.1 -A.3.2 in *Appendix A.3.1*.

### 3.6 Spatial abilities measures

#### 3.6.1 MRT

The MRT score was not related to the number of choices in any strategy (*estimate* = −0.09, *t*_(69)_ = −0.4, *p* = 0.68) for cue, (*estimate* = −0.08, *t*_(69)_ = −0.4, *p* = 0.68) for place, and (*estimate* = −0.08, *t*_(69)_ = −0.4, *p* = 0.68) for response. In addition, no interaction was obtained with the VR modality (*p* > 0.3 for all conditions and strategies, the models’ results are detailed in Supplementary Tables A.6.1-A.6.3, and Supplementary Fig. A.1.6a in *Appendix A.6*).

There was no difference in the distribution of the MRT score between the VR modalities (p > 0.34 for both 2D and VR using the controller conditions).

#### 3.6.2 SOT

The SOT score was not related to the number of choices in cue strategy (*estimate* = −0.16, *t*_(69)_ = −0.9, *p* = 0.39), and in place strategy (*estimate* = 0.27, *t*_(69)_ = 1.5, *p* = 0.13). Nevertheless, there was a marginally significant relation of the SOT score to the number of choices in response strategy (*estimate* = 0.33, *t*_(69)_ = 1.8, *p* = 0.07). Here as well, no interaction was observed with the VR modality (*p* > 0.17 for all conditions and strategies, the models’ results are detailed in Supplementary Tables A.6.4-A.6.6, and Supplementary Fig. A.1.6b in *Appendix A.6*).

Participants in the VR walking condition scored significantly higher in the SOT test compared to both the 2D (*estimate* = −15.5, *t*_(72)_ = −2.2, *p* = 0.03) and the VR controller conditions (*estimate* = −17.35, *t*_(72)_ = −2.5, *p* = 0.02).

When included as covariates in exploratory analysis of visual attention and learning performance, SOT but not MRT scores showed marginal relations with learning effectiveness (full model results are presented in Supplementary Tables A.4.3-A.4.14 in *Appendix A.4.2*).

### 3.7 Demographic data

#### 3.7.1 Sex

There was no difference in the sex distribution between the two samples (*estimate* = −0.09, *t*_(115)_ = −0.6, *p* = 0.53) for the sample main, or any of the interactions with the VR modality (p > 0.6 for both interaction terms).

#### 3.7.2 Age

There was also no age difference between the two samples (*estimate* = 1, *t*_(115)_ = 0.59, *p* = 0.55) for the sample main effect or any of the interactions with the VR modality (p > 0.8 for both interaction terms).

#### 3.7.3 Past VR experience

There was a marginal effect of the sample on the past VR experience (*estimate* = 0.27, *t*_(115)_ = 1.9, *p* = 0.06), and a significant interaction with the controller condition group (*estimate* = −0.48, *t*_(115)_ = −2.3, *p* = 0.02). Contrast analysis found a marginally higher number of participants with past VR experience in the second sample compared to the first in the VR walking group (*estimate* = −0.27, *t*_(115)_ = −1.9, *p* = 0.06, Bonferroni corrected for 3 tests).

#### 3.7.4 Frequency of playing computer games

There was no difference in the frequency of playing computer games between the two samples (*estimate* = 0.01, *t*_(115)_ = 0.03, *p* = 0.98) for the sample main effect, or any of the interactions with the VR modality (p > 0.5 for both interaction terms).

## 4 Discussion

This study aimed to investigate how different levels of immersion of VR modalities, their locomotion interfaces, and proprioception influence spatial learning measures, particularly strategy selection. Specifically, we aimed to (1) translate the classic T-maze task from rodents to humans and test whether it elicits the three strategies (cue, place, and response) in humans, (2) examine how VR modality and locomotion interface affect spatial learning measures, including strategy use, and subjective experience (e.g., spatial presence, perception, awareness), and (3) assess gaze behavior during navigation, its modulation by VR modality, and its relationship to strategy choice. For this purpose, to the best of our knowledge, we translated for the first time the classic T-maze task from (Barnes et al. 1980) to humans, aimed to disentangle three possible learning strategies: cue, place, and response. Using this paradigm, we compared three conditions: 3D HMD VR with physical walking, 3D HMD VR with controller-based movement, and 2D desktop VR with mouse and keyboard navigation. In addition, by leveraging the built-in eye-tracker in the two HMD VR conditions, we assessed whether VR modality level of immersion and locomotion methods influence visual attention during spatial navigation and its relationship to strategy usage. Lastly, we sought the predictive potential of ET data for strategy identification across walking and controller-based VR conditions and attempted to predict strategy usage both globally and on a trial-by-trial basis. Our results, obtained on two samples, provide several key insights into human spatial navigation, gaze behavior, and the impact of VR modalities on spatial learning.

### 4.1 Translation of the T-maze task to humans

The T-maze task was effectively adapted to humans in all VR modalities. This was demonstrated by the participants’ ability to encode navigating to the reward by reaching the task’s learning criteria (4 consecutive successful trials in the training session, like the original study). Human participants fulfilled the learning criterion nearly twice as fast as the rodents in (Barnes et al. 1980). This is of course in line with the realization that the T-maze task is indeed a rather simple task for humans relative to rodents.

Nearly 80% success was achieved in probe trials that were identical to training, which implies that the participants remembered the learned scheme during the probe phase as well. Since each test probe trial was preceded and followed by these regular probe trials, this result suggests that the participants’ choice of strategy in the test probe trials was not derived from a new encoding of the test trials design.

While our task design completely followed the original design of (Barnes et al. 1980), we adopted a different analytic approach to interpret the behavioral results in the test probe trials. The original analysis in (Barnes et al. 1980) assigned a probability score of 1 to a turn to the arm that corresponds to a single strategy, and 0.5 to each of the two strategies when the overlapping side was selected. Instead, our analysis focused on quantifying the number of explicit strategy choices, that is, trials in which participants turned to the arm corresponding uniquely to a single strategy over the other two. Notably, no participant consistently chose the arm aligned with two strategies across all probe trials; each participant made at least one clear, non-ambiguous choice of a single strategy. We believe this approach reduced the uncertainty inherent in the probe trial structure and enabled a more interpretable measure of strategy usage that is more directly aligned with the participants’ observed behavior during navigation.

In correspondence to the original study in rodents, different participants employed all place, response, and cue strategies when navigating the virtual T-maze in all VR modalities. In the original study (Barnes et al. 1980), both young and older rats showed a general preference for response strategy, with a between-group interaction, where the older rodents chose the simpler response strategy over place strategy compared to the younger rats. Similarly, our human participants also demonstrated a general tendency to use one of the S-R strategies, but differently than the rodents, they used the cue strategy more than the other two.

Across all conditions, we observed that most participants employed a mixture of place, response, and cue strategies when navigating the virtual T-maze between the two sessions and even within a session. About half of the participants who used only one strategy in each session changed it in the second session. This finding aligns with previous research demonstrating the flexibility of navigation strategies in the plus maze paradigm in rodents (Packard and McGaugh 1996) and humans (de Condappa and Wiener 2016), as well as in Tolman’s sunburst translation to humans (Doner et al. 2023).

### 4.2 Impact of VR modality on spatial learning performance

Learning performance (pace and effectiveness) showed no significant differences between the two immersive VR conditions utilizing physical walking and controller-based movement. This finding contributes to an ongoing discussion in the field of spatial cognition regarding the impact of different VR modalities and proprioception on navigation performance. On the one hand, (Huffman and Ekstrom 2019) found no significant behavioral differences in pointing accuracy, rate of learning, or spatial memory performance, suggesting that spatial retrieval tasks are modality-independent across different VR levels of body-based and joystick movement, supporting our results. On the other hand, (Ruddle et al. 2011) found that proprioceptive feedback significantly improved the accuracy of cognitive maps in VR environments. These contrasting findings suggest that the effect of VR modality on spatial learning may be task-dependent and warrant further investigation across various spatial navigation paradigms.

No significant difference in learning performance was observed between the 2D and VR walking conditions in the preregistered, powered sample (the small difference observed in the first, pilot sample, did not replicate). Bayes factor analysis showed anecdotal to no support for the corresponding hypothesis, suggesting that a larger sample size would be required for conclusive results. The results of the second sample are supported by previous research. A comparison of HMD VR and desktop versions of an 8-arm RAM found no significant differences in performance between the two modalities (Kim et al. 2018), and no differences were reported, also when three strategies were tested in a virtual sunburst maze (Doner et al. 2023).

Notably, the learning pace in the 2D condition increased significantly from the first session to the second in both samples. This suggests that participants required time to familiarize themselves with the modality, which initially affected their learning pace. This effect diminished as they became accustomed to the interface in the second session. Since we aimed to test and reveal the effect VR modality has on spatial learning characteristics, unlike most studies, our experiment began the training trials immediately, without any practice time for navigating the environment. This allowed us to explore the effect of modality habituation and naturalness of use on learning. Our results demonstrate the importance of considering learning effects when comparing different VR modalities and emphasize the need for adequate training periods in spatial navigation studies, where natural habituation wasn’t a research question, like in our case. Given that most studies typically include a practice period for participants to familiarize themselves with the navigation controls before the actual experiment begins, reporting the average amount of time it requires could contribute greatly to the field.

### 4.3 Impact of VR modality on spatial learning strategy

Examination of strategy usage across different VR modalities revealed that in both samples, there were no significant differences in the probability of using any strategy between the 2D and the VR controller conditions, and no differences between any of the modalities for the cue strategy. The powered sample showed a decrease in using cue strategy between the two sessions, which was not observed in the first sample (the VR walking condition in the pilot sample obtained other effects that were not replicated, see Appendix A). Overall, the pattern in the powered sample results corresponds to insights from (Doner et al. 2023), where no differences were reported between modalities for strategy usage, even though flexibility in strategy selection was observed within each modality.

The lack of consistent findings across samples was supported by Bayes factor analysis of the corresponding effects to various degrees, with some Bayes factors classified as anecdotal, emphasizing the need for larger double-sized participant groups to accumulate more evidence in favor of one hypothesis over the other. To better assess the practical relevance of our findings, we calculated the SMDs (Cohen’s *d*) for all reported ORs. This analysis demonstrated that all observed effects exhibited medium-to-large effect sizes (according to Cohen’s criteria), suggesting that the findings are, in fact, practically significant. Therefore, the inconsistencies across samples cannot be attributed to generally small practical effects. Additional factors likely contributed to these inconsistencies, including the small effect sizes observed, the relatively modest statistical significance of some effects (p-values exceeding 0.001), and the variability in strategy usage within participants. In addition, we posit that the limited variance of the raw measurement scale for strategy usage, specifically, the low resolution of the measure (only three probe trials for each strategy, restricting the raw score range to 0 to 3), may have constrained the variability of the underlying scores and consequently limited the stability and consistency of the effects across samples. Furthermore, the differences in instructional emphasis between samples, particularly for the 2D and controller groups in the second sample, may have further influenced the observed discrepancies. Finally, the sample size of the second group was determined by power analysis to achieve 80% power, which inherently leaves a 20% chance of failing to replicate effects even though they may exist in the population.

We additionally analyzed demographic data to assess possible factors for the variety in some of the samples’ results. There were no differences between the samples for sex, age, and frequency of playing computer games. A marginally significantly higher past VR experience was observed in the second sample compared to the first, particularly in the VR walking group. This difference in VR familiarity could potentially contribute to the variability in results between the two samples. For some participants, the experience with VR could be with gaming. Therefore, since gaming experience has been associated with improved spatial abilities and navigation performance in virtual environments (Richardson et al. 2011), these subtle demographic differences could partially explain the inconsistencies observed between the two samples in strategy usage and learning measures.

These results highlight individual variability in strategy utilization and the potential that gaze behavior analysis may provide for deeper insights into the mechanisms driving strategy usage.

### 4.4 Pupil size and blink rate

Interestingly, pupil size decreased significantly from the first session to the second across both conditions and samples. This consistent finding may reflect participants’ adaptation to the VR display over time (Kiefer et al. 2016; Iskander et al. 2019), potentially reducing cognitive load or visual strain. Previous research has shown that pupil size can be influenced by factors such as cognitive load, emotional arousal, and visual adaptation (Mathôt 2018). In our case, the decrease might reflect reduced cognitive effort as participants become more familiar with the task and environment. Such adjustments have been reported in prior spatial learning studies linking pupil size to task engagement and cognitive demands (de Condappa and Wiener 2016). However, the implications of this change remain speculative and warrant further investigation to understand its relationship to navigation strategy and VR adaptation.

One area of inconsistency was blink behavior, where results varied between samples and conditions. Blink rate has been associated with various cognitive processes, including attention, cognitive load, and fatigue (Stern et al. 1994). The inconsistency in our findings suggests that the blinks’ role in spatial navigation tasks may be influenced by factors beyond cognitive demands, such as VR-specific visual strains or individual variability in blink patterns.

### 4.5 Visual attention to maze cues

One of our primary goals was to assess whether visual attention to maze cues differed between VR modalities and sessions. Our findings revealed consistent patterns across both samples, particularly in the first session. In both samples, participants in the walking condition spent significantly more time gazing at objects on the reward side and the entire set of surrounding objects compared to those in the controller condition in the first session (and in the second session only in the pilot sample). This suggests that physical walking, being a more natural form of locomotion for humans, may allow for greater allocation of attention to environmental cues. In contrast, the need to adapt to controller-based movement may initially reduce attention to the environment. This interpretation is supported by a study (Drewes et al. 2021) that found several gaze features affected by the experimental condition of real-world navigation compared to controller-based VR, reflecting altered visual exploration due to VR constraints. Furthermore, the increased attention to reward-side objects during walking reinforces the importance of task immersion and locomotion modality in shaping gaze behavior.

Prior studies linked attentional allocation to navigation performance (Hejtmánek et al. 2018; Kapaj et al. 2024), suggesting that early visual exploration of environmental objects may predict spatial knowledge. In our exploratory analysis, a marginal trend suggested that higher gaze time on all objects was associated with a faster learning pace in the second sample.

Additional visual-attention effects were observed in the first (pilot) sample, which may reflect an interplay between behavioral strategy choice and visual attention, but these effects were not replicated (see Appendix A). This highlights the potential for individual differences in visual exploration strategies. This variability in gaze behavior may also stem from the employment of a mix of strategies, as observed by many participants. This aligns with previous findings (de Condappa and Wiener 2016), which noted mixed gaze behaviors in navigation tasks, reflecting the interplay of multiple strategies. This variability underscores the importance of using multiple ET features in our predictive models to capture the full range of gaze behaviors.

### 4.6 Prediction of navigation strategies

Our attempts to predict navigation strategies based on ET and behavioral data yielded mixed results. While classification models showed some success in identifying strategy usage, their accuracy varied across strategies, conditions, and sessions. For instance, predicting strategy choice was more successful in the walking condition, suggesting that physical locomotion may elicit more distinct gaze and behavioral patterns conducive to strategy identification.

The accuracy of our models varied depending on whether we used data from all trials, training trials only, or single trials. This variability corresponds with our finding that most participants did not rely exclusively on a single strategy but employed a mix of strategies throughout the task. As mentioned above, this finding aligns with previous research (de Condappa and Wiener 2016) that examined three possible strategies and also observed mixed gaze behavior associated with different navigation strategies. Interestingly, although the T-maze paradigm may seem as a simple task for human subjects, it elicited complex behavior.

The challenge in predicting strategies may also be due to the relatively small number of probe trials in our design compared to traditional classification tasks. Additionally, the unbalanced class distributions across some of the strategies likely influenced model performance and statistical significance of our results, even when accuracy was relatively high. Future studies could benefit from larger sample sizes and from incorporating more probe trials to obtain more data for each participant and between participants.

### 4.7 Awareness of strategy usage across VR modality

The relationship between participants’ perception of strategy usage and their actual choices during the task was significantly correlated across all conditions for the cue strategy in both samples, indicating that participants were generally aware of their reliance on this strategy. However, for the response strategy, this positive correlation was found in the 2D condition in both samples, and the VR walking condition in the second sample (effects obtained only in the first sample and not replicated are reported in Appendix A). Interestingly, in the VR conditions, participants often perceived that they used the response strategy even when their actual behavior indicated otherwise. This discrepancy suggests that participants may conflate the concept of habitual turns with more complex navigation strategies, or they may recall their intended actions rather than their actual choices during navigation. The dissociation between the perception of using a strategy and behavior, particularly for the response strategy, aligns with the idea that spatial navigation often involves implicit learning processes, as suggested by (Chrastil and Warren 2014).

The analysis of participants’ reported awareness of the information of each strategy and actual strategy implementation adds to these intriguing findings. While participants generally reported high awareness of the spatial or motoric cues associated with each strategy across modalities and samples, their explicit choices during the task were not associated with it. This implies that choices in one strategy were not derived from not noticing the other strategies.

These findings echo previous findings (Lawton 1994), where self-reported navigation strategies often aligned with spatial perception ability but were not always predictive of observed behaviors. Similarly, another study (Bray et al. 1999) noted strong correlations between self-reports and observed strategies but highlighted cases of implicit strategy use that participants failed to report. Our structured approach, where participants selected their perceived strategy from a predefined scale, contrasts with other studies (Kim et al. 2018; Gammeri et al. 2022), which utilized verbal reports.

Furthermore, the use of multiple strategies by many participants across sessions and probe trials likely contributed to the variability in the relationship between awareness, perception, and behavior. These findings emphasize the value of combining objective behavioral measures with subjective reports to capture the full complexity of spatial learning strategies, particularly in immersive and dynamic environments.

### 4.8 Presence and ecological validity

The analysis of presence factors showed significantly higher ratings of spatial presence, engagement, and ecological validity in both HMD VR conditions compared to the 2D desktop condition in both samples. This aligns with previous research demonstrating the benefits of immersive VR for creating more ecologically valid experimental environments compared to 2D displays (Kim et al. 2018). The enhanced sense of presence in HMD VR may contribute to more effective spatial learning by providing a more realistic context for encoding spatial information.

Interestingly, we did not observe significant differences in any of the presence measures between the two HMD VR conditions (walking vs. controller-based movement), also in congruence with previous findings (Kim et al. 2018). This suggests that the immersive visual experience provided by HMD VR may be more critical for creating a sense of presence than the specific locomotion interface used.

Notwithstanding, in the second sample, significantly higher negative effects were reported in both VR conditions, indicating a possible trade-off between increased ecological validity and user comfort that needs to be considered in experimental design. These cybersickness effects, including dizziness, nausea, or eyestrain, are well documented in HMD-based VR literature (Rebenitsch and Owen 2016; Weech et al. 2019; Stanney et al. 2020), and are typically linked to sensory conflicts between visual, vestibular, and proprioceptive cues. Such sensations may have been amplified by the minimalistic, floating maze environment, and could be reduced in future studies using more grounded, naturalistic, object-rich contexts or by introducing short breaks between sessions. The negative effects did not appear to influence learning performance, as learning effectiveness reached ceiling levels in all conditions, and an initial faster learning pace was found in both HMD VR conditions compared to 2D. However, they might have contributed to individual differences in spatial learning strategies or to the variability observed between the two samples.

### 4.9 Psychometric measures of spatial abilities relation to strategy

We investigated the relationship between two widely used spatial navigation abilities tests MRT and SOT and the probability for choice in strategy. These tests are frequently used to assess spatial reasoning and orientation skills, and their scores have been correlated to navigation performance in various tasks as RAM (Astur et al. 2004; Mennenga et al. 2014), MWM (Astur et al. 2004) and others (Moffat et al. 1998), and other spatial abilities scales (Huang and Voyer 2017; Friedman et al. 2020). However, in our work, we did not observe significant correlations between strategy usage and MRT or SOT scores.

A potential explanation for this discrepancy is that most participants employed multiple strategies, place, response, and cue, during the task, which could obscure direct links between specific strategies and overall spatial ability scores. This variability highlights the complexity of strategy usage in spatial tasks, aligning with prior findings that navigation involves a dynamic interplay of cognitive processes rather than reliance on a single ability (Bohbot et al. 2007; de Condappa and Wiener 2016; Doner et al. 2023). Future work should consider how mixed strategy usage moderates the relationship between standardized cognitive tests and navigation performance.

### 4.10 Limitations and future directions

While our study provides valuable insights into human spatial learning in virtual environments, several limitations should be addressed in future research. A critical consideration in interpreting the present findings lies in the inherent difference between laboratory spatial learning tasks traditionally adapted from animal models and the complex nature of real-world human wayfinding. A thorough taxonomy of wayfinding tasks (Wiener et al. 2009) highlighted that real-world navigation involves various cognitive demands that extend beyond those assessed by simplified maze-based paradigms. Specifically, real-world wayfinding incorporates dynamic environmental constraints, multitasking, widespread acquisition and application of spatial knowledge at multiple levels, and more diverse goals than typically encompassed in controlled mazes. Although the virtual T-maze paradigm effectively disentangles cue, place, and response strategies under controlled conditions, translating these discrete navigation processes to everyday wayfinding requires acknowledging the differing knowledge requirements and task structures. Consequently, while VR-based analogues of animal tasks offer valuable mechanistic insights, further research integrating more naturalistic environments and complex wayfinding scenarios is necessary to fully understand human spatial cognition in ecologically valid contexts.

The mixed results regarding the impact of VR modality on spatial learning highlight the need for more nuanced investigations of how different VR interfaces affect cognitive processes. Future studies should consider both task-specific factors and individual differences when comparing across VR modalities. Specifically, individual spatial abilities were found to be related to spatial learning performance (Hejtmánek et al. 2018; Kapaj et al. 2024). Therefore, a notable limitation of our study is the absence of a priori control for individual differences in spatial abilities. We relied on random assignment and conducted post-hoc analyses on the distribution of psychometric measures (MRT, SOT) in the second sample across conditions. Unfortunately, even though we did not find differences in MRT distributed between VR modalities, the SOT score was significantly higher in the VR walking condition compared to the two other conditions. In exploratory analyses, SOT was included as a covariate in models examining the relation between visual attention and learning performance; marginal trends suggested that lower SOT scores were associated with higher learning effectiveness in some of these models. These outcomes suggest that the group differences could have affected the spatial learning performance and the used strategy, and particularly the inconsistencies between the two samples’ outcomes. Therefore, a more robust control for this factor is necessary to fully parse the effects of VR modality, accounting for these individual differences.

Some of the inconsistencies between the samples results suggest a limited ability to detect subtle effects as reflected in the Bayes factor analysis, implying that larger-scale studies are needed to confirm and extend our findings. Additionally, the number of three test probe trials per strategy per session may have constrained the accuracy of detecting a preferred strategy based on ET and behavioral measures. Increasing the number of probe trials could enhance the reliability of these measurements and improve both the statistical significance and effect sizes.

The integration of ET into VR headsets offers significant advantages for investigating spatial learning and strategies. ET features provide direct insights into visual attention, which is crucial for understanding navigation strategies, as demonstrated by studies linking gaze dynamics to strategy shifts and environmental complexity (Walter et al., 2022; Zhu et al., 2022). However, the relatively minimalistic design of our task may have limited gaze behavior complexity. Therefore, designing more naturalistic and rich environments in future studies could enhance strategy differentiation based on ET data. Furthermore, in the present study, ET was available only in the HMD-based conditions and not in the 2D desktop condition, which limits direct comparisons of gaze behavior across modalities. Future work could address this by incorporating desktop-based ET to allow a more comprehensive assessment of visual exploration across different VR modalities.

Our findings contribute to the ongoing discussion about the impact of VR modality on spatial cognition. Even though some studies have suggested that the neural correlates of spatial retrieval are modality-independent (Huffman and Ekstrom 2019), others have found that proprioceptive feedback can improve cognitive map accuracy (Ruddle et al. 2011). Our results suggest that the effect of VR modality on spatial learning may be task-dependent and individual and warrants further investigation.

Utilizing ET data for predictive models underscores its potential for identifying individual navigation strategies. However, the variability in gaze and behavioral patterns across trials and conditions presents challenges inherent in such endeavors. In addition to additional probe trials, and richer VR environments suggested above, future studies should incorporate more sophisticated predictive models that account for extremely unbalanced class distribution. Likewise, combining ET data with other physiological measures (e.g., heart rate variability, and skin conductance) could assist in improving strategy prediction and deepen our understanding of the interplay between navigation behavior and physiological processes.

An additional future direction involves uncovering the neural substrates of spatial navigation strategies in the three-strategy T-maze task. Animal models have shown that spatial strategies engage distinct brain regions, with the hippocampus playing a key role in allocentric (place) strategies and the striatum supporting egocentric (response) strategies (Packard and McGaugh 1996). Subsequent human studies have provided evidence of similar neural mechanisms, demonstrating plasticity-related structural changes in London taxi drivers (Maguire et al. 2000, 2006), fMRI activations (Iaria et al. 2003) and voxel-based morphometry (VBM) analysis (Bohbot et al. 2007) in RAM adaptations, and functional connectivity in virtual MWM (Woolley et al. 2015). These findings highlight the value of translational research in bridging animal and human models of spatial navigation. Given that our study found no differences in learning effectiveness between 2D and 3D HMD VR displays after participants adapted to the interface, nor consistent differences in strategy use, employing fMRI to examine the T-maze task could offer novel insights into its neural correlates. Studying the neural mechanisms underlying the three-strategy paradigm through neuroimaging techniques would further enhance our understanding of spatial strategy selection and execution.

Our findings have potential implications for understanding and treating neurodegenerative and psychiatric conditions affecting spatial cognition. Such distinctions have been related to old age (Barnes et al. 1980; Marighetto et al. 1999; Etchamendy et al. 2012; Wiener et al. 2013) and observed in conditions like amnestic mild cognitive impairment (aMCI) (Weniger et al. 2011), Parkinson’s disease (PD) (Schneider et al. 2017), obsessive-compulsive disorder (OCD) (Marsh et al. 2015), stress (Schwabe et al. 2007), Alzheimer’s disease (AD) (Lithfous et al. 2013), Attention deficit hyperactivity disorder (ADHD) (Robaey et al. 2016), and cocaine dependence (Tau et al. 2014). Our task’s ability to disentangle the strategies involved in these studies underlines its potential to provide a sensitive tool for detecting subtle cognitive changes in these conditions. Using HMD VR-based tasks in clinical assessments offers additional advantages, as the immersive nature of VR can enhance patient engagement and motivation during evaluation procedures (Pieri et al. 2023). Importantly, by manipulating environmental features, patients could be guided toward using specific navigation strategies in our VR-based T-maze, providing a targeted approach for rehabilitating, allowing clinicians to tailor interventions to enhance particular cognitive functions or compensate for specific deficits, and foster neural plasticity.

Eventually, examining how spatial learning strategies and performance evolve over extended periods could provide insights into the potential of VR for long-term cognitive training. Future studies should explore the long-term retention of spatial knowledge acquired in different VR modalities and investigate how learning performance and the use of strategy may change over extended periods of practice.

### 4.11 Conclusions

This study provides novel insights into how VR modality level of immersion, locomotion interfaces, and proprioception influence spatial learning, strategy selection, and gaze behavior. Our successful translation of the classic T-maze task from (Barnes et al. 1980) that disentangles cue, place, and response strategies to human participants bridged the gap between controlled laboratory studies in animals and real-world human navigation research. We demonstrated that human navigation strategies mirror those observed in animal models while also exhibiting distinct flexibility in strategy utilization across VR conditions. The results indicated no significant differences in learning performance (pace and effectiveness) between the two HMD VR conditions (walking vs. controller). Nevertheless, the study highlighted the importance of considering modality habituation effects, particularly in 2D desktop VR, where learning pace increased significantly from the first session to the second, suggesting participants required time to adjust the navigation in that interface. This effect was supported by the participants’ reports of significantly lower presence and naturalness in the 2D condition relative to the HMD VR conditions. Our findings contribute to the ongoing debate on the role of VR immersion in spatial cognition and its related gaze behavior by revealing that the accuracy of strategy prediction models based on gaze measures varied substantially depending on the data we used (all trials, training trials, or single trials), condition, and session number. This variability corresponds with the behavioral mixture of strategies employed by most participants, highlighting the importance of considering task dependency and individual differences, and habituation effects. The analysis of ET data offered insights into visual attention allocation by demonstrating that physical walking significantly increased visual attention allocation to environmental cues (reward-side objects and surrounding objects) compared to controller-based movement, particularly during the first session, thereby suggesting that physical walking may enhance attention to environmental cues. Overall, our translation of the T-maze task to various VR modalities contributes to a more nuanced understanding of human spatial cognition and paves the way for the development of VR-based tools for both basic research and clinical applications.

## Funding resources

This study was supported by the Minducate Science of Learning Research and Innovation Center of the Sagol School of Neuroscience, Tel Aviv University, and the European Research Council (ERC) under the European Union’s Horizon Europe research and innovation programme (grant number 101087869). The funders had no role in the study design, data collection and analysis, decision to publish, or manuscript preparation.

## Acknowledgments

The authors would like to acknowledge the Minerva Center for Human Intelligence in immersive, augmented and mixed Realities at Tel Aviv University.

We thank Harry Heiberger and Hanry Heiberger for their help in adapting the eye-tracking code from Tobii’s SDK to the SRanipal.

## Author contributions

The authors confirm their contribution to the paper as follows:

Michal Gabay: Conceptualization, Data curation, Formal analysis, Funding acquisition, Investigation, Methodology, Project administration, Software, Visualization, Writing – original draft, Writing – review and editing.

Tom Schonberg: Conceptualization, Funding acquisition, Methodology, Resources, Supervision, Validation, Writing – review and editing.

## Declarations

### Data availability

All the data and the codes for data handling, pre-processing, analysis, and Unity task projects are available in the Open Science Framework (OSF, project page: https://osf.io/arfy9/).

### Compliance with Ethical Standards

This study involved human participants and was approved by the ethics committee of Tel-Aviv University. All participants provided their written informed consent prior to the experiment.

