## Supplementary material for "The effect of virtual reality modality level of immersion and locomotion on spatial learning and gaze measures"

Appendix A - Supplementary material

Unless stated otherwise, the analyses presented in sections A.1-A.4 correspond to the pilot dataset (the first sample). These results were used to identify candidate effects and guide the design of the preregistered powered analysis for the second sample. While these results are reported here for transparency, they were not used for inferential conclusions and should be interpreted as exploratory.

### A.1 Learning measures

**Learning pace**

In the first sample, the learning pace in the 2D condition obtained a median of 10 trials to reach the learning criterion, which was significantly slower relative to a median of 6 trials in the VR walking condition (*estimate* = 23.3, $t_{(85.9)}$ = 2.6, $p$ = 0.009, see Supplementary Fig. A.1.1 and Supplementary Table A.1.1). Bayes factor analysis provided anecdotal evidence in favor of the alternative hypothesis (BF10 = 1.8, 2,2, and 2.7 for the different prior scales).

Contrast analysis showed that during the first session, there was a significant difference in the learning pace between the 2D and VR walking conditions (*estimate* = -23.4, $t_{(85.9)}$ = -2.6, $p$ = 0.027, Bonferroni corrected for 3 tests). This result was supported by anecdotal to moderate evidence with Bayes factor analysis (BF10 = 2.3, 3.3, and 4.1).

Another contrast indicated that the participants' learning pace was significantly faster in the second session compared to the first one in the 2D condition (*estimate* = 25.3, $t_{(43)}$ = 2.8, $p$ = 0.007, Bonferroni corrected for 3 tests).

Nevertheless, a main effect for the second session was not obtained (*estimate* = -14, $t_{(43)}$ = -1.6, $p$ = 0.11), nor any of its interactions with the other experimental conditions (*estimate* = -11.3, $t_{(43)}$ = -0.9, $p$ = 0.36) for Condition2D x Session2, and (*estimate* = 5.1, $t_{(43)}$ = 0.4, $p$ = 0.67) for ConditionController x Session2. Finally, there was no difference in the learning pace between the two VR conditions (*estimate* = 4.2, $t_{(85.9)}$ = 0.5, $p$ = 0.63).

**Learning effectiveness**

In the first sample, in the VR walking condition, participants obtained a median of maximum successful probe trials that were identical to training (9) which was significantly higher by 22.2% (2 trials) compared to 2D (*estimate* =-17.7, $t_{(85.4)}$ = -2.2, $p$ = 0.033, see Supplementary Fig. A.1.1 and Supplementary Table B.1.1). Bayes factor analysis provided moderate support for the alternative hypothesis over the null (BF10 = 4, 6.8, and 8.7 for the different prior scales).

Contrast analysis showed that during the second session, there was a significant difference in the learning effectiveness between the 2D and VR while walking conditions in the first sample (*estimate* =21.8, $t_{(85.4)}$ = -2.2, $p$ = 0.03, Bonferroni corrected for 3 tests). Bayes factor analysis obtained anecdotal evidence for the alternative hypothesis over the null (BF10 = 2.2, 1.8, and 2.3).

There was no significant difference in the learning effectiveness between the two VR conditions as assessed by the main effect of the controller condition (*estimate* = -12.62, $t_{(85.4)}$ = -1.52, $p$ = 0.13). In addition, there was no significant difference in the learning effectiveness between the two sessions, as a main effect (*estimate* = 4.75, $t_{(43)}$ = 0.61, $p$ = 0.55) or as an interaction with the controller (*estimate* = 9.31, $t_{(43)}$ = 0.83, $p$ = 0.41) or the 2D conditions (*estimate* = -3.88, $t_{(43)}$ = -0.35, $p$ = 0.73).

**Supplementary Table A.1.1:** Learning measures summary of results.

| Effect | First sample | Second sample | Replication |
|---|---|---|---|
| Pace (VR Walk vs 2D main effect) | + | - | Not replicated |
| Pace (VR Walk vs Controller main effect) | - | - | Replicated |
| Pace (Session1 vs 2 main effect) | - | - | Replicated |
| Pace (Session x 2D interaction) | - | - | Replicated |
| Pace (Session x Controller interaction) | - | - | Replicated |
| Pace (Session 1 VR Walk vs 2D contrast) | + | - | Not replicated |
| Pace (Session 1 vs 2, 2D contrast) | + | + | Replicated |
| Effectiveness (VR Walk vs 2D main effect) | + | - | Not replicated |
| Effectiveness (VR Walk vs Controller main effect) | - | - | Replicated |
| Effectiveness (Session 1 vs 2 main effect) | - | - | Replicated |
| Effectiveness (Session x 2D interaction) | - | - | Replicated |
| Effectiveness (Session x Controller interaction) | - | - | Replicated |
| Effectiveness (Session 2 VR Walk vs 2D contrast) | + | - | Not replicated |

Summary of the learning measures mixed linear models' results for each effect, and sample. (where '+': significant effect; '.' Marginally significant effect; '-': non-significant effect.

##### Learning strategy

Most participants used a combination of strategies across the different probe trials and sessions.

*Cue strategy:* There were no significant effects of any fixed factors or interactions in the pilot sample for the probability of using cue strategy (OR = 0.45, 95% CI = [0.09, 2.16], $p = 0.32$, $d = -0.44$) for the 2D condition, and (OR = 0.58, 95% CI = [0.12, 2.83], $p = 0.5$) for the controller condition (Supplementary Fig. A.1.2).

In addition, no effects were observed for the session main effect (OR = 1, 95% CI = [0.38, 2.62], $p = 1$, , $d = 0$), as well as for the Condition2D x Session2 (OR = 1.3, 95% CI = [0.32, 5.13], $p = 0.71$, $d = -0.63$), and the ConditionController x Session2 interaction(OR = 1.78, 95% CI = [0.43, 7.27], $p = 0.74$, $d = 0.32$). Bayes factor analysis showed strong to extreme evidence for the null for the session main effect (BF01 = 15.7, 56.2, and 3157.5), anecdotal to strong evidence for the null for the interaction with the 2D condition (BF01 = 2.3, 3.5, and 13.4), and anecdotal to moderate evidence for the null for the interaction with the controller condition (BF01 = 1.8, 2.5, and 9.3).

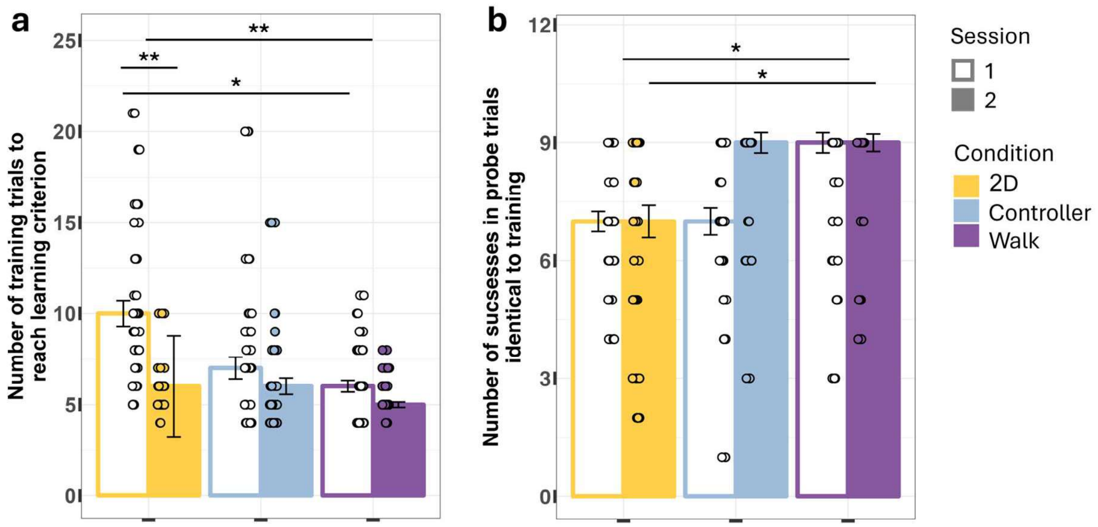


**Supplementary Fig. A.1.1.** Learning measures. Bar plots of learning measures of the first sample according to VR modality and session. Since the models were performed on the measures' ranks, the height of each bar presents the median, with error bars indicating ± 1 SEM (standard error of the mean). Individual data points are shown as dots overlaid on the bars, with y axis limit of mean ±2 SD (standard deviations). ($p < 0.01$ '**', $p < 0.05$'*').

*Place strategy:* In the model for the probability of an explicit choice of place strategy, a significant main effect of the second session was obtained in the first sample (OR = 3.84, 95% CI [1.38, 10.67], $p = 0.009$, $d = 0.74$) as well as a significant interaction between session 2 and the VR using the controller condition relative to the model's default which was session 1 and the VR walking condition (OR = 0.11, 95% CI [0.02, 0.53], $p = 0.006$, $d = -1.22$). Bayes factor analysis provided anecdotal to no support for the alternative model relative to the null for the second session main effect (BF10 = 2.5, 1.1, and 0 across priors scales), and strong to moderate evidence for the alternative hypothesis relative to the null for the interaction (BF10 = 13.1, 15.9, and 6.5).

Contrast analysis revealed that the probability of using place strategy significantly increased in the second session relative to the first in the VR walking condition in the first sample (OR = 0.26, 95% CI [0.09, 0.72], p = 0.0098, , *d* = -0.74, Supplementary Fig. A.1.2, Bonferroni corrected for 3 tests). In addition, during the second session, the probability of using the place strategy was higher in the VR walking condition relative to VR using the controller condition (OR = 8.5, 95% CI [1.33, 54.1], *p* = 0.016, *d* = 1.18). Bayes factor analysis provided support for the alternative model, with moderate to strong evidence for the increase in the walking condition between the two sessions (BF10 = 11.7, 10, and 3.1) and anecdotal to moderate evidence for the difference between the two VR conditions in the second session (BF10 = 1.7, 2.9, and 3.2).

No main effects were found for the 2D or controller conditions, nor was a significant interaction observed between the second session and the 2D condition (OR = 1.42, 95% CI [0.34, 5.86], *p* = 0.63, *d* = 0.19) for the 2D condition main effect; (OR = 1.03, 95%CI [0.24, 4.37], *p* = 0.96, *d* = 0.02) for the controller condition main effect; and (OR = 0.43, 95% CI [0.1, 1.73], *p* = 0.23, *d* = -0.47) for the 2D interaction with session 2.

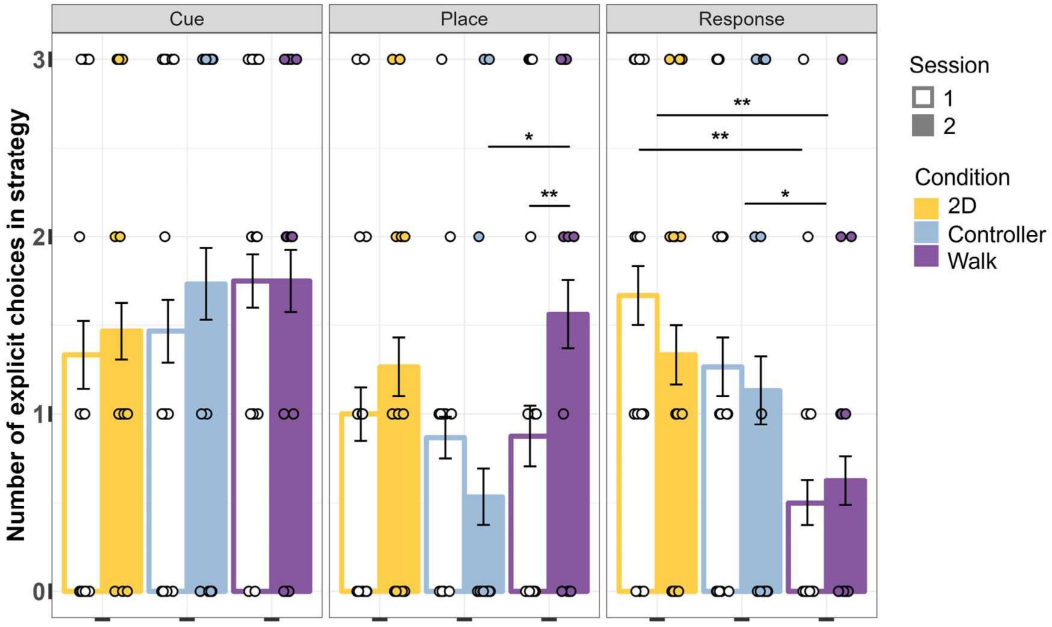


**Supplementary Fig. A.1.2:** Choice of strategy. Choice of strategy. Bar plots of the number of explicit choices in each strategy during strategy test probe trials of the first sample according to VR modality and session. The height of each bar presents the mean, with error bars indicating ± 1 SEM. Individual data points are shown as dots overlaid on the bars. ($p < 0.001$ '***', $p < 0.01$ '**', $p < 0.05$'*').

*Response strategy:* The probability for an explicit choice of response strategy (turn left/right) was significantly lower in the VR walking condition relative to both other conditions only in the first sample (OR = 4.92, 95% CI [1.24, 19.39], *p* = 0.03, *d* = 0.88) and (OR = 9.63, 95%

CI [2.42, 38.3], $p$ = 0.001, $d$ = 1.25) for the VR controller and the 2D conditions respectively, Supplementary Fig. A.1.2). Bayes factor analysis provided strong to moderate support in the alternative hypothesis (BF10 = 21.4, 19.6, and 6.3 for the different priors SD) for the 2D condition, and anecdotal to no support in the alternative model for the controller condition (BF10 = 2.9, 2.3, and 0.9).

Contrast analysis showed that during the first session, the lower probability for using response strategy in the walking condition relative to 2D was significant (OR = 0.1, 95% CI [0.02, 0.56], $p$ = 0.004, $d$ = -1.27, Bonferroni corrected for 3 tests) and it was marginal compared to using the controller (OR = 0.2, 95% CI [0.04, 1.1], $p$ = 0.07, $d$ = -0.89). Bayes factor analysis showed moderate to strong evidence in favor of the alternative (BF10 = 9.7, 14.5, and 7.5) for the difference between the 2D and VR walking conditions in the first session.

The session did not obtain a significant effect on the probability for using response strategy (OR = 1.37, 95% CI [0.45, 4.16], $p$ = 0.57, $d$ = 0.17) for the session main effect (OR = 0.41, 95% CI [0.13, 2.48], $p$ = 0.23, $d$ = -0.49); for the ConditionController x Session2 interaction; and (OR = 1.37, 95% CI [0.45, 4.16], $p$ = 0.57, $d$ = 0.17) for the Condition2D x Session2 interaction.

**Supplementary Table A.1.2:** Choice of strategy summary of results.

| **Effect** | **First sample** | **Second sample** | **Replication** |
|---|---|---|---|
| Cue (VR Walk vs 2D main effect) | - | - | Replicated |
| Cue (VR Walk vs Controller main effect) | - | - | Replicated |
| Cue (Session 1 vs 2 main effect) | - | + | Not replicated |
| Cue (Session x 2D interaction) | - | + | Not replicated |
| Cue (Session x Controller interaction) | - | + | Not replicated |
| Cue (Session 1 vs 2 VR Walk contrast) | - | + | Not replicated |
| Place (VR Walk vs 2D main effect) | - | - | Replicated |
| Place (VR Walk vs Controller main effect) | - | - | Replicated |
| Place (Session 1 vs 2 main effect) | + | - | Not replicated |
| Place (Session x 2D interaction) | - | - | Replicated |
| Place (Session x Controller interaction) | + | - | Not replicated |
| Place (Session 1 vs 2, VR Walk contrast) | + | - | Not replicated |
| Place (Session 2 VR Walk vs Controller contrast) | + | - | Not replicated |
| Response (VR Walk vs 2D main effect) | + | - | Not replicated |
| Response (VR Walk vs Controller main effect) | + | - | Not replicated |
| Response (Session 1 vs 2 main effect) | - | - | Replicated |
| Response (Session x 2D interaction) | - | - | Replicated |
| Response (Session x Controller interaction) | - | - | Replicated |
| Response (Session 1 VR Walk vs 2D contrast) | + | - | Not replicated |

Summary of the choice of strategy generalized mixed linear models' results for each effect, and sample. (where '+': significant effect; '.' Marginally significant effect; '-': non-significant effect.

#### A.2 Presence measures

In the first sample, a significantly higher sense of spatial presence was obtained in the VR walking condition relative to the 2D condition (*estimate* = -13.4, $t_{(43)}$ = -3, $p$ = 0.004, Supplementary Fig. B.2.1). Contrast analysis showed a significantly lower sense of spatial presence in the 2D condition relative to both VR conditions (*estimate* = 13.4, $t_{(43)}$ = 3, $p$ = 0.01, and *estimate* = -13, $t_{(43)}$ = -2.9, $p$ = 0.02 for walking and controller conditions, respectively, Bonferroni corrected for 3 tests). There was no significant difference in the sense of spatial presence between the two VR conditions (*estimate* = -0.3, $t_{(43)}$ = -0.07, $p$ = 0.94).

In addition, a significantly higher sense of engagement was found in the VR walking condition relative to the 2D condition (*estimate* = -10.6, $t_{(43)}$ = -2.3, $p$ = 0.03). Contrast analysis showed a marginally higher sense of engagement in the VR walking condition relative to the 2D condition (*estimate* = 10.6, $t_{(43)}$ = 2.3, $p$ = 0.08, Bonferroni corrected for 3 tests). No difference was observed for the sense of engagement in the 2D condition relative to both VR conditions (*estimate* = 10.6, $t_{(43)}$ = 2.3, $p$ = 0.08) for the walking and (*estimate* = -6.3, $t_{(43)}$ = -1.3, $p$ = 0.56) for the controller conditions, with equal support for both hypotheses (BF01 = 1 for all prior scales). Finally, no significant difference was found between the two VR conditions for this factor as well (*estimate* = -4.3, $t_{(43)}$ = -0.9, $p$ = 0.36).

The ecological validity/ naturalness of the environment was also found to be significantly higher in the VR walking condition relative to the 2D condition (*estimate* = -10.1, $t_{(43)}$ = -2.2, $p$ = 0.03). Contrast analysis revealed that the sense of ecological validity in the 2D condition was significantly lower relative to the VR controller condition (*estimate* = -12.4, $t_{(43)}$ = -2.7, $p$ = 0.03), and marginally lower relative to the VR walking condition (*estimate* = 10.1, $t_{(43)}$ = 2.2, $p$ = 0.09, Bonferroni corrected for 3 tests). No significant difference was observed between the two VR conditions, also for the ecological validity (*estimate* = -2.3, $t_{(43)}$ = 0.5, $p$ = 0.61).

No significant effects were found for the negative effects factor in the first sample (*estimate* = -5.3, $t_{(43)}$ = -1.1, $p$ = 0.27, *estimate* = 5.3, $t_{(43)}$ = 1.1, $p$ = 0.82, *estimate* = -6.5, $t_{(43)}$ = -1.3, $p$ =0.57, for the 2D - walking main effect, 2D - walking contrast, and 2D - controller contrast respectively), with none to anecdotal support in the null (BF01 = 1, 1.1, and 1.1 for both contrasts). In addition, there was no difference in the negative effects between the two VR conditions in the first sample (*estimate* = 1.2, $t_{(43)}$ = 0.2, $p$ = 0.81).

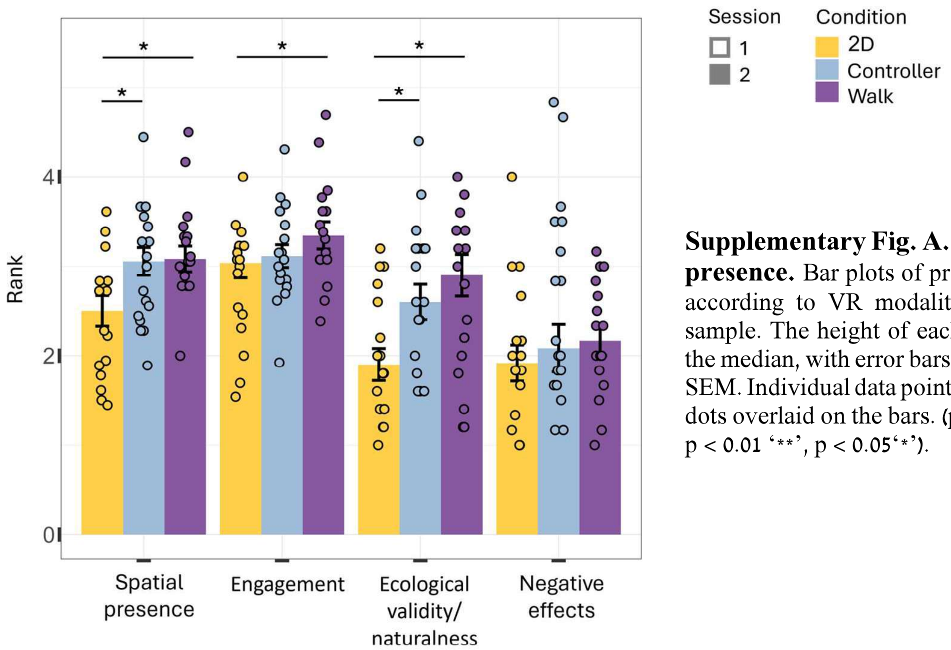


**Supplementary Fig. A.2.1: Sense of presence.** Bar plots of presence factors according to VR modality of the first sample. The height of each bar presents the median, with error bars indicating ± 1 SEM. Individual data points are shown as dots overlaid on the bars. ($p < 0.001$ '***', $p < 0.01$ '**', $p < 0.05$'*').

**Supplementary Table A.2.1:** Presence questionnaire summary of results.

| Effect | First sample | Second sample | Replication |
|---|---|---|---|
| Spatial presence (VR Walk vs 2D main effect) | + | + | Replicated |
| Spatial presence (VR Walk vs Controller main effect) | - | - | Replicated |
| Spatial presence (VR Walk vs 2D contrast) | + | + | Replicated |
| Spatial presence (Controller vs 2D contrast) | + | + | Replicated |
| Engagement (VR Walk vs 2D main effect) | + | + | Replicated |
| Engagement (VR Walk vs Controller main effect) | - | - | Replicated |
| Engagement (VR Walk vs 2D contrast) | . | + | Mid replicated |
| Engagement (Controller vs 2D contrast) | - | + | Not replicated |
| Ecological validity (VR Walk vs 2D main effect) | + | + | Replicated |
| Ecological validity (VR Walk vs Controller main effect) | - | - | Replicated |
| Ecological validity (VR Walk vs 2D contrast) | . | + | Mid replicated |
| Ecological validity (Controller vs 2D contrast) | + | + | Replicated |
| Negative effects (VR Walk vs 2D main effect) | - | + | Not replicated |

Summary of the presence questionnaire factors linear models results for each effect, and sample. (where '+': significant effect; '.' Marginally significant effect; '-': non-significant effect.

### A.3 Awareness measures

#### A.3.1 Main results

There was a significant positive relation between the perception of using the cue strategy and the number of choices of it (*estimate* = 0.6, $t_{(38)} = 2.7$, $p = 0.009$) for the perception of strategy main effect (Supplementary Fig. A.3.1a).

A significant positive relation between the perception of using place strategy and the actual number of choices of it was obtained (*estimate* = 0.65, $t_{(38)} = 2.5$, $p = 0.018$). Bayes factor analysis indicated strong to anecdotal evidence for the alternative hypothesis (BF10 = 13.6, 4.3, and 2).

An interaction was found between the perception of using the response strategy and the controller condition (*estimate* = 0.75, $t_{(38)} = 2.3$, $p = 0.03$) with moderate to no evidence for the alternative (BF10 = 2.6, 1.4, and 1). Although this effect was not preregistered for each condition separately, post hoc analyses were conducted due to the observed interaction. Perception of using response strategy predicted the actual use of that strategy in both the controller and 2D conditions, with a slightly stronger effect for the controller (*estimate* = 0.89, $t_{(38)} = 3.8$, $p < 0.001$) than the 2D (*estimate* = 0.65, $t_{(38)} = 3.1$, $p = 0.003$), but not in the walking condition (*estimate* = 0.14, $t_{(38)} = 0.63$, $p = 0.53$).

The awareness level of the strategies' information was not related to the actual choices of these strategies (p > 0.2 in all models for all strategies for main effects and interactions with the experimental conditions, Supplementary Fig. A.3.2a and Supplementary Table A.3.2; the models' results are detailed in the summary tables in *Appendix A.3.2*).

**Supplementary Table A.3.1:** Relation between perception of using strategy and number of choices, summary of results.

| Effect | First sample | Second sample | Replication |
|---|---|---|---|
| Cue (main effect) | + | + | Replicated |
| Place (main effect) | + | - | Not replicated |
| Response (main effect) | - | . | Mid replicated |
| Response (Controller interaction) | + | - | Not replicated |

Summary of the relation between perception of using strategy and number of choices linear models' results for each effect, and sample. (where '+': significant effect; '.' Marginally significant effect; '-': non-significant effect.

**Supplementary Table A.3.2:** Relation between awareness of strategy-relevant information and number of choices, summary of results.

| Effect | First sample | Second sample | Replication |
|---|---|---|---|
| Cue (main effect) | - | - | Replicated |
| Place (main effect) | - | - | Replicated |
| Response (main effect) | - | - | Replicated |

Summary of the relation between awareness of the strategy-relevant information in the scene and number of choices, linear models' results for each effect, and sample. (where '+': significant effect; '.' Marginally significant effect; '-': non-significant effect.

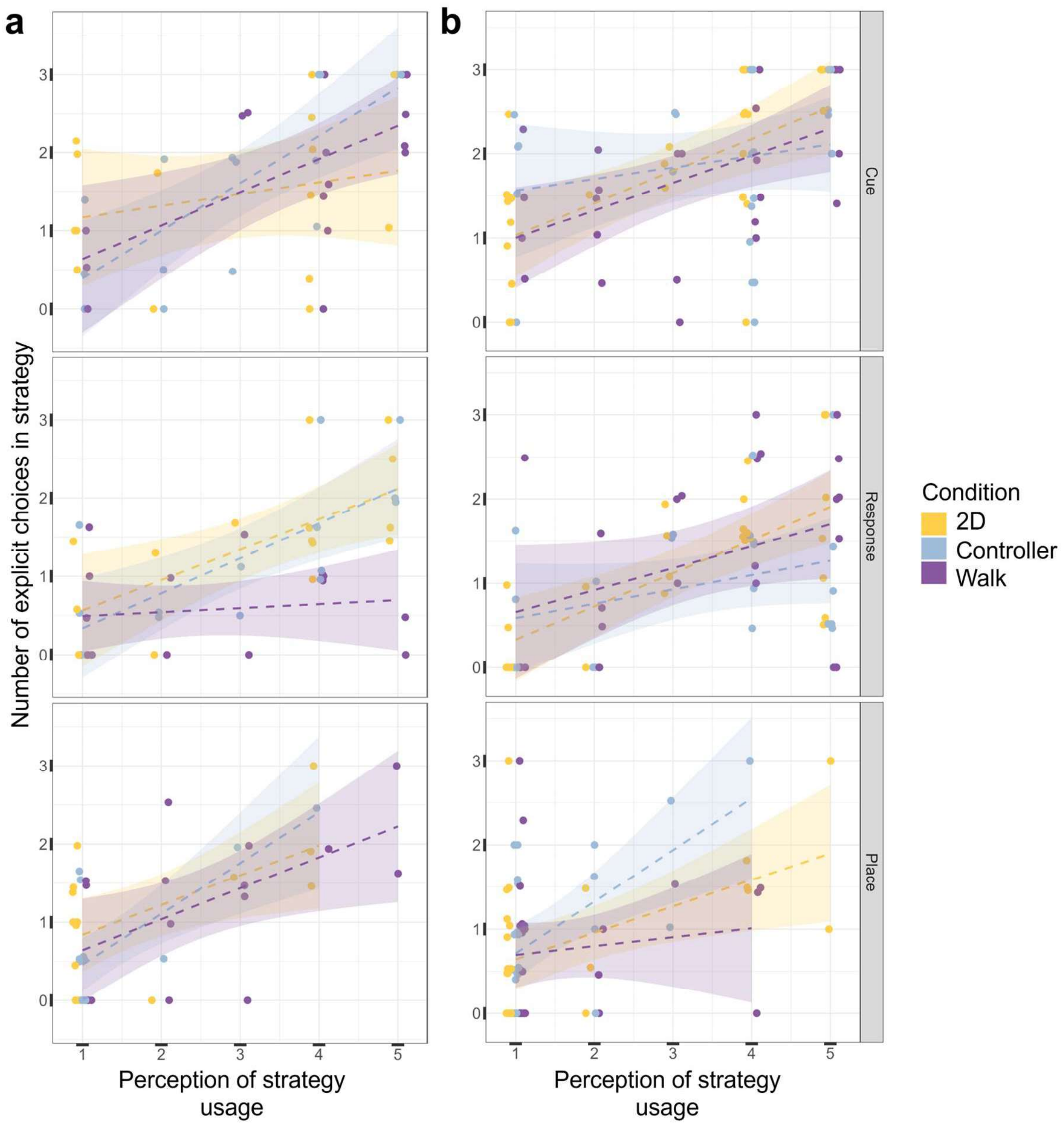


**Fig. A.3.1: Perception of using strategy.** Scatter plots of the mean across sessions of the number of explicit choices in strategy, in each strategy test probe trial, as a function of the reported perception of using the strategy. Individual data points are shown as dots slightly jittered around the x-axis for display purposes. The dashed lines represent the linear models fit for each strategy and each VR modality (fitted lines are surrounded by polygons of the 95% confidence interval). **a:** The first sample's results. **b:** The second sample's results.

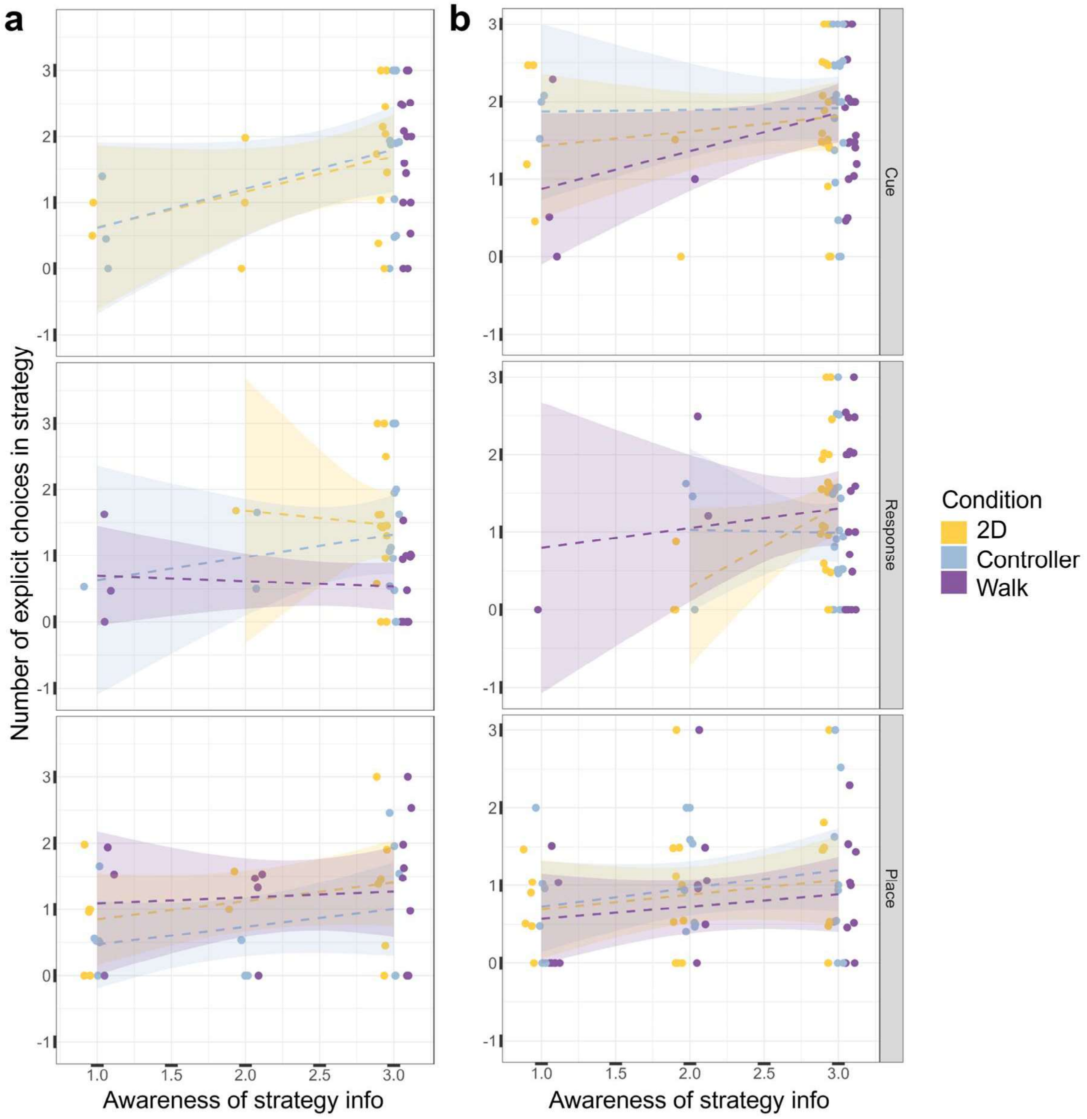


**Fig. A.3.2: Awareness of strategy information.** Scatter plots of the mean across sessions of the number of explicit choices in strategy in each strategy test probe trial, as a function of the reported awareness of the strategy information in the scene. Individual data points are shown as dots slightly jittered around the x-axis for display purposes. The dashed lines represent the linear models fit for each strategy and each VR modality (fitted lines are surrounded by polygons of the 95% confidence interval). **a:** The first sample's results. **b:** The second sample's results.

#### A.3.2: Detailed statistics tables

**Supplementary Table A.3.3:** Choices of cue strategy in the first sample

| | rank(N_Explicit_choices) | | |
|---|---|---|---|
| *Predictors* | *Estimates* | *CI* | *p* |
| (Intercept) | 13.93 | -11.61 – 39.47 | 0.276 |
| rank(Awareness level) | -0.17 | -0.96 – 0.62 | 0.673 |
| rank(Perception using strategy level) | 0.60 | 0.16 – 1.04 | **0.009** |
| Condition [2D] | -2.61 | -31.31 – 26.09 | 0.855 |
| Condition [Controller] | -9.50 | -31.97 – 12.97 | 0.397 |
| rank(Awareness level) x Condition [2D] | 0.66 | -0.38 – 1.71 | 0.208 |
| rank(Perception using strategy level) x Condition [2D] | -0.58 | -1.29 – 0.12 | 0.103 |
| rank(Perception using strategy level) x Condition [Controller] | 0.44 | -0.36 – 1.24 | 0.273 |
| Observations | 46 | | |
| $R^2$ / $R^2$ adjusted | 0.415 / 0.307 | | |

Results of the linear model on ranks for the mean number of explicit choices in cue strategy as a function of perception of using this strategy, awareness of the strategy information, and the VR modality in the first sample.

**Supplementary Table A.3.4:** Choices of cue strategy in the second sample

| | **rank(N_Explicit_choices)** | | |
|---|---|---|---|
| *Predictors* | *Estimates* | *CI* | *p* |
| (Intercept) | 11.94 | -11.50 – 35.38 | 0.313 |
| rank(Awareness level) | 0.08 | -0.56 – 0.73 | 0.798 |
| rank(Perception using strategy level) | 0.54 | 0.11 – 0.97 | **0.014** |
| Condition [2D] | 5.79 | -24.88 – 36.46 | 0.708 |
| Condition [Controller] | 23.55 | -12.03 – 59.12 | 0.191 |
| rank(Awareness level) x Condition [2D] | -0.26 | -1.11 – 0.59 | 0.544 |
| rank(Awareness level) x Condition [Controller] | -0.19 | -1.12 – 0.73 | 0.679 |
| rank(Perception using strategy level) x Condition [2D] | 0.23 | -0.37 – 0.83 | 0.446 |
| rank(Perception using strategy level) x Condition [Controller] | -0.30 | -0.90 – 0.30 | 0.317 |
| Observations | 75 | | |
| $R^2$ / $R^2$ adjusted | 0.285 / 0.199 | | |

Results of the linear model on ranks for the mean number of explicit choices in cue strategy as a function of perception of using this strategy, awareness of the strategy information, and the VR modality in the second sample.

**Supplementary Table A.3.5:** Choices of response strategy in the first sample

| | rank(N_Explicit_choices) | | |
|---|---|---|---|
| *Predictors* | *Estimates* | *CI* | *p* |
| (Intercept) | 17.63 | 4.38 – 30.88 | **0.011** |
| rank(Awareness level) | -0.22 | -0.82 – 0.38 | 0.471 |
| rank(Perception using strategy level) | 0.14 | -0.32 – 0.60 | 0.532 |
| Condition [2D] | 10.26 | -20.52 – 41.03 | 0.504 |
| Condition [Controller] | -4.17 | -24.17 – 15.83 | 0.675 |
| rank(Awareness level) x Condition [2D] | -0.41 | -1.58 – 0.76 | 0.481 |
| rank(Awareness level) x Condition [Controller] | -0.18 | -1.08 – 0.72 | 0.690 |
| rank(Perception using strategy level) x Condition [2D] | 0.50 | -0.12 – 1.13 | 0.111 |
| rank(Perception using strategy level) x Condition [Controller] | 0.75 | 0.09 – 1.41 | **0.028** |
| Observations | 46 | | |
| $R^2$ / $R^2$ adjusted | 0.533 / 0.432 | | |

Results of the linear model on ranks for the mean number of explicit choices in response strategy as a function of perception of using this strategy, awareness of the strategy information, and the VR modality in the first sample.

**Supplementary Table A.3.6:** Choices of response strategy in the second sample

| | rank(N_Explicit_choices) | | |
|---|---|---|---|
| *Predictors* | *Estimates* | *CI* | *p* |
| (Intercept) | 31.46 | 4.59 – 58.34 | **0.022** |
| rank(Awareness level) | -0.15 | -0.84 – 0.54 | 0.658 |
| rank(Perception using strategy level) | 0.38 | -0.04 – 0.79 | 0.073 |
| Condition [2D] | -24.75 | -63.17 – 13.67 | 0.203 |
| Condition [Controller] | 0.88 | -37.83 – 39.59 | 0.964 |
| rank(Awareness level) x Condition [2D] | 0.49 | -0.52 – 1.49 | 0.337 |
| rank(Awareness level) x Condition [Controller] | -0.04 | -1.04 – 0.96 | 0.936 |
| rank(Perception using strategy level) x Condition [2D] | 0.17 | -0.41 – 0.76 | 0.557 |
| rank(Perception using strategy level) x Condition [Controller] | -0.14 | -0.72 – 0.44 | 0.633 |
| Observations | 75 | | |
| $R^2$ / $R^2$ adjusted | 0.207 / 0.111 | | |

Results of the linear model on ranks for the mean number of explicit choices in response strategy as a function of perception of using this strategy, awareness of the strategy information, and the VR modality in the second sample.

**Supplementary Table A.3.7:** Choices of place strategy in the first sample

| | **rank(N_Explicit_choices)** | | |
|---|---|---|---|
| *Predictors* | *Estimates* | *CI* | *p* |
| (Intercept) | 5.73 | -16.20 – 27.65 | 0.600 |
| rank(Awareness level) | 0.04 | -0.50 – 0.58 | 0.889 |
| rank(Perception using strategy level) | 0.65 | 0.12 – 1.18 | **0.018** |
| Condition [2D] | 9.06 | -17.08 – 35.20 | 0.487 |
| Condition [Controller] | -1.75 | -28.35 – 24.85 | 0.895 |
| rank(Awareness level) x Condition [2D] | -0.08 | -0.87 – 0.71 | 0.837 |
| rank(Awareness level) x Condition [Controller] | -0.09 | -0.90 – 0.71 | 0.815 |
| rank(Perception using strategy level) x Condition [2D] | -0.15 | -0.98 – 0.67 | 0.706 |
| rank(Perception using strategy level) x Condition [Controller] | 0.26 | -0.68 – 1.19 | 0.580 |
| Observations | 46 | | |
| $R^2$ / $R^2$ adjusted | 0.337 / 0.194 | | |

Results of the linear model on ranks for the mean number of explicit choices in place strategy as a function of perception of using this strategy, awareness of the strategy information, and the VR modality in the first sample.

**Supplementary Table A.3.8:** Choices of place strategy in the second sample

| | **rank(N_Explicit_choices)** | | |
|---|---|---|---|
| *Predictors* | *Estimates* | *CI* | *p* |
| (Intercept) | 25.71 | 3.74 – 47.67 | **0.022** |
| rank(Awareness level) | 0.13 | -0.30 – 0.56 | 0.546 |
| rank(Perception using strategy level) | 0.10 | -0.44 – 0.64 | 0.716 |
| Condition [2D] | -4.69 | -35.14 – 25.76 | 0.759 |
| Condition [Controller] | -6.54 | -38.12 – 25.04 | 0.680 |
| rank(Awareness level) x Condition [2D] | -0.30 | -0.96 – 0.37 | 0.379 |
| rank(Awareness level) x Condition [Controller] | -0.09 | -0.70 – 0.53 | 0.775 |
| rank(Perception using strategy level) x Condition [2D] | 0.50 | -0.25 – 1.24 | 0.189 |
| rank(Perception using strategy level) x Condition [Controller] | 0.45 | -0.33 – 1.24 | 0.250 |
| Observations | 75 | | |
| $R^2$ / $R^2$ adjusted | 0.160 / 0.058 | | |

Results of the linear model on ranks for the mean number of explicit choices in place strategy as a function of perception of using this strategy, awareness of the strategy information, and the VR modality in the second sample.

### A.4 Eye-tracking measures

#### A.4.1 Main results

The analyses presented in this section correspond to the pilot dataset (the first sample). These results were used to identify candidate effects and guide the design of the preregistered powered analysis for the second sample. While these results are reported here for transparency, they were not used for inferential conclusions and should be interpreted as exploratory.

**Pupil size**

Pupil size was assessed using the mean of the left pupil size in each trial (similar values were also observed for the right pupil data). The pupil size was significantly smaller in the second session (*estimate* = -0.14, $t_{(1180)} = -4.6$, $p = 3.3e{-6}$, Supplementary Fig. A.4.1a). There was no main effect of condition (*estimate* = -0.05, $t_{(29.7)} = -0.3$, $p = 0.77$), and no interaction between session and condition was obtained (*estimate* = -0.06, $t_{(1180)} = -1.4$, $p = 0.14$). Contrast analysis showed that the pupil size decreased from the first session to the second in both VR conditions (*estimate* = 0.14, $t_{(1181)} = 4.7$, $p < 0.0001$) and (*estimate* = 0.2, $t_{(1181)} = 6.8$, $p < 0.0001$, Bonferroni corrected for 2 tests) for the walking and the controller conditions, respectively.

**Blinks**

Blink behavior was assessed as the mean number of blinks normalized by trial duration. A significant main effect of session was found, with more blinks observed in the second session compared to the first (*estimate* = 0.52, $t_{(1499)} = 3.6$, $p = 0.0003$). In addition, a marginal interaction was found between session and condition (*estimate* = -0.37, $t_{(1499)} = -1.8$, $p = 0.07$). Contrast analysis showed a significant increase in the number of blinks from the first session to the second in the walking condition (*estimate* = -0.52, $t_{(1500)} = -3.6$, $p = 0.0003$, Bonferroni corrected for 2 tests, Supplementary Fig. A.4.1b), but not in the controller condition (*estimate* = -0.15, $t_{(1500)} = -1$, $p = 0.03$, Bonferroni corrected for 2 tests).

**Visual attention measured by gaze time**

<u>Gaze time at cue on the reward side:</u> The percentage of time participants gazed at the cue on the reward side showed no significant main effects for condition (*estimate* = -0.32, $t_{(41)} = -0.4$, $p = 0.7$), and (*estimate* = 0.03, $t_{(929)} = 0.05$, $p = 0.96$) and session. In addition, no interaction between condition and session was observed (*estimate* = -0.93, $t_{(925)} = -1.25$, $p = 0.23$, Supplementary Fig. A.4.2).

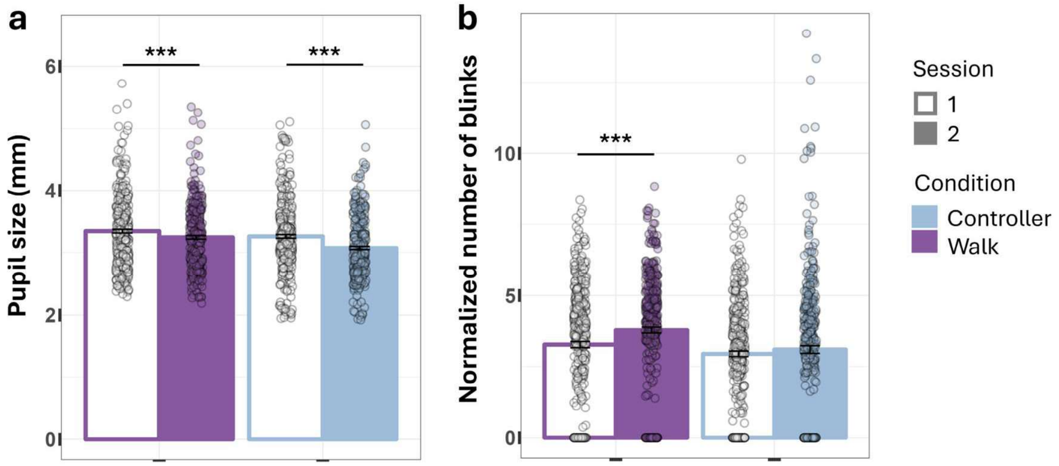


**Supplementary Fig. A.4.1:** Eye-tracking (ET) measures. Bar plots of ET measures according to VR modality of the first sample. The height of each bar presents the mean, with error bars indicating ± 1 SEM. Individual trial-based data points are shown as dots overlaid on the bars. ($p < 0.001$ '***', $p < 0.01$ '**', $p < 0.05$'*').

**Supplementary Table A.4.1:** ET measures summary of results.

| Effect | First sample | Second sample | Replication |
|---|---|---|---|
| Pupil (VR Walk vs Controller main effect) | - | - | Replicated |
| Pupil (Session 1 vs 2 main effect) | + | + | Replicated |
| Pupil (Session x Controller interaction) | - | + | Not replicated |
| Pupil (Session 1 vs 2, VR Walk contrast) | + | + | Replicated |
| Pupil (Session 1 vs 2, Controller contrast) | + | + | Replicated |
| Blinks (VR Walk vs Controller main effect) | - | - | Replicated |
| Blinks (Session 1 vs 2 main effect) | + | - | Not replicated |
| Blinks (Session x Controller interaction) | . | + | Mid replicated |
| Blinks (Session 1 vs 2, VR Walk contrast) | + | - | Not replicated |
| Blinks (Session 1 vs 2, Controller contrast) | - | + | Not replicated |

Summary of the ET measures mixed linear models results for each effect, and sample. (where '+': significant effect; '.' Marginally significant effect; '-': non-significant effect.

Gaze time at surrounding objects on the reward side: The average percentage of time spent gazing at surrounding objects on the reward side was significantly lower by 1.9% in the controller condition relative to the walking condition (*estimate* = -1.9, $t_{(46)} = -2.5$, $p = 0.014$). A significant main effect of session was found (*estimate* = 0.9, $t_{(770)} = 2.1$, $p = 0.03$), but no interaction with condition was observed (*estimate* = -0.56, $t_{(771)} = -0.8$, $p = 0.4$).

Contrast analysis found that the percentage of time the objects in the reward side were gazed at significantly increased from the first session to the second in the walking condition (*estimate* = -0.9, $t_{(770)}$ = -2.1, $p$ = 0.03, Bonferroni corrected for 2 tests, Supplementary Fig. A.4.2). In addition, in both sessions, the percentage of time the objects on the reward side were gazed at, was significantly higher in the walking condition compared to the controller (*estimate* = 1.9, $t_{(43.8)}$ = 2.5, $p$ = 0.014) for the first session, and (*estimate* = 2.5, $t_{(44.1)}$ = 3.3, $p$ = 0.002, Bonferroni corrected for 2 tests) for the second session).

Gaze time at all surrounding objects: Participants gazed at all the surrounding objects (objects $O_1 - O_5$ in Fig. 4) for 8.3% of the trial duration on average. The percentage of time all the objects were gazed at was significantly lower in the controller condition compared to walking (*estimate* = -3.3, $t_{(39)}$ = -3.1, $p$ = 0.003). However, there was no difference between the two sessions as a main effect of session (*estimate* = -0.06, $t_{(1058)}$ = -0.1, $p$ = 0.91), or as an interaction between session and condition (*estimate* = -0.01, $t_{(1059)}$ = -0.02, $p$ = 0.98).

Contrast analysis showed that the condition effect was present in both sessions (*estimate* = 3.3, $t_{(38.1)}$ = 3.1, $p$ = 0.003) for the first session, and (*estimate* = 3.3, $t_{(40)}$ = 3.1, $p$ = 0.004, Bonferroni corrected for 2 tests) for the second session.

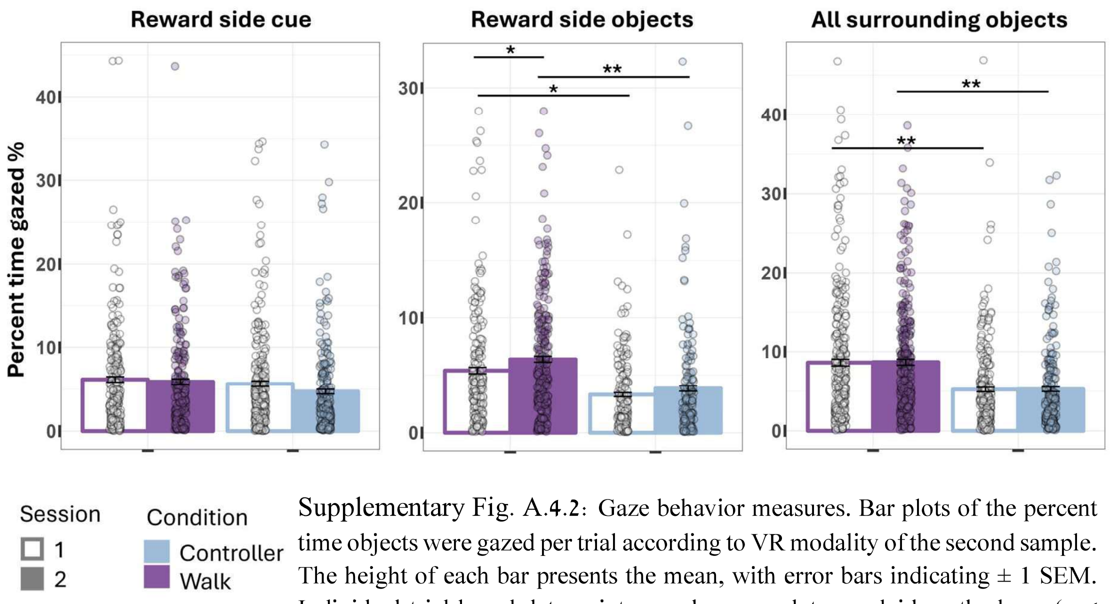


Supplementary Fig. A.4.2: Gaze behavior measures. Bar plots of the percent time objects were gazed per trial according to VR modality of the second sample. The height of each bar presents the mean, with error bars indicating ± 1 SEM. Individual trial-based data points are shown as dots overlaid on the bars. (p < 0.001 '***', p < 0.01 '**', p < 0.05'*').

**Supplementary Table A.4.2:** Visual attention summary of results.

| **Effect** | **First sample** | **Second sample** | **Replication** |
|---|---|---|---|
| Cue on the reward side (VR Walk vs Controller main effect) | - | - | Replicated |
| Cue on the reward side (Session 1 vs 2 main effect) | - | - | Replicated |
| Cue on the reward side (Session x Controller interaction) | - | + | Not replicated |
| Cue on the reward side (Session 1 vs 2, Controller contrast) | - | + | Not replicated |
| Cue on the reward side (Session 2 VR Walk vs Controller contrast) | - | + | Not replicated |
| Objects on the reward side (VR Walk vs Controller main effect) | + | + | Replicated |
| Objects on the reward side (Session 1 vs 2 main effect) | + | - | Not replicated |
| Objects on the reward side (Session x Controller interaction) | - | + | Not replicated |
| Objects on the reward side (Session 1 vs 2, VR Walk contrast) | + | - | Not replicated |
| Objects on the reward side (Session 1 vs 2, Controller contrast) | - | + | Not replicated |
| Objects on the reward side (Session 1 VR Walk vs Controller contrast) | + | + | Replicated |
| Objects on the reward side (Session 2 VR Walk vs Controller contrast) | + | - | Not replicated |
| All objects (VR Walk vs Controller main effect) | + | + | Replicated |
| All objects (Session 1 vs 2 main effect) | - | - | Replicated |
| All objects (Session x Controller interaction) | - | - | Replicated |
| All objects (Session 1 VR Walk vs Controller contrast) | + | + | Replicated |
| All objects (Session 2 VR Walk vs Controller contrast) | + | - | Not replicated |

Summary of the visual attention measured by gaze time mixed linear models results for each effect, and sample. (where '+': significant effect; '.' Marginally significant effect; '-': non-significant effect.

#### A.4.2 Visual attention relation to performance

**Supplementary Table A.4.3:** Correlation between learning pace and gaze at the cue on the reward side in the first sample.

| | rank(n_train) | | |
|---|---|---|---|
| *Predictors* | *Estimates* | *CI* | *p* |
| (Intercept) | 38.12 | 16.97 – 59.27 | **0.001** |
| rank(Maze Reward cue percent time gazed) | -0.12 | -0.67 – 0.43 | 0.666 |
| Condition [Controller] | -1.56 | -31.57 – 28.45 | 0.917 |
| Sess [2] | -15.35 | -42.38 – 11.69 | 0.260 |
| rank(Maze Reward cue percent time gazed) x Condition [Controller] | 0.12 | -0.68 – 0.93 | 0.759 |
| rank(Maze Reward cue percent time gazed) x Sess [2] | 0.19 | -0.55 – 0.92 | 0.614 |
| Condition [Controller] x Sess [2] | 9.73 | -28.78 – 48.23 | 0.614 |
| (rank(Maze Reward cue percent time gazed) x Condition [Controller]) x Sess [2] | -0.19 | -1.25 – 0.87 | 0.716 |
| **Random Effects** | | | |
| $\sigma^2$ | 331.73 | | |
| $\tau_{00\ SubID}$ | 0.00 | | |
| $N_{SubID}$ | 31 | | |
| Observations | 62 | | |
| Marginal $R^2$ / Conditional $R^2$ | 0.062 / NA | | |

**Supplementary Table A.4.4:** Correlation between learning pace and gaze at the cue on the reward side in the second sample.

| | rank(n_train) | | |
|---|---|---|---|
| *Predictors* | *Estimates* | *CI* | *p* |
| (Intercept) | 65.37 | 30.60 – 100.13 | **<0.001** |
| rank(Maze Reward cue percent time gazed) | -0.29 | -0.68 – 0.11 | 0.152 |
| Condition [Controller] | -15.28 | -48.04 – 17.48 | 0.357 |
| Sess [2] | -27.13 | -57.85 – 3.60 | 0.083 |
| rank(SOT) | 0.09 | -0.18 – 0.35 | 0.517 |
| rank(MRT) | -0.01 | -0.27 – 0.25 | 0.911 |
| rank(Maze Reward cue percent time gazed) x Condition [Controller] | 0.31 | -0.25 – 0.88 | 0.275 |
| rank(Maze Reward cue percent time gazed) x Sess [2] | 0.39 | -0.16 – 0.93 | 0.160 |
| Condition [Controller] x Sess [2] | 1.79 | -43.40 – 46.98 | 0.937 |
| (rank(Maze Reward cue percent time gazed) x Condition [Controller]) x Sess [2] | -0.10 | -0.88 – 0.69 | 0.807 |
| **Random Effects** | | | |
| $\sigma^2$ | 757.97 | | |
| $\tau_{00}$ SubID | 61.67 | | |
| ICC | 0.08 | | |
| N SubID | 49 | | |
| Observations | 100 | | |
| Marginal $R^2$ / Conditional $R^2$ | 0.078 / 0.148 | | |

**Supplementary Table A.4.5:** Correlation between learning pace and gaze at objects on the reward side in the first sample.

| | rank(n_train) | | |
|---|---|---|---|
| *Predictors* | *Estimates* | *CI* | *p* |
| (Intercept) | 28.96 | 11.69 – 46.24 | **0.001** |
| rank(Objects Reward percent time gazed) | 0.17 | -0.32 – 0.65 | 0.497 |
| Condition [Controller] | 18.89 | -6.78 – 44.55 | 0.146 |
| Sess [2] | 1.87 | -30.24 – 33.99 | 0.907 |
| rank(Objects Reward percent time gazed) x Condition [Controller] | -0.62 | -1.46 – 0.21 | 0.141 |
| rank(Objects Reward percent time gazed) x Sess [2] | -0.32 | -1.11 – 0.48 | 0.428 |
| Condition [Controller] x Sess [2] | -19.10 | -59.89 – 21.68 | 0.352 |
| (rank(Objects Reward percent time gazed) x Condition [Controller]) x Sess [2] | 0.78 | -0.36 – 1.92 | 0.175 |
| **Random Effects** | | | |
| $\sigma^2$ | 318.48 | | |
| $\tau_{00\ SubID}$ | 0.00 | | |
| $N_{SubID}$ | 31 | | |
| Observations | 62 | | |
| Marginal $R^2$ / Conditional $R^2$ | 0.095 / NA | | |

**Supplementary Table A.4.6:** Correlation between learning pace and gaze at objects on the reward side in the second sample.

| | rank(n_train) | | |
|---|---|---|---|
| *Predictors* | *Estimates* | *CI* | *p* |
| (Intercept) | 43.07 | 4.76 – 81.38 | **0.028** |
| rank(Objects Reward percent time gazed) | 0.12 | -0.32 – 0.55 | 0.596 |
| Condition [Controller] | 8.28 | -27.43 – 43.99 | 0.646 |
| Sess [2] | -5.03 | -42.10 – 32.03 | 0.788 |
| rank(SOT) | 0.11 | -0.16 – 0.38 | 0.424 |
| rank(MRT) | -0.01 | -0.28 – 0.26 | 0.939 |
| rank(Objects Reward percent time gazed) x Condition [Controller] | -0.14 | -0.76 – 0.47 | 0.644 |
| rank(Objects Reward percent time gazed) x Sess [2] | -0.05 | -0.66 – 0.56 | 0.867 |
| Condition [Controller] x Sess [2] | -3.27 | -51.29 – 44.75 | 0.893 |
| (rank(Objects Reward percent time gazed) x Condition [Controller]) x Sess [2] | 0.03 | -0.80 – 0.86 | 0.942 |
| **Random Effects** | | | |
| $\sigma^2$ | 812.75 | | |
| $\tau_{00\ SubID}$ | 44.96 | | |
| ICC | 0.05 | | |
| $N_{SubID}$ | 49 | | |
| Observations | 100 | | |
| Marginal $R^2$ / Conditional $R^2$ | 0.038 / 0.088 | | |

**Supplementary Table A.4.7:** Correlation between learning pace and gaze at all objects in the first sample.

| | rank(n_train) | | |
|---|---|---|---|
| *Predictors* | *Estimates* | *CI* | *p* |
| (Intercept) | 30.96 | 6.19 – 55.72 | **0.015** |
| rank(Objects percent time gazed) | 0.08 | -0.51 – 0.66 | 0.793 |
| Condition [Controller] | 20.67 | -9.82 – 51.16 | 0.180 |
| Sess [2] | -6.60 | -39.00 – 25.81 | 0.685 |
| rank(Objects percent time gazed) x Condition [Controller] | -0.66 | -1.49 – 0.17 | 0.119 |
| rank(Objects percent time gazed) x Sess [2] | -0.07 | -0.87 – 0.73 | 0.869 |
| Condition [Controller] x Sess [2] | -9.71 | -49.40 – 29.98 | 0.625 |
| (rank(Objects percent time gazed) x Condition [Controller]) x Sess [2] | 0.47 | -0.62 – 1.57 | 0.388 |
| **Random Effects** | | | |
| $\sigma^2$ | 307.50 | | |
| $\tau_{00\ SubID}$ | 0.00 | | |
| $N_{SubID}$ | 31 | | |
| Observations | 62 | | |
| Marginal $R^2$ / Conditional $R^2$ | 0.124 / NA | | |

**Supplementary Table A.4.8:** Correlation between learning pace and gaze at all objects in the second sample.

| | rank(n_train) | | |
|---|---|---|---|
| *Predictors* | *Estimates* | *CI* | *p* |
| (Intercept) | 24.30 | -15.02 – 63.62 | 0.223 |
| rank(Objects percent time gazed) | 0.36 | -0.05 – 0.78 | 0.086 |
| Condition [Controller] | 23.79 | -12.73 – 60.31 | 0.199 |
| Sess [2] | 2.47 | -35.22 – 40.16 | 0.897 |
| rank(SOT) | 0.15 | -0.12 – 0.43 | 0.270 |
| rank(MRT) | 0.00 | -0.26 – 0.27 | 0.994 |
| rank(Objects percent time gazed) x Condition [Controller] | -0.37 | -0.98 – 0.24 | 0.230 |
| rank(Objects percent time gazed) x Sess [2] | -0.16 | -0.74 – 0.42 | 0.587 |
| Condition [Controller] x Sess [2] | -16.44 | -64.74 – 31.86 | 0.500 |
| (rank(Objects percent time gazed) x Condition [Controller]) x Sess [2] | 0.27 | -0.57 – 1.12 | 0.521 |
| **Random Effects** | | | |
| $\sigma^2$ | 757.61 | | |
| $\tau_{00\ SubID}$ | 70.52 | | |
| ICC | 0.09 | | |
| $N_{SubID}$ | 50 | | |
| Observations | 100 | | |
| Marginal $R^2$ / Conditional $R^2$ | 0.072 / 0.151 | | |

**Supplementary Table A.4.9:** Correlation between learning effectiveness and gaze at the cue on the reward side in the first sample.

| | rank(Probe_Succ) | | |
|---|---|---|---|
| *Predictors* | *Estimates* | *CI* | *p* |
| (Intercept) | 42.31 | 24.94 – 59.68 | **<0.001** |
| rank(Maze Reward cue percent time gazed) | -0.29 | -0.74 – 0.17 | 0.211 |
| Condition [Controller] | -19.39 | -44.03 – 5.26 | 0.121 |
| Sess [2] | -7.63 | -29.62 – 14.37 | 0.490 |
| rank(Maze Reward cue percent time gazed) x Condition [Controller] | 0.31 | -0.35 – 0.97 | 0.347 |
| rank(Maze Reward cue percent time gazed) x Sess [2] | 0.33 | -0.29 – 0.94 | 0.291 |
| Condition [Controller] x Sess [2] | 18.26 | -12.76 – 49.27 | 0.243 |
| (rank(Maze Reward cue percent time gazed) x Condition [Controller]) x Sess [2] | -0.35 | -1.22 – 0.52 | 0.420 |
| **Random Effects** | | | |
| $\sigma^2$ | 186.05 | | |
| $\tau_{00\ SubID}$ | 44.51 | | |
| ICC | 0.19 | | |
| $N_{SubID}$ | 31 | | |
| Observations | 62 | | |
| Marginal $R^2$ / Conditional $R^2$ | 0.105 / 0.278 | | |

**Supplementary Table A.4.10:** Correlation between learning effectiveness and gaze at the cue on the reward side in the second sample.

| | rank(Probe_Succ) | | |
|---|---|---|---|
| *Predictors* | *Estimates* | *CI* | *p* |
| (Intercept) | 73.95 | 40.34 – 107.57 | **<0.001** |
| rank(Maze Reward cue percent time gazed) | -0.26 | -0.65 – 0.12 | 0.174 |
| Condition [Controller] | -17.99 | -49.75 – 13.78 | 0.264 |
| Sess [2] | -19.35 | -49.25 – 10.55 | 0.202 |
| rank(SOT) | -0.23 | -0.49 – 0.02 | 0.073 |
| rank(MRT) | 0.07 | -0.19 – 0.32 | 0.608 |
| rank(Maze Reward cue percent time gazed) x Condition [Controller] | 0.23 | -0.32 – 0.78 | 0.411 |
| rank(Maze Reward cue percent time gazed) x Sess [2] | 0.38 | -0.14 – 0.91 | 0.153 |
| Condition [Controller] x Sess [2] | 19.98 | -23.97 – 63.92 | 0.369 |
| (rank(Maze Reward cue percent time gazed) x Condition [Controller]) x Sess [2] | -0.26 | -1.02 – 0.50 | 0.493 |
| **Random Effects** | | | |
| $\sigma^2$ | 720.16 | | |
| $\tau_{00\ SubID}$ | 49.80 | | |
| ICC | 0.06 | | |
| $N_{SubID}$ | 49 | | |
| Observations | 100 | | |
| Marginal $R^2$ / Conditional $R^2$ | 0.101 / 0.159 | | |

**Supplementary Table A.4.11:** Correlation between learning effectiveness and gaze at the objects on the reward side in the first sample.

| | **rank(Probe_Succ)** | | |
|---|---|---|---|
| *Predictors* | *Estimates* | *CI* | *p* |
| (Intercept) | 28.75 | 14.41 – 43.09 | **<0.001** |
| rank(Objects Reward percent time gazed) | 0.12 | -0.28 – 0.52 | 0.543 |
| Condition [Controller] | 0.83 | -20.46 – 22.12 | 0.938 |
| Sess [2] | 7.95 | -17.76 – 33.66 | 0.538 |
| rank(Objects Reward percent time gazed) x Condition [Controller] | -0.36 | -1.05 – 0.33 | 0.298 |
| rank(Objects Reward percent time gazed) x Sess [2] | -0.14 | -0.78 – 0.50 | 0.656 |
| Condition [Controller] x Sess [2] | 3.40 | -29.36 – 36.16 | 0.836 |
| (rank(Objects Reward percent time gazed) x Condition [Controller]) x Sess [2] | 0.13 | -0.80 – 1.07 | 0.775 |
| **Random Effects** | | | |
| $\sigma^2$ | 179.42 | | |
| $\tau_{00\ SubID}$ | 46.86 | | |
| ICC | 0.21 | | |
| $N_{SubID}$ | 31 | | |
| Observations | 62 | | |
| Marginal $R^2$ / Conditional $R^2$ | 0.121 / 0.303 | | |

**Supplementary Table A.4.12:** Correlation between learning effectiveness and gaze at the objects on the reward side in the second sample.

| | rank(Probe_Succ) | | |
|---|---|---|---|
| *Predictors* | *Estimates* | *CI* | *p* |
| (Intercept) | 47.97 | 11.14 – 84.80 | **0.011** |
| rank(Objects Reward percent time gazed) | 0.20 | -0.21 – 0.61 | 0.334 |
| Condition [Controller] | 16.95 | -16.78 – 50.68 | 0.321 |
| Sess [2] | 12.34 | -22.19 – 46.87 | 0.479 |
| rank(SOT) | -0.20 | -0.47 – 0.06 | 0.130 |
| rank(MRT) | 0.06 | -0.20 – 0.33 | 0.640 |
| rank(Objects Reward percent time gazed) x Condition [Controller] | -0.48 | -1.06 – 0.11 | 0.109 |
| rank(Objects Reward percent time gazed) x Sess [2] | -0.23 | -0.80 – 0.34 | 0.422 |
| Condition [Controller] x Sess [2] | -18.42 | -63.22 – 26.38 | 0.416 |
| (rank(Objects Reward percent time gazed) x Condition [Controller]) x Sess [2] | 0.53 | -0.26 – 1.31 | 0.185 |
| **Random Effects** | | | |
| $\sigma^2$ | 677.78 | | |
| $\tau_{00\ SubID}$ | 95.77 | | |
| ICC | 0.12 | | |
| N $_{SubID}$ | 49 | | |
| Observations | 100 | | |
| Marginal $R^2$ / Conditional $R^2$ | 0.102 / 0.213 | | |

**Supplementary Table A.4.13:** Correlation between learning effectiveness and gaze at all objects in the first sample.

| | rank(Probe_Succ) | | |
|---|---|---|---|
| *Predictors* | *Estimates* | *CI* | *p* |
| (Intercept) | 21.18 | 0.48 – 41.89 | **0.045** |
| rank(Objects percent time gazed) | 0.28 | -0.20 – 0.77 | 0.247 |
| Condition [Controller] | 9.41 | -16.08 – 34.91 | 0.462 |
| Sess [2] | 17.09 | -9.31 – 43.48 | 0.200 |
| rank(Objects percent time gazed) x Condition [Controller] | -0.55 | -1.25 – 0.15 | 0.119 |
| rank(Objects percent time gazed) x Sess [2] | -0.35 | -1.01 – 0.30 | 0.285 |
| Condition [Controller] x Sess [2] | -8.31 | -40.70 – 24.07 | 0.609 |
| (rank(Objects percent time gazed) x Condition [Controller]) x Sess [2] | 0.39 | -0.51 – 1.29 | 0.391 |
| **Random Effects** | | | |
| $\sigma^2$ | 187.17 | | |
| $\tau_{00}$ SubID | 31.50 | | |
| ICC | 0.14 | | |
| N SubID | 31 | | |
| Observations | 62 | | |
| Marginal $R^2$ / Conditional $R^2$ | 0.140 / 0.264 | | |

**Supplementary Table A.4.14:** Correlation between learning effectiveness and gaze at all objects in the second sample.

| | rank(Probe_Succ) | | |
|---|---|---|---|
| *Predictors* | *Estimates* | *CI* | *p* |
| (Intercept) | 72.06 | 33.61 – 110.50 | **<0.001** |
| rank(Objects percent time gazed) | -0.17 | -0.57 – 0.24 | 0.416 |
| Condition [Controller] | -10.79 | -46.30 – 24.73 | 0.548 |
| Sess [2] | -8.05 | -44.49 – 28.39 | 0.662 |
| rank(SOT) | -0.25 | -0.52 – 0.02 | 0.069 |
| rank(MRT) | 0.06 | -0.20 – 0.32 | 0.634 |
| rank(Objects percent time gazed) x Condition [Controller] | 0.03 | -0.57 – 0.62 | 0.932 |
| rank(Objects percent time gazed) x Sess [2] | 0.12 | -0.44 – 0.68 | 0.677 |
| Condition [Controller] x Sess [2] | 9.23 | -37.50 – 55.97 | 0.696 |
| (rank(Objects percent time gazed) x Condition [Controller]) x Sess [2] | 0.01 | -0.81 – 0.82 | 0.989 |
| **Random Effects** | | | |
| $\sigma^2$ | 700.81 | | |
| $\tau_{00\ SubID}$ | 84.61 | | |
| ICC | 0.11 | | |
| $N_{SubID}$ | 50 | | |
| Observations | 100 | | |
| Marginal $R^2$ / Conditional $R^2$ | 0.088 / 0.186 | | |

#### *A.5 Identification of individual strategy usage based on eye tracking and path features*

**Table A.5.1:** Classification of the choice in strategy per trial in sample 1.

| Strategy | Condition | Session | Accuracy | P-value | Positive class proportion |
|---|---|---|---|---|---|
| cue | 2D | 1 | 0.4 | 1 | 0.44 |
| cue | 2D | 2 | 0.51 | 0.218 | 0.49 |
| cue | Walk | 1 | 0.54 | 0.396 | 0.58 |
| cue | Walk | 2 | 0.37 | 1 | 0.58 |
| cue | Controller | 1 | 0.4 | 1 | 0.49 |
| cue | Controller | 2 | 0.47 | 1 | 0.58 |
| place | 2D | 1 | 0.62 | 0.703 | 0.33 |
| place | 2D | 2 | 0.42 | 1 | 0.42 |
| place | Walk | 1 | 0.62 | 0.96 | 0.29 |
| place | Walk | 2 | 0.6 | 0.069 | 0.52 |
| place | Controller | 1 | 0.67 | 0.861 | 0.29 |
| place | Controller | 2 | 0.78 | 0.99 | 0.18 |
| response | 2D | 1 | 0.27 | 1 | 0.56 |
| response | 2D | 2 | 0.31 | 1 | 0.44 |
| response | Walk | 1 | 0.75 | 1 | 0.17 |
| response | Walk | 2 | 0.79 | 0.861 | 0.21 |
| response | Controller | 1 | 0.42 | 1 | 0.42 |
| response | Controller | 2 | 0.42 | 1 | 0.38 |

Results of the models of classification of the choice in strategy per trial. For each strategy, VR modality, and session number, the model's accuracy, P-value, and positive class proportion are presented (a value close to 0.5 describes balanced data). Models with significant results greater than chance are marked in bold.

**Table A.5.2:** Classification of whether a strategy was preferred based on training trials only in sample 1.

| Strategy | Condition | Session | Accuracy | P-value | Positive class proportion |
|---|---|---|---|---|---|
| cue | 2D | 1 | 0.4 | 1 | 0.4 |
| cue | 2D | 2 | 0.53 | 0.861 | 0.33 |
| **cue** | **Walk** | **1** | **0.75** | **0.04** | **0.69** |
| cue | Walk | 2 | 0.25 | 1 | 0.5 |
| cue | Controller | 1 | 0.47 | 1 | 0.4 |
| cue | Controller | 2 | 0.13 | 1 | 0.53 |
| place | 2D | 1 | 0.8 | 0.96 | 0.2 |
| place | 2D | 2 | 0.33 | 1 | 0.4 |
| place | Walk | 1 | 0.62 | 0.624 | 0.31 |
| place | Walk | 2 | 0.44 | 1 | 0.44 |
| place | Controller | 1 | 0.8 | 0.911 | 0.2 |
| place | Controller | 2 | 0.87 | 1 | 0.13 |
| response | 2D | 1 | 0.2 | 1 | 0.53 |
| response | 2D | 2 | 0.53 | 0.871 | 0.33 |
| response | Walk | 1 | 0.88 | 1 | 0.12 |
| response | Walk | 2 | 0.88 | 1 | 0.06 |
| response | Controller | 1 | 0.4 | 1 | 0.4 |
| response | Controller | 2 | 0.33 | 1 | 0.4 |

Results of the models of classification of a preferred strategy based on data from training trials only. For each strategy, VR modality, and session number, the model's accuracy, P-value, and positive class proportion are presented (a value close to 0.5 describes balanced data). Models with significant results greater than chance are marked in bold.

**Table A.5.3:** Classification of whether a strategy was preferred based on all trials in sample 1.

| Strategy | Condition | Session | Accuracy | P-value | Positive class proportion |
|---|---|---|---|---|---|
| cue | 2D | 1 | 0.47 | 1 | 0.4 |
| cue | 2D | 2 | 0.47 | 1 | 0.33 |
| **cue** | **Walk** | **1** | **0.81** | **0.03** | **0.69** |
| cue | Walk | 2 | 0.56 | 0.198 | 0.5 |
| cue | Controller | 1 | 0.6 | 0.277 | 0.4 |
| cue | Controller | 2 | 0.2 | 1 | 0.53 |
| place | 2D | 1 | 0.8 | 0.921 | 0.2 |
| place | 2D | 2 | 0.4 | 1 | 0.4 |
| place | Walk | 1 | 0.56 | 0.832 | 0.31 |
| place | Walk | 2 | 0.56 | 0.307 | 0.44 |
| place | Controller | 1 | 0.8 | 0.95 | 0.2 |
| place | Controller | 2 | 0.87 | 0.98 | 0.13 |
| response | 2D | 1 | 0.27 | 1 | 0.53 |
| response | 2D | 2 | 0.6 | 0.752 | 0.33 |
| response | Walk | 1 | 0.88 | 1 | 0.12 |
| response | Walk | 2 | 0.88 | 1 | 0.06 |
| response | Controller | 1 | 0.33 | 1 | 0.4 |
| response | Controller | 2 | 0.47 | 1 | 0.4 |

Results of the models of classification of a preferred strategy based on data from all trials. For each strategy, VR modality, and session number, the model's accuracy, P-value, and positive class proportion are presented (a value close to 0.5 describes balanced data). Models with significant results greater than chance are marked in bold.

**Table A.5.4:** Regression of the number of explicit choices in strategy based on training trials only in sample 1.

| Strategy | Condition | Session | $R^2$ | RMSE | P-value $R^2$ | P-value RMSE | Mean #choices |
|---|---|---|---|---|---|---|---|
| cue | 2D | 1 | -0.89 | 1.79 | 1 | 1 | 1.33 |
| cue | 2D | 2 | -0.99 | 1.53 | 1 | 1 | 1.47 |
| cue | Walk | 1 | -0.07 | 1.07 | 1 | 1 | 1.75 |
| **cue** | **Walk** | **2** | **0.2** | **1.07** | **0.04** | **0.04** | **1.75** |
| cue | Controller | 1 | -0.53 | 1.49 | 1 | 1 | 1.47 |
| cue | Controller | 2 | -1.46 | 2.18 | 1 | 1 | 1.73 |
| place | 2D | 1 | -0.66 | 1.33 | 1 | 1 | 1 |
| place | 2D | 2 | -0.86 | 1.53 | 1 | 1 | 1.27 |
| place | Walk | 1 | -0.74 | 1.54 | 1 | 1 | 0.88 |
| place | Walk | 2 | 0.09 | 1.26 | 0.089 | 0.089 | 1.56 |
| place | Controller | 1 | -0.63 | 1.03 | 1 | 1 | 0.87 |
| place | Controller | 2 | -0.45 | 1.31 | 1 | 1 | 0.53 |
| response | 2D | 1 | -1.27 | 1.71 | 1 | 1 | 1.67 |
| response | 2D | 2 | -0.43 | 1.36 | 1 | 1 | 1.33 |
| response | Walk | 1 | 0.11 | 0.82 | 0.079 | 0.079 | 0.5 |
| response | Walk | 2 | -0.15 | 1 | 1 | 1 | 0.62 |
| response | Controller | 1 | -0.81 | 1.51 | 1 | 1 | 1.27 |
| response | Controller | 2 | -0.99 | 1.85 | 1 | 1 | 1.13 |

Results of the models of regression of the number of explicit choices in strategy based on data from training trials only. For each strategy, VR modality, and session number, the model's R2, RMSE, R2 P-value, RMSE P-value, and mean number of choices in strategy are presented. Models with significant results greater than chance are marked in bold.

**Table A.5.5:** Regression of the number of explicit choices in strategy based on all trials in sample 1.

| Strategy | Condition | Session | R2 | RMSE | P-value R2 | P-value RMSE | Mean #choices |
|---|---|---|---|---|---|---|---|
| cue | 2D | 1 | -0.87 | 1.78 | 1 | 1 | 1.33 |
| cue | 2D | 2 | -0.92 | 1.51 | 1 | 1 | 1.47 |
| cue | Walk | 1 | -0.28 | 1.16 | 1 | 1 | 1.75 |
| cue | Walk | 2 | 0.04 | 1.17 | 0.099 | 0.099 | 1.75 |
| cue | Controller | 1 | -0.37 | 1.41 | 1 | 1 | 1.47 |
| cue | Controller | 2 | -1.14 | 2.03 | 1 | 1 | 1.73 |
| place | 2D | 1 | -0.41 | 1.23 | 1 | 1 | 1 |
| place | 2D | 2 | -0.41 | 1.33 | 1 | 1 | 1.27 |
| place | Walk | 1 | -0.57 | 1.46 | 1 | 1 | 0.88 |
| place | Walk | 2 | -0.32 | 1.52 | 1 | 1 | 1.56 |
| place | Controller | 1 | -0.05 | 0.82 | 1 | 1 | 0.87 |
| place | Controller | 2 | -0.2 | 1.19 | 1 | 1 | 0.53 |
| response | 2D | 1 | -0.31 | 1.3 | 1 | 1 | 1.67 |
| response | 2D | 2 | -0.48 | 1.38 | 1 | 1 | 1.33 |
| **response** | **Walk** | **1** | **0.4** | **0.67** | **0.01** | **0.01** | **0.5** |
| **response** | **Walk** | **2** | **0.43** | **0.7** | **0.04** | **0.04** | **0.62** |
| response | Controller | 1 | -1.28 | 1.7 | 1 | 1 | 1.27 |
| response | Controller | 2 | -0.51 | 1.61 | 1 | 1 | 1.13 |

Results of the models of regression of the number of explicit choices in strategy based on all trials' data. For each strategy, VR modality, and session number, the model's R2, RMSE, R2 P-value, RMSE P-value, and mean number of choices in strategy are presented. Models with significant results greater than chance are marked in bold.

**Table A.5.6:** Classification of the choice in strategy per trial in sample 2.

| Strategy | Condition | Session | Accuracy | P-value | Positive class proportion |
|---|---|---|---|---|---|
| cue | 2D | 1 | 0.59 | 0.079 | 0.55 |
| cue | 2D | 2 | 0.59 | 0.465 | 0.61 |
| cue | Walk | 1 | 0.61 | 0.723 | 0.65 |
| cue | Walk | 2 | 0.33 | 1 | 0.48 |
| cue | Controller | 1 | 0.52 | 0.921 | 0.61 |
| cue | Controller | 2 | 0.6 | 0.842 | 0.65 |
| place | 2D | 1 | 0.63 | 0.871 | 0.33 |
| place | 2D | 2 | 0.73 | 0.416 | 0.27 |
| place | Walk | 1 | 0.79 | 0.782 | 0.21 |
| **place** | **Walk** | **2** | **0.76** | **0.04** | **0.27** |
| place | Controller | 1 | 0.63 | 0.297 | 0.36 |
| place | Controller | 2 | 0.68 | 0.713 | 0.29 |
| response | 2D | 1 | 0.49 | 1 | 0.43 |
| response | 2D | 2 | 0.56 | 0.931 | 0.37 |
| response | Walk | 1 | 0.59 | 0.416 | 0.39 |
| **response** | **Walk** | **2** | **0.64** | **0.05** | **0.47** |
| **response** | **Controller** | **1** | **0.69** | **0.02** | **0.35** |
| response | Controller | 2 | 0.64 | 0.634 | 0.32 |

Results of the models of classification of the choice in strategy per trial. For each strategy, VR modality, and session number, the model's accuracy, P-value, and positive class proportion are presented (a value close to 0.5 describes balanced data). Models with significant results greater than chance are marked in bold.

**Table A.5.7:** Classification of whether a strategy was preferred based on training trials only in sample 2.

| Strategy | Condition | Session | Accuracy | P-value | Positive class proportion |
|---|---|---|---|---|---|
| cue | 2D | 1 | 0.56 | 0.455 | 0.6 |
| cue | 2D | 2 | 0.52 | 0.515 | 0.6 |
| cue | Walk | 1 | 0.56 | 0.683 | 0.64 |
| cue | Walk | 2 | 0.44 | 1 | 0.44 |
| cue | Controller | 1 | 0.48 | 1 | 0.64 |
| cue | Controller | 2 | 0.52 | 0.782 | 0.64 |
| place | 2D | 1 | 0.76 | 0.851 | 0.24 |
| place | 2D | 2 | 0.8 | 0.911 | 0.2 |
| place | Walk | 1 | 0.8 | 0.851 | 0.2 |
| place | Walk | 2 | 0.8 | 0.832 | 0.2 |
| place | Controller | 1 | 0.8 | 0.921 | 0.2 |
| place | Controller | 2 | 0.76 | 0.713 | 0.24 |
| response | 2D | 1 | 0.68 | 0.564 | 0.32 |
| response | 2D | 2 | 0.64 | 0.713 | 0.32 |
| response | Walk | 1 | 0.56 | 0.644 | 0.36 |
| **response** | **Walk** | **2** | **0.72** | **0.05** | **0.48** |
| response | Controller | 1 | 0.72 | 0.584 | 0.28 |
| response | Controller | 2 | 0.72 | 0.584 | 0.28 |

Results of the models of classification of a preferred strategy based on data from training trials only. For each strategy, VR modality, and session number, the model's accuracy, P-value, and positive class proportion are presented (a value close to 0.5 describes balanced data). Models with significant results greater than chance are marked in bold.

**Table A.5.8:** Classification of whether a strategy was preferred based on all trials in sample 2.

| Strategy | Condition | Session | Accuracy | P-value | Positive class proportion |
|---|---|---|---|---|---|
| cue | 2D | 1 | 0.72 | 0.079 | 0.6 |
| cue | 2D | 2 | 0.52 | 0.584 | 0.6 |
| cue | Walk | 1 | 0.56 | 0.545 | 0.64 |
| **cue** | **Walk** | **2** | **0.68** | **0.05** | **0.44** |
| cue | Controller | 1 | 0.6 | 0.426 | 0.64 |
| cue | Controller | 2 | 0.56 | 0.624 | 0.64 |
| place | 2D | 1 | 0.76 | 0.861 | 0.24 |
| place | 2D | 2 | 0.8 | 0.901 | 0.2 |
| place | Walk | 1 | 0.8 | 0.842 | 0.2 |
| place | Walk | 2 | 0.8 | 0.802 | 0.2 |
| place | Controller | 1 | 0.8 | 0.861 | 0.2 |
| place | Controller | 2 | 0.76 | 0.634 | 0.24 |
| response | 2D | 1 | 0.68 | 0.554 | 0.32 |
| response | 2D | 2 | 0.68 | 0.455 | 0.32 |
| response | Walk | 1 | 0.68 | 0.158 | 0.36 |
| response | Walk | 2 | 0.44 | 1 | 0.48 |
| response | Controller | 1 | 0.72 | 0.594 | 0.28 |
| response | Controller | 2 | 0.72 | 0.485 | 0.28 |

Results of the models of classification of a preferred strategy based on data from all trials. For each strategy, VR modality, and session number, the model's accuracy, P-value, and positive class proportion are presented (a value close to 0.5 describes balanced data). Models with significant results greater than chance are marked in bold.

**Table A.5.9:** Regression of the number of explicit choices in strategy based on training trials only in sample 2.

| Strategy | Condition | Session | $R^2$ | RMSE | P-value $R^2$ | P-value RMSE | Mean #choices |
|---|---|---|---|---|---|---|---|
| cue | 2D | 1 | -0.48 | 1.41 | 1 | 1 | 1.64 |
| cue | 2D | 2 | -0.45 | 1.51 | 1 | 1 | 1.84 |
| cue | Walk | 1 | -0.31 | 1.14 | 1 | 1 | 1.96 |
| cue | Walk | 2 | -0.94 | 1.76 | 1 | 1 | 1.44 |
| **cue** | **Controller** | **1** | **0.17** | **1.02** | **0.04** | **0.04** | **1.84** |
| cue | Controller | 2 | -0.61 | 1.41 | 1 | 1 | 1.96 |
| place | 2D | 1 | -0.05 | 0.96 | 1 | 1 | 1 |
| place | 2D | 2 | -0.37 | 1.32 | 1 | 1 | 0.8 |
| place | Walk | 1 | -0.38 | 1.15 | 1 | 1 | 0.64 |
| place | Walk | 2 | -0.06 | 1.09 | 1 | 1 | 0.8 |
| place | Controller | 1 | -0.63 | 1.4 | 1 | 1 | 1.08 |
| place | Controller | 2 | -0.3 | 1.13 | 1 | 1 | 0.88 |
| response | 2D | 1 | -0.68 | 1.49 | 1 | 1 | 1.28 |
| response | 2D | 2 | -0.4 | 1.51 | 1 | 1 | 1.12 |
| response | Walk | 1 | -0.91 | 1.59 | 1 | 1 | 1.16 |
| response | Walk | 2 | -0.24 | 1.44 | 1 | 1 | 1.4 |
| response | Controller | 1 | -0.67 | 1.57 | 1 | 1 | 1.04 |
| response | Controller | 2 | -0.06 | 1.03 | 1 | 1 | 0.96 |

Results of the models of regression of the number of explicit choices in strategy based on data from training trials only. For each strategy, VR modality, and session number, the model's R2, RMSE, R2 P-value, RMSE P-value, and mean number of choices in strategy are presented (maximum is 3). Models with significant results greater than chance are marked in bold.

**Table A.5.10:** Regression of the number of explicit choices in strategy based on all trials in sample 2.

| Strategy | Condition | Session | R2 | RMSE | P-value R2 | P-value RMSE | Mean #choices |
|---|---|---|---|---|---|---|---|
| cue | 2D | 1 | -0.2 | 1.27 | 1 | 1 | 1.64 |
| **cue** | **2D** | **2** | **0.08** | **1.2** | **0.04** | **0.04** | **1.84** |
| cue | Walk | 1 | -0.47 | 1.21 | 1 | 1 | 1.96 |
| cue | Walk | 2 | -0.56 | 1.59 | 1 | 1 | 1.44 |
| cue | Controller | 1 | -1.12 | 1.63 | 1 | 1 | 1.84 |
| cue | Controller | 2 | -0.45 | 1.34 | 1 | 1 | 1.96 |
| place | 2D | 1 | 0.04 | 0.92 | 0.069 | 0.069 | 1 |
| place | 2D | 2 | -0.31 | 1.3 | 1 | 1 | 0.8 |
| place | Walk | 1 | -0.46 | 1.18 | 1 | 1 | 0.64 |
| **place** | **Walk** | **2** | **0.37** | **0.84** | **0.01** | **0.01** | **0.8** |
| place | Controller | 1 | 0.13 | 1.02 | 0.089 | 0.089 | 1.08 |
| place | Controller | 2 | -0.35 | 1.15 | 1 | 1 | 0.88 |
| response | 2D | 1 | -0.36 | 1.34 | 1 | 1 | 1.28 |
| response | 2D | 2 | -0.78 | 1.7 | 1 | 1 | 1.12 |
| response | Walk | 1 | -0.1 | 1.21 | 1 | 1 | 1.16 |
| response | Walk | 2 | -0.4 | 1.53 | 1 | 1 | 1.4 |
| response | Controller | 1 | -0.78 | 1.62 | 1 | 1 | 1.04 |
| response | Controller | 2 | -0.47 | 1.21 | 1 | 1 | 0.96 |

Results of the models of regression of the number of explicit choices in strategy based on all trials' data. For each strategy, VR modality, and session number, the model's R2, RMSE, R2 P-value, RMSE P-value, and mean number of choices in strategy are presented (maximum is 3). Models with significant results greater than chance are marked in bold.

#### *A.6 Spatial abilities measures*

**Table A.6.1:** Correlation between choices of cue strategy and MRT

| | rank(n_choices_Cue) | | |
|---|---|---|---|
| *Predictors* | *Estimates* | *CI* | *p* |
| (Intercept) | 38.62 | 21.11 – 56.14 | **<0.001** |
| rank(MRT) | -0.09 | -0.51 – 0.33 | 0.680 |
| Condition [2D] | -1.09 | -27.09 – 24.92 | 0.934 |
| Condition [Controller] | 4.41 | -20.01 – 28.84 | 0.720 |
| rank(MRT) x Condition [2D] | 0.08 | -0.51 – 0.66 | 0.792 |
| rank(MRT) x Condition [Controller] | 0.04 | -0.54 – 0.63 | 0.887 |
| Observations | 75 | | |
| $R^2$ / $R^2$ adjusted | 0.017 / -0.054 | | |

Results of the linear model on ranks for the mean number of explicit choices in cue strategy as a function of MRT score and VR modality.

**Table A.6.2:** Correlation between choices of response strategy and MRT

| | rank(n_choices_Response) | | |
|---|---|---|---|
| *Predictors* | *Estimates* | *CI* | *p* |
| (Intercept) | 43.35 | 25.87 – 60.84 | **<0.001** |
| rank(MRT) | -0.09 | -0.51 – 0.34 | 0.686 |
| Condition [2D] | -5.08 | -31.05 – 20.89 | 0.697 |
| Condition [Controller] | -6.11 | -30.50 – 18.28 | 0.619 |
| rank(MRT) x Condition [2D] | 0.11 | -0.48 – 0.69 | 0.718 |
| rank(MRT) x Condition [Controller] | 0.01 | -0.57 – 0.60 | 0.970 |
| Observations | 75 | | |
| $R^2$ / $R^2$ adjusted | 0.018 / -0.053 | | |

Results of the linear model on ranks for the mean number of explicit choices in response strategy as a function of MRT score and VR modality.

**Table A.6.3:** Correlation between choices of place strategy and MRT

| | rank(n_choices_Place) | | |
|---|---|---|---|
| *Predictors* | *Estimates* | *CI* | *p* |
| (Intercept) | 37.26 | 20.37 – 54.14 | **<0.001** |
| rank(MRT) | -0.08 | -0.49 – 0.32 | 0.681 |
| Condition [2D] | 4.91 | -20.17 – 29.98 | 0.698 |
| Condition [Controller] | 17.10 | -6.45 – 40.65 | 0.152 |
| rank(MRT) x Condition [2D] | -0.00 | -0.57 – 0.56 | 0.993 |
| rank(MRT) x Condition [Controller] | -0.28 | -0.85 – 0.28 | 0.323 |
| Observations | 75 | | |
| $R^2$ / $R^2$ adjusted | 0.070 / 0.003 | | |

Results of the linear model on ranks for the mean number of explicit choices in place strategy as a function of MRT score and VR modality.

**Table A.6.4:** Correlation between choices of cue strategy SOT

| | rank(n_choices_Cue) | | |
|---|---|---|---|
| *Predictors* | *Estimates* | *CI* | *p* |
| (Intercept) | 42.88 | 23.81 – 61.94 | **<0.001** |
| rank(SOT) | -0.16 | -0.54 – 0.21 | 0.387 |
| Condition [2D] | -6.94 | -31.81 – 17.93 | 0.580 |
| Condition [Controller] | 5.75 | -21.45 – 32.95 | 0.674 |
| rank(SOT) x Condition [2D] | 0.20 | -0.35 – 0.75 | 0.473 |
| rank(SOT) x Condition [Controller] | -0.04 | -0.65 – 0.58 | 0.898 |
| Observations | 75 | | |
| $R^2$ / $R^2$ adjusted | 0.034 / -0.036 | | |

Results of the linear model on ranks for the mean number of explicit choices in cue strategy as a function of SOT score and VR modality.

**Table A.6.5:** Correlation between choices of response strategy and SOT

| | **rank(n_choices_Response)** | | |
|---|---|---|---|
| *Predictors* | *Estimates* | *CI* | *p* |
| (Intercept) | 25.21 | 6.55 – 43.87 | **0.009** |
| rank(SOT) | 0.33 | -0.03 – 0.70 | 0.074 |
| Condition [2D] | 13.56 | -10.78 – 37.89 | 0.270 |
| Condition [Controller] | 0.15 | -26.47 – 26.76 | 0.991 |
| rank(SOT) x Condition [2D] | -0.32 | -0.86 – 0.22 | 0.237 |
| rank(SOT) x Condition [Controller] | -0.07 | -0.68 – 0.53 | 0.806 |
| Observations | 75 | | |
| $R^2$ / $R^2$ adjusted | 0.073 / 0.006 | | |

Results of the linear model on ranks for the mean number of explicit choices in response strategy as a function of SOT score and VR modality.

**Table A.6.6:** Correlation between choices of place strategy and SOT

| | **rank(n_choices_Place)** | | |
|---|---|---|---|
| *Predictors* | *Estimates* | *CI* | *p* |
| (Intercept) | 21.78 | 3.73 – 39.84 | **0.019** |
| rank(SOT) | 0.28 | -0.08 – 0.63 | 0.126 |
| Condition [2D] | 19.62 | -3.94 – 43.17 | 0.101 |
| Condition [Controller] | 0.67 | -25.10 – 26.43 | 0.959 |
| rank(SOT) x Condition [2D] | -0.36 | -0.88 – 0.16 | 0.170 |
| rank(SOT) x Condition [Controller] | 0.25 | -0.33 – 0.83 | 0.390 |
| Observations | 75 | | |
| $R^2$ / $R^2$ adjusted | 0.118 / 0.055 | | |

Results of the linear model on ranks for the mean number of explicit choices in response strategy as a function of SOT score and VR modality.

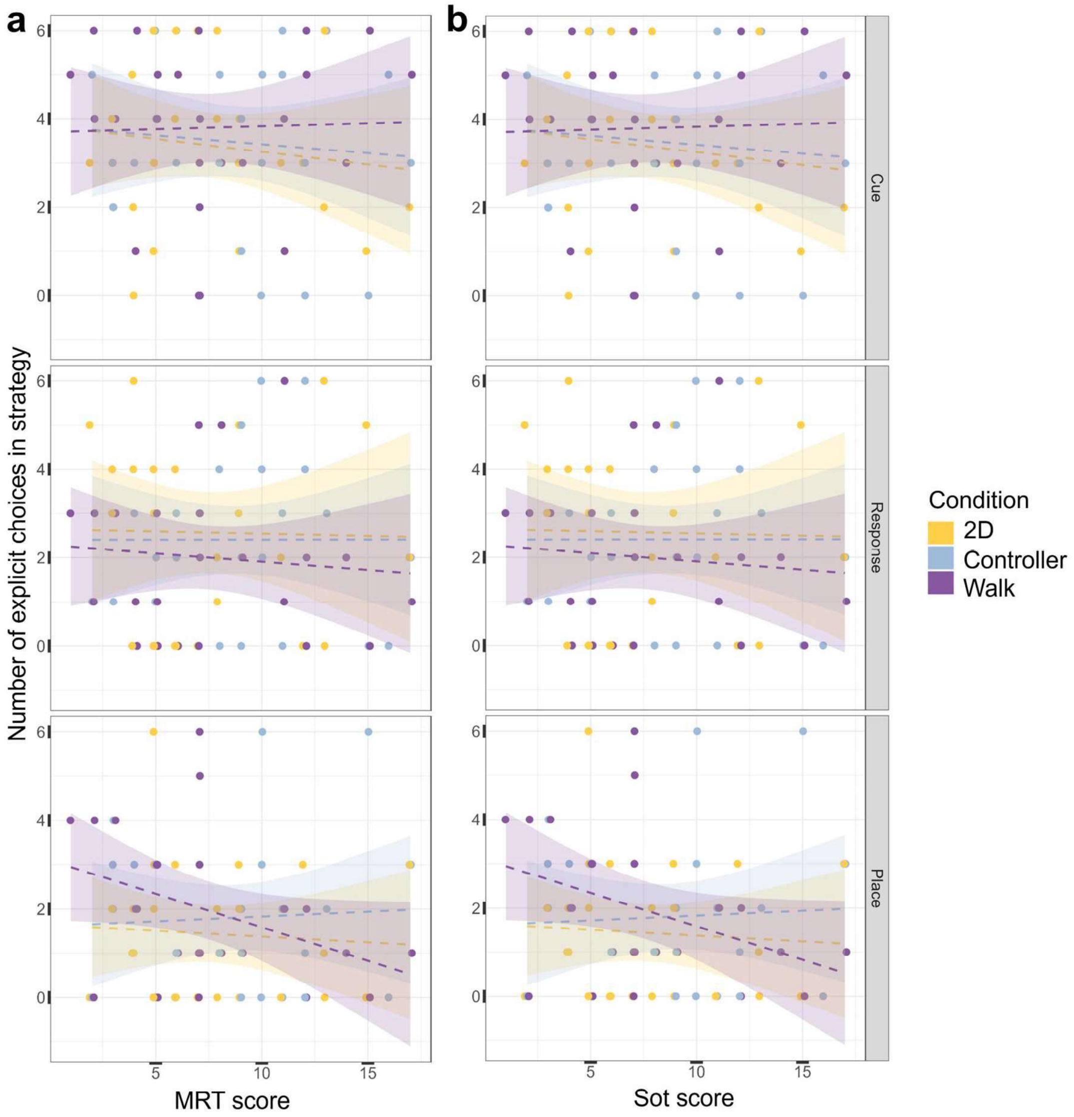


**Fig. A.6.1: Spatial abilities measures.** Scatter plots of the mean across sessions of the number of explicit choices in strategy in each strategy test probe trial, as a function of the score in spatial abilities tests. Individual data points are shown as dots slightly jittered around the x-axis for display purposes. The dashed lines represent the linear models fit for each strategy and each VR modality (fitted lines are surrounded by polygons of the 95% confidence interval). **a:** MRT results. **b:** SOT results.
